# High-throughput Activity Reprogramming of Proteases (HARP)

**DOI:** 10.1101/2025.03.27.640893

**Authors:** Samantha G Martinusen, Ethan W Slaton, Seyednima Ajayebi, Marian A Pulgar, Cassidy F Simas, Sage E Nelson, Amit Dutta, Julia T Besu, Steven Bruner, Carl A Denard

## Abstract

Developing potent and selective protease inhibitors remains a grueling, iterative, and often unsuccessful endeavor. Although macromolecular inhibitors can achieve single-enzyme specificity, platforms used for macromolecular inhibitor discovery are optimized for high-affinity binders, requiring extensive downstream biochemical characterization to isolate rare inhibitors. Here, we developed the High-throughput Activity Reprogramming of Proteases (HARP) platform, HARP is a yeast-based functional screen that isolates protease-inhibitory macromolecules from large libraries by coupling their inhibition of endoplasmic reticulum-resident proteases to a selectable phenotype on the cell surface. Endowed with high dynamic range and resolution, HARP enabled the isolation of low-nanomolar-range inhibitory nanobodies against tobacco etch virus protease and human kallikrein 6, including a rare 7.6 nM *K_I_* TEVp uncompetitive inhibitor. Structural modeling and deep sequencing all provide insights into the molecular determinants of inhibitors and reinforce HARP’s foundational findings. Overall, HARP is a premier platform for discovering modulatory macromolecules from various synthetic scaffolds against enzyme targets.

**Graphical Abstract:** *Workflow of HARP:* A yeast-based reporter of the interaction between a protease, its canonical substrate, and a modulator library within the yeast cell. Quantifying this interaction occurs by fluorescently labeling the displayed substrate cassettes on the surface of the cells, where the desired function (correlating with phenotype) can be selected using fluorescent-activated cell sorting (FACS). Isolated populations are sequenced and purified in preparation for secondary characterization to determine modulator effects and interaction strengths between the modulator and the protease target.

## Introduction

Proteases catalyze the hydrolysis of peptide bonds in proteins. Through this reaction, they regulate critical biochemical pathways, thereby maintaining cellular proteostasis^1–6^. Unsurprisingly, protease dysregulation is a hallmark of various diseases, making them important therapeutic targets in cancer, autoimmune disorders, cardiovascular diseases, neuropathic pain, and infectious diseases^7–13^. Although proteases comprise ∼2% of the proteome and represent 5-10% of all drug targets (2010 estimate)^7^, only a small fraction of therapeutically relevant proteases (i.e., viral proteases) have been successfully targeted, resulting in only ∼20 FDA-approved protease inhibitors^10^. These inhibitors are primarily small molecules and peptidomimetics that engage their target’s active site. Unfortunately, these molecules often exhibit poor selectivity and undesirable off-target effects because they cannot differentiate between similar active-site topologies of related proteases^14, 15^. These challenges are exemplified by the failed clinical trials of matrix metalloprotease inhibitors^16^ for cancer therapy and BACE1 inhibitors to treat Alzheimer’s Disease^17^. While targeting allosteric sites confers higher selectivity, discovering protease allosteric sites and small-molecule allosteric modulators is hampered by an incomplete understanding of the general principles underpinning allosteric and distal regulation^18^. Biased high-throughput screening strategies, residue tethering, activity-based profiling^19^, exosite-targeted screens and computational methods aim to close this gap. Still, few of these techniques can be applied across all protease families^20^. Taken together, discovery campaigns for selective protease inhibitors remain slow, arduous, and arguably over-reliant on serendipity^18, 21^.

Macromolecular proteinaceous inhibitors can address selectivity and pharmacological limitations faced by small molecules^22, 23^. Particularly, natural protein scaffolds, which include antibodies and their fragments (nanobodies, ScFvs, Fabs), ankyrin repeats, macrocyclic peptides, and cysteine knots, can be screened to find exquisitely selective and potent protease inhibitors^15, 24^. By burying more surface area upon binding, macromolecules gain more potency and specificity than small molecules^25^. Furthermore, whereas natural endogenous inhibitors usually do not take advantage of allosteric movements of proteins, protease-inhibitory macromolecules derived from natural scaffolds can attain allosteric regulation by binding to less conserved sequences on the enzyme target and targeting specific conformations^26^. These interactions can elucidate mechanisms that have not been seen in nature and might not have been discovered using traditional high-throughput screening techniques.

There is a critical unmet need to accelerate the discovery and engineering of protease-inhibitory macromolecules. Unfortunately, current selection/screening methods, including hybridoma technology^27, 28^, phage panning^29–33^, cell^34–36^ and bead^37–39^ surface display coupled with flow cytometry is primarily based on binding, not inhibition. As a result, extensive downstream characterization is required to isolate inhibitors for large pools of binders. Typically, fewer than 5% of binders isolated through these methods inhibit the targeted protease^40^. Crucially, potentially inhibitory clones with moderate binding affinities are lost during repeated cycles of competitive binding^41^. Previously reported *E. coli*-based genetic selections for protease inhibitory Fabs^42^ and ScFvs^43^ have a narrow discovery window (isolates many weak inhibitors with > 1 µM *K_I_*), require optimization for each protease target and may be limited by the activity/toxicity of human proteases in bacteria.

To address these challenges, we present HARP (High-throughput Activity Reprogramming of Proteases), a yeast-based strategy that couples the modulation of endoplasmic reticulum (ER) targeted proteases to selectable phenotypes on the yeast cell surface. In this fashion, HARP enables one to discover protease-inhibitory macromolecules from large combinatorial protein binder libraries via fluorescence-activated cell sorting (FACS). Using an MMP8/MMP8 inhibitory nanobody (Nb) pair, we establish key principles of the HARP platform to isolate protease inhibitory Nbs. We then showcase HARP’s potential by isolating potent orthosteric Nbs and an allosteric inhibitor Nb (*K_I_* ranging from 7-50 nM) against tobacco etch virus protease (TEVp) from a synthetic Nb library for the first time. Deep sequencing of the TEVp Nb library reveals that inhibition phenotypes from yeast correlate linearly with *in vitro* performance, establishing HARP as a robust and quantitative inhibitor discovery platform. ESM-2 embedding followed by supervised dimensionality reduction of TEVp inhibitory Nb sequences shows distinct clustering and a significant demarcation between inhibitory and non-inhibitory sequences. Lastly, we showcase HARP’s ability to isolate a highly selective and potent kallikrein 6 inhibitory Nb. Considering that >30% of proteases are secreted extracellularly or membrane-bound^44^, HARP is a premier platform to discover and engineer protease modulatory macromolecules that could become valuable reagents and therapeutic agents. With minor modifications, HARP should be straightforwardly expanded to various synthetic scaffolds and enzyme targets beyond proteases.

## Results and Discussion

### HARP capitalizes on yeast ER sequestration and screening concepts for protease modulator discovery

ER sequestration and screening, empowered by effective transcriptional and post-translational strategies, has proven instrumental for the high-throughput interrogation and engineering of post-translational modifications enzyme activities, including proteases, kinases, and acetyltransferases^45–48^. We reasoned that one could leverage this approach to isolate genetically encoded macromolecules that modulate the activity of ER-sequestered proteases, for example, through inhibition. Thus, we set out to build the HARP system, a high-throughput functional screen for protease modulatory macromolecule discovery and engineering. HARP consists of three transcriptional units: a protease, a substrate cassette, and a protein-based modulator (**Figure 1A**). The protease contains a 5’ ER targeting sequence and a 3’ ER retention signal (ERS), functionally immobilizing it in the yeast ER. Similarly, the protein-based protease modulator is targeted to the ER and may contain an ERS to increase its ER concentration and interaction time with the protease. Finally, the substrate cassette, which reports the activity of the ER-resident protease, is an AGA2 fusion polypeptide that transits through the ER on its way to the cell surface. Substrate ER residence time can be manipulated by appending an ERS at the 3’ end of the AGA2-substrate cassette. The AGA2-substrate polypeptide fusion contains unique epitope tags flanking the substrate sequence that can be stained with fluorescently labeled antibodies once on the cell surface. Therefore, this substrate cassette serves as an extracellular reporter of enzyme activities. During the operation of the HARP platform with a protease, three phenotypes are possible: (1) no surface display, (2) a cleaved substrate (resulting from the protease remaining active), and (3) an intact substrate (resulting from an inhibited protease or a substrate-targeted inhibitor) (**Figure 1B**). Cells displaying the desired phenotype (in this case, an intact substrate) can be selected and enriched using FACS.

**Figure 1.**
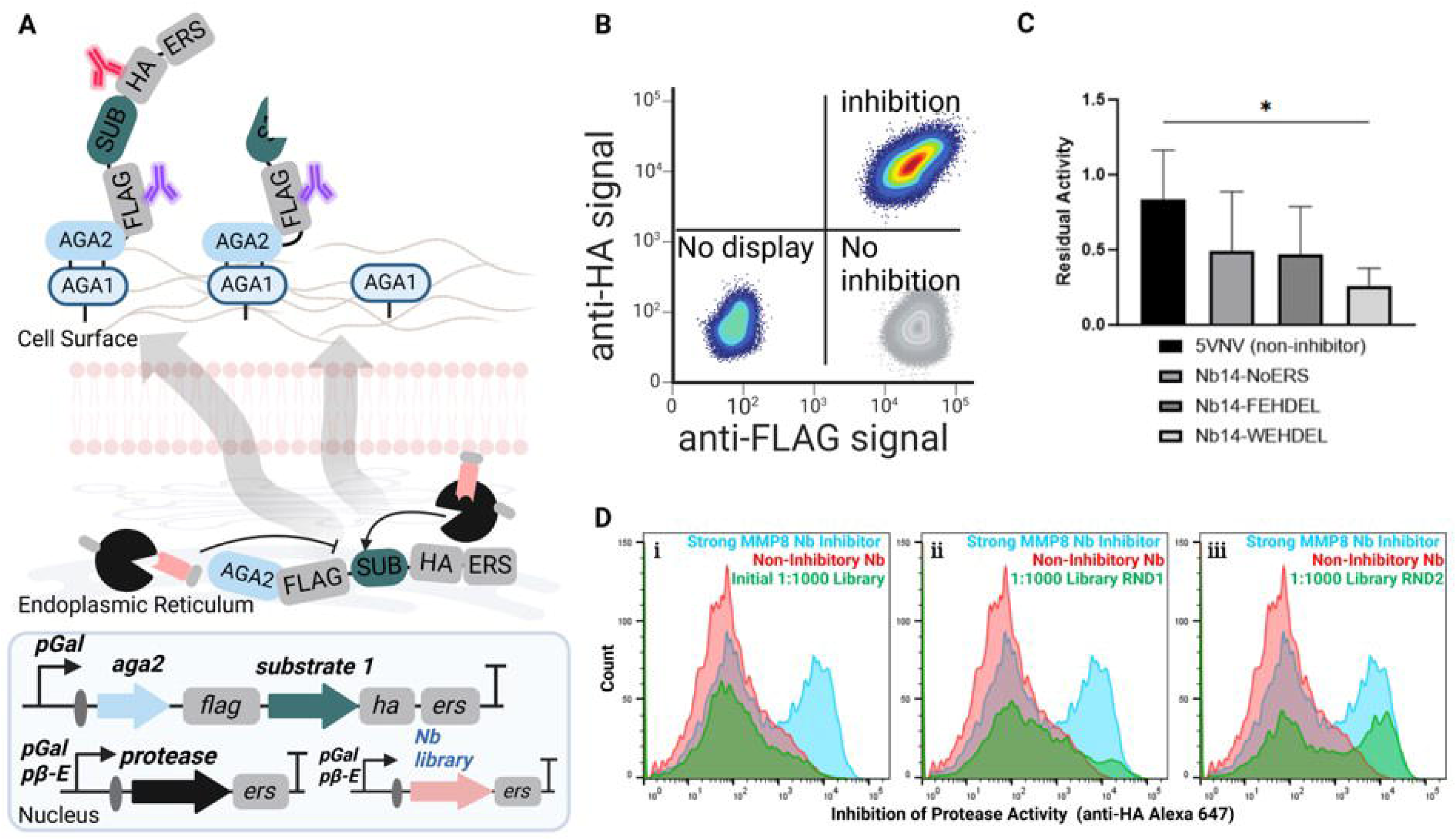
High-throughput Activity screen for the functional Reprogramming of Proteases (HARP). (A) Schematic representation of the HARP workflow in yeast. (B) Using HARP to isolate inhibitors, Yeast Surface Display (YSD) phenotypes are possible. (C) Quantification of observed inhibition of MMP8 as a function of Nb14 ER retention. (D) MMP8 Nanobody (Nb) Inhibition Mock Sort Histograms illustrating the enrichment of protease inhibition (green, anti-HA Alexa 647 signal) going from left to right. Controls included for signal comparison: strong MMP8 Nb inhibitor (red) and non-binder, non-inhibitory Nb (light blue).

To demonstrate the HARP system, we conducted a proof-of-concept experiment using a published MMP8 inhibitory nanobody (Nb), Nb14. Nb14, discovered from a phage display Nb library, has a dissociation constant (*K_D_*) of 1.33 ± 0.14 nM but a poor half maximal inhibitory concentration (IC_50_) of 4.36 ± 0.24 µM^41^. This highlights a common issue where strong enzyme binders selected through display methods often function as weak inhibitors^42, 49–51^. To evaluate Nb14-mediated MMP8 inhibition in yeast, we co-expressed Nb14 and MMP8 in the yeast ER with an ER-transiting AGA2-MMP8 substrate fusion. MMP8 and its substrate contained a C-terminal WEHDEL^52^ ERS to ensure high MMP8 activity in the ER. A β-estradiol-inducible promoter controlled MMP8 expression to avoid cytotoxicity observed with *pGAL1*-driven expression. As a control, Nb14 was replaced with the non-binding, non-inhibitory Nb 5vnv^53^. Results showed that 1) 5vnv showed no inhibition of MMP8 activity, and 2) Nb14’s inhibitory potency depended on its C-terminal ERS strength. The WEHDEL ERS inhibited MMP8 by 74%, while the FEHDEL ERS reduced MMP8 activity by 53% (**Figure 1C, Supplemental Figures 1 and 2**). When the ERS was completely removed, MMP8 activity was comparable (51%) to that observed when using the FEHDEL.

**Figure 2.**
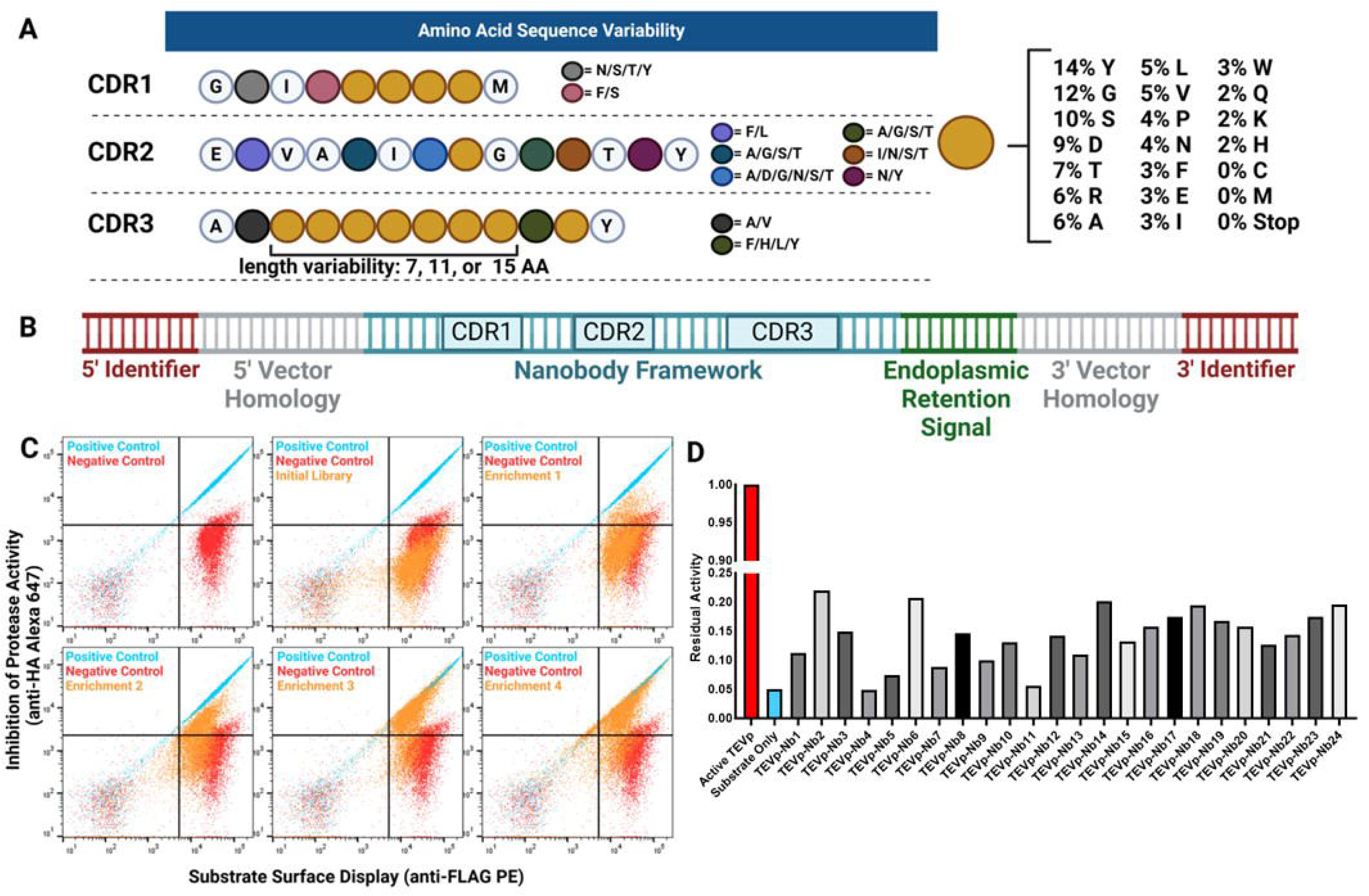
Synthesis of a large Nb library and isolation of inhibitory Nbs against TEVp. (A) Amino acid and length (CDR3) variation present within the NbLibrary_NE_TEVp. (B) Schematic of NbLibrary_NE_TEVp construction prepared for homologous directed recombination (HDR). Complementary determining region (CDR) locations are approximate, as well as the endoplasmic reticulum retention signal (if applicable). (C) Flow plots for each FACS round of the NbLibrary_NE_TEVp sort with light blue populations corresponding to the positive control (TEVp substrate integrated), red populations corresponding to the negative control (TEVp active proteases integrated), and orange populations corresponding to the NbLibrary_NE_TEVp library populations after each FACS round. (D) Representation of the anti-FLAG/anti-HA ratio (normalized to active TEVp protease) of the selected constructs isolated from NbLibrary_NE_TEVp. Dot plots are represented here on a log scale.

Next, we performed a mock sort to evaluate HARP’s ability to isolate inhibitory proteins from a large pool of non-inhibitors. We mixed cells expressing Nb14/MMP8/MMP8 substrate with cells expressing 5vnv/MMP8/MMP8 substrate at a 1:1000 ratio. All constructs included a strong C-terminal WEHDEL ERS. After two rounds of sorting for an inhibitory phenotype, indicated by high anti-FLAG and high anti-HA signals, the library was fully enriched for Nb14-expressing cells, as verified by Sanger sequencing of DNA isolated from ten isolated clones **(Supplemental Figures 3 and 4**). This result demonstrated HARP’s capability to effectively identify inhibitory proteins from a large, diverse population **(Supplemental Figure 5)**.

### High-throughput discovery of TEVp inhibitory NbsHigh-throughput discovery of TEVp inhibitory Nbs

With HARP’s screening capability established, we aimed to isolate protease inhibitory Nbs from a large synthetic library, choosing TEVp as a test case. We selected Nbs as the inhibitory scaffold because their longer complementary determining regions (CDRs) (average length of 15 amino acids) can adopt convex paratope structures that allow them to penetrate enzyme active sites^54^. Nb CDRs can engage distal pockets or surface loops on their targets that could confer allostery. Lastly, since Nbs are relatively small (15 kDa) and can be easily expressed in the periplasm of *E. coli*, they fit within our streamlined *in vitro* characterization pipeline. Two TEVp inhibitors have been reported to date. Both are cyclic peptides chemically modified with a covalent warhead, with IC_50_s of 700 nM^30^ and 80 nM^55^. Note that the covalent warhead accounts for the majority of their inhibitory potency. Rather than chemically modified macromolecules, fully recombinant protease inhibitory Nbs are easier to scale and translate *in vivo*. Moreover, inhibitory Nbs against TEVp and related proteases could be used as components of feedback loop designs in protein circuits^56, 57^. More broadly, inhibitory Nbs against proteases of the potyviridae family could provide a novel and specific method to generate virus-resistant engineered plants^56, 57^.

For the inhibitor isolation campaign, we emulated the synthetic camelid Nb library previously reported by Kruse and coworkers^58^. This 1-billion-member library, built on a curated set of highly stable and biochemically well-behaved Nb structures from the PDB, contains CDRs varying in amino acid composition and CDR3 length (10,14, and 18 amino acids) (**Figure 2A-B**). The 10 amino acid CDR3 comprises 7 completely randomized positions (herein called CDR-7), the 14 aa CDR3 has 11 randomized positions (CDR-11), and the 18 aa CDR3 has 15 randomized positions (CDR-15). In an initial attempt to discover TEVp inhibitory Nbs, an FEHDEL ERS was appended to the C-terminus of the Nb library and transformed using homologous directed recombination (HDR) into a yeast strain containing a chromosomally integrated TEVp and TEVp substrate, pGAL1-AGA2-FLAG-ENLYFQS-HA-FEHDEL. Unfortunately, Nbs isolated from this configuration showed little to no TEVp inhibition *in vitro*. Moreover, the best candidate failed to retain their inhibitory phenotype in yeast when retested without an ERS **(Supplemental Figures 6 and 7**), suggesting that 1) artificially enhancing Nb: protease contact time in the ER leads to false positives and that 2) an Nb with an ERS can outcompete the protease for binding to ERS receptors^52^.

These results highlighted two key considerations for successfully isolating potent inhibitors. First, high enzyme activity (reflected by an anti-FLAG/anti-HA signal normalized ratio of 1) (**Figure 1C**) creates a sufficiently large inhibitor selection window. Using strong promoters to drive protease and substrate expression and appending strong ERSs to both cassettes satisfy this criterion. Second, removing the ERS from potential inhibitor gene cassettes can eliminate weak inhibitors during screening. The steady-state ER concentration of an inhibitor population without an ERS is lower than that of the protease, as most potential modulators translocate to the cytoplasm. Therefore, while weak and non-binders are “washed out” of the ER, moderate to strong binders remain. However, being a functional screen, HARP only isolates binders that inhibit enzyme activity via FACS. Thus, setting a high protease activity (strong ERS) and reduced inhibitor concentration (No ERS) in HARP applies selective pressure to enrich for potent inhibitors. Therefore, we reformatted the dsDNA Nb library, devoid of an ERS, to generate NbLib_TEVp-NE library **(Supplementary Table 1)**, comprising approximately 2 × 10L variants.

To enrich for TEVp inhibitors, cells harboring the NbLib_TEVp-NE (10L cells) library were induced in galactose media and stained with fluorescently labeled antibodies—anti-HA Alexa 647 (BioLegend, cat# 682404) and anti-FLAG PE (BioLegend, cat# 637309) - and subjected to FACS. In every round, two control strains were included to guide the gating strategy: a substrate display-only control lacking TEVp (No TEVp) and the TEVp reporter strain **(Supplemental Table 1)**. The goal was to isolate Nb-harboring cells phenotypically resembling those of the substrate display-only control, indicative of potent TEVp inhibition. Therefore, gating stringencies were increased from round to round **(Supplemental Figures 8 and 9**). After four rounds of FACS, during which the NbLib_TEVp-NE library became progressively enriched for cells exhibiting an inhibitory phenotype (populating in the upper-right quadrant), beginning with ∼1% (Initial Library) and enriching to ∼60% (ENR4) of the total population (**Figure 2C**). At the end of the Nb discovery campaign (ENR4), 24 individual clones were rescreened in yeast to evaluate the diversity of inhibitory phenotypes (**Figure 2D**). All clones harbored Nbs that inhibited TEVp by ≥ 80% (≤ 20% residual activity). Sanger sequencing revealed 14 unique sequences **(Supplemental Table 2)**, Repeat clones had similar in-yeast performances, showcasing experimental reproducibility.

### In vitro characterization of TEVp inhibitory Nbs

Following single clone screening, we characterized four Nb clones with varying percent residual activity (PRA) in yeast: TEVp-Nb4 (4.8%), TEVp-Nb5 (7.4%), TEVp-Nb7 (8.9%), and TEVp-Nb9 (10%) (**Figure 2D**). The aim was to determine whether inhibitory phenotypes observed in yeast correlated with *in vitro* potencies. Nb IC_50_ and inhibition constants (*K_I_*) were determined using an Abz-TENLYFQSGK(Dnp) FRET peptide and are summarized in Table 1. Excitingly, TEVp-Nb4, which had the lowest PRA in yeast, was the strongest inhibitor *in vitro*, with an IC_50_ and *K_I_* of 29 and 8 nM, respectively. This makes TEVp-Nb4 the most potent TEVp inhibitor to date. Furthermore, TEVp-Nb5 (PRA: 7.4%, IC_50_: 201 nM, *K_I_*: 53 nM), TEVp-Nb7 (PRA, 8.9%, IC_50_: 420 nM, *K_I_*: 110 nM), and TEVP-Nb9 (PRA: 10%, IC_50_:178 nM, *K_I_*: 47 nM) all showed significant inhibitory potencies (**Table 1, Supplemental Figure 10)**.

**Table 1.**
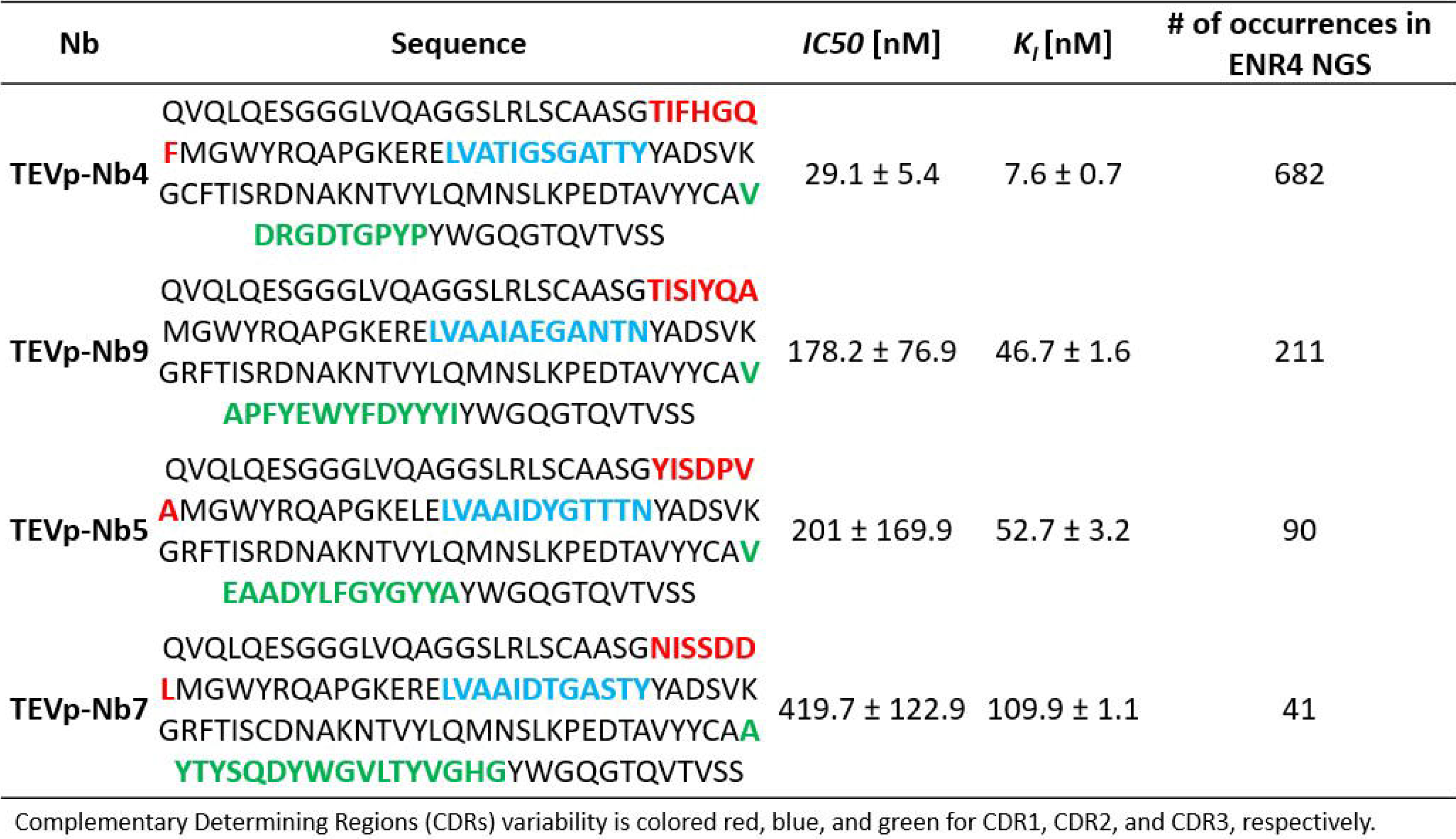
Inhibition potency characterization of isolated TEVp-Nbs.

These results suggested Nb *in vitro* potencies generally correlated with their in-yeast performance. On the one hand, the strongest inhibitor, TEVp-Nb4, showed the highest in-yeast performance. On the other hand, TEVp-Nb7, which showed a lower in-yeast performance, had a lower potency *in vitro*. We hypothesized that HARP could provide a robust and quantitative measurement of protease inhibition, meaning that Nbs with the highest read counts at the end of the campaign would likely be more inhibitory *in vitro*. To test this hypothesis, we sequenced the post-round two libraries (26,604 unique sequences) and the final-round library (16,006 unique sequences) **(Dataset 1)** and determined the number of occurrences for all unique sequences in the final round. Above all, the four Nbs randomly selected for *in vitro* characterization ranked among the top 50 most abundant sequences **(Supplemental Table 3)**. Pleasingly, their *K_I_s* corresponded with their positions in the NGS dataset: TEVp-Nb4 (*K_I_*= 7.6 nM, 3^rd^ most abundant, 682 occurrences), TEVp-Nb9 (*K_I_* = 46.7 nM, 13^th^ position, 211 occurrences), TEVp-Nb5 (*K_I_*= 52.7 nM, 23^rd^ position, 90 occurrences), and TEVp-Nb7 (*K_I_*= 109.9 nM, 48^th^ position, 41 occurrences) (**Table 1, Supplemental Table 3)**. Furthermore, the positive correlation between in-yeast performance, read count and, by extension, *in vitro* potency generally carried over to the unique clones verified by Sanger Sequencing **(Supplemental Table 2)**, as 13/14 aforementioned unique clones were all found within the top 80 most abundant sequences. The remarkable resolution provided by HARP further cements its potential for protease inhibitor discovery to the degree that derived sequence-inhibition potency correlations will be vital for developing machine learning algorithms for function prediction and inhibitory Nb design.

### Biochemical characterization and structural modeling of TEVp inhibitory Nbs

To investigate whether TEVp inhibitory Nbs are orthosteric or allosteric inhibitors, we performed separate experiments where we measured the binding affinities of TEVp-Nbs 4, 5, 7, and 9 to deactivated TEVp (dTEVp) and to the dTEVp-substrate complex by Biolayer Interferometry (BLI) (**Table 2, Supplemental Figure 11)**. Deactivated TEVp is a catalytic cysteine-to-alanine mutant (C151A) that retains substrate binding capability and whose X-ray structure overlaps exactly with that of TEVp^59^, allowing us to perform assays with a substrate-bound enzyme. We chose dTEVp rather than TEVp for another reason. TEVp, both commercial and in-house grade, is purified as an MBP-TEV cleavage site-His tag-TEVp fusion during which TEVp cleaves itself from the MBP solubilizing protein, allowing it to be purified by affinity chromatography. Interestingly, the residual product, MBP-ENLYFQ or ENLYFQ, can remain in the enzyme active site^59^. We reasoned that this issue could inadvertently yield misleading results in binding assays. In fact, Nb binding assays using TEVp, rather than dTEVp, showed moderate to poor Nb binding **(Supplemental Table 4, Supplemental Figure 12)**.

**Table 2.** Binding characterization of isolated TEVp-Nbs.

| Nb | $K_D$ [nM]<br>dTEVp | $K_D$ [nM]<br>dTEVp + Sub | $K_D$ [nM]<br>TVMVp |
| --- | --- | --- | --- |
| TEVp-Nb4 | 2050 ± 130 | 66 ± 13.5 | N/D |
| TEVp-Nb9 | 18 ± 2.2 | N/D | N/D |
| TEVp-Nb5 | 163 ± 21.1 | N/D | N/D |
| TEVp-Nb7 | 17 ± 2.6 | N/D | N/D |

N/D: Not Detectable

TEVp-Nbs 5, 7, and 9 bind to dTEVp with *K_D_s* of 18 nM, 163 nM, and 17 nM, respectively - values in line with their *K_I_s* - qualifying them as monovalent binders (**Table 2**). However, they do not bind to the dTEV-substrate complex, supporting a competitive inhibition mechanism. In lieu of Nb-TEVp X-ray co-crystal structures (not yet available), we generated Alphafold3 models of Nbs 5, 7, and 9 with dTEVp **(Supplemental Figure 13)**. These models had acceptable predicted template modeling (pTM) scores (>0.5), indicating reasonable overall structural accuracy^60^. However, their low interface-predicted template modeling (ipTM) values (<0.5) suggest low confidence in the relative positioning of interacting residues^60^. Nonetheless, Nb5, 7, and 9 models bound to dTEVp all corroborate active-site inhibition (**Supplemental Figure 13**). Overall, the long CDR3s (variable lengths ≥ 11) of Nbs 5, 7, and 9 engage TEVp’s active site in a non-substrate-like manner, significantly disrupting the positioning and, in some cases, the secondary structures of the highly flexible C-terminus of TEVp (Gln195-Pro226), which includes S2-S5 pocket residues. Additionally, parts of the CDR3s of Nb7 and Nb9 may form parallel β-sheets with those of TEVp’s S2 subsite, likely contributing to stabilizing the large structural distortions.

### Nb4 is an uncompetitive inhibitor that engages a positively charged pocket outside TEVp’s S2 subsite

While Nbs 5, 7, and 9 were identified as active-site inhibitors, BLI results suggest that TEV-Nb4 functions as an uncompetitive inhibitor. TEVp-Nb4 has a *K_I_* of 6.7 nM but binds weakly to dTEVp alone (∼2 µM). Yet, its binding affinity increases significantly to 66 nM in the presence of the dTEVp-substrate complex. AlphaFold3 models of TEVp-Nb4 bound to the dTEVp-substrate complex yielded reliable pTM (0.62) and ipTM (0.63) values (**Figure 3**), providing preliminary insights into this novel binding mechanism. As illustrated in Figure 3A, the highly charged CDR3 of TEVp-Nb4 (97-VDRGDTGPYP-106) interacts with a positively charged shallow depression on the opposite side of the S2 subsite through an intricate network of hydrogen bonds and hydrophobic interactions (**Figure 3A-B**). Specifically, the CDR3 forms a tight loop anchored by a salt-bridge interaction between Asp101^Nb4^ and Arg99^Nb4^. This presents the β-carbon of Asp101 toward a hydrophobic cleft that includes the indole of Trp211^TEVp^ and sidechains of Val209^TEVp^ and Leu210^TEVp^. On the periphery of the hydrophobic pocket, hydrogen bonding interactions with each amino acid in the loop (Asp98 to Thr102) interact with TEVp including Asp77-Thr102, Lys44–Asp101, Asp100-Asn204 and Asp98-Asn204. Hydrogen bonding between Thr102^Nb4^ and Asp78 ^TEVp^ anchors the CDR3 against one of the pocket’s flexible loops (Leu76-Met82). Meanwhile, the sidechains of Thr102^Nb4^ and Asp98^Nb4^ form hydrogen bonds with Asn204^TEVp^ and Asp77^TEVp^, respectively. Along with hydrogen bonding interactions of the backbone carbonyls of Asp101^Nb4^ and Asp100^Nb4^TEVp, the loop stabilizes the CDR3 against the opposite, lengthy, flexible loop (Ala194-Ser208). Interestingly, interactions mediated by CDR2 and framework residues further contribute to Nb4’s high binding affinity **(Supplemental Figure 14)**. We hypothesize that, while the conformation of the shallow pocket may be engendered by substrate binding, Nb4 binding likely restricts the movement of the flexible loops and S2 subsite, effectively locking the TEVp-substrate complex into a non-productive state. It is intriguing that Nb4 interacts with the S2 subsite of TEVp. Although not reported for TEVp, the S2 subsite of 3C proteases from Human Rhinovirus, Enterovirus A, and SARS-CoV-2 share a similar fold to TEVp. This fold is characterized by hydrophobic residues which can adopt an open or closed conformation^61^, and has been the target for drug discovery due to its importance in substrate recognition^62^. Future molecular dynamics (MD) simulations and X-ray crystal structures will elucidate the molecular mechanisms of Nb4-mediated inhibition.

**Figure 3.**
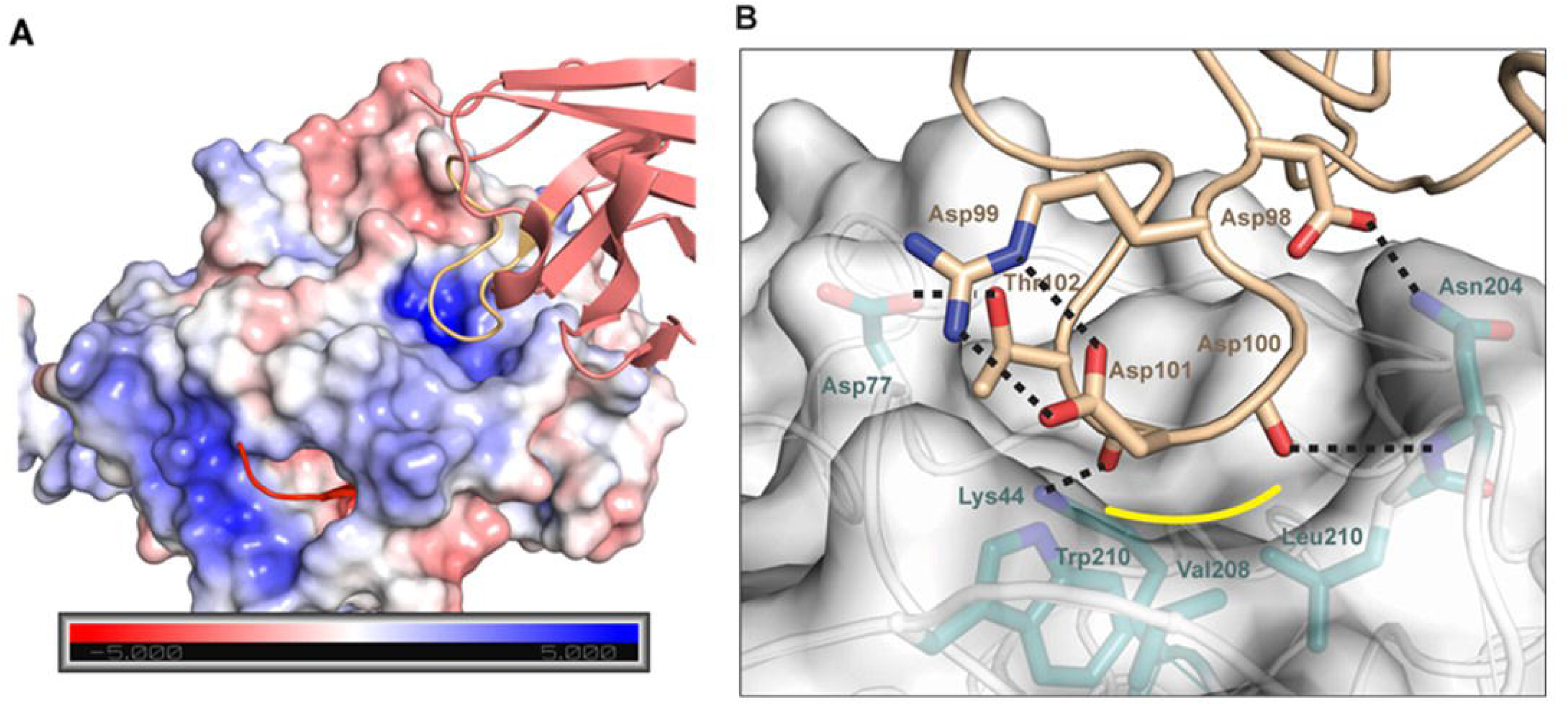
TEVp-Nb4 binding mechanism predicted by AlphaFold3. (A) Electrostatic surface map of dTEVp bound to its peptide substrate (red ribbon) and nanobody Nb4. Nb4 binds to a positively charged surface pocket opposite the S2 subsite of dTEVp. (B) Expanded view of the interaction of the Nb4 CDR3 (tan) loop and the dTEVp (cyan) surface. Residues of the loop (98-102) make specific interactions including hydrogen bonds (dashed lines) along with a hydrophobic interface (yellow).

### Deep sequencing and analysis of TEVp-Nb CDR3s provide insights into the molecular determinants of inhibition

The CDR3 regions of antibodies and nanobodies play a crucial role in determining binding affinity and, in our case, inhibition. Deep sequencing of Nb libraries revealed that while the sorted library contained 34,138 unique Nb sequences, it included only 1,268 distinct CDR3 sequences, with lengths of 10 (32%), 14 (52%), and 18 (14%) amino acids—mirroring the distribution of the starting library (**Figure 4**). Highly abundant CDR3s appeared in Nb sequences with high and low read counts, indicating that CDR3s underwent more stringent selection than CDRs 1 and 2. Additionally, while CDR3s of variable length 11 (CDR-11) yielded more inhibitory sequences, CDR-7 inhibitors produced more potent inhibitors (**Figure 4A**), consistent with our identification of TEV-Nb4 as the most effective inhibitor in single-clone screening.

**Figure 4.**
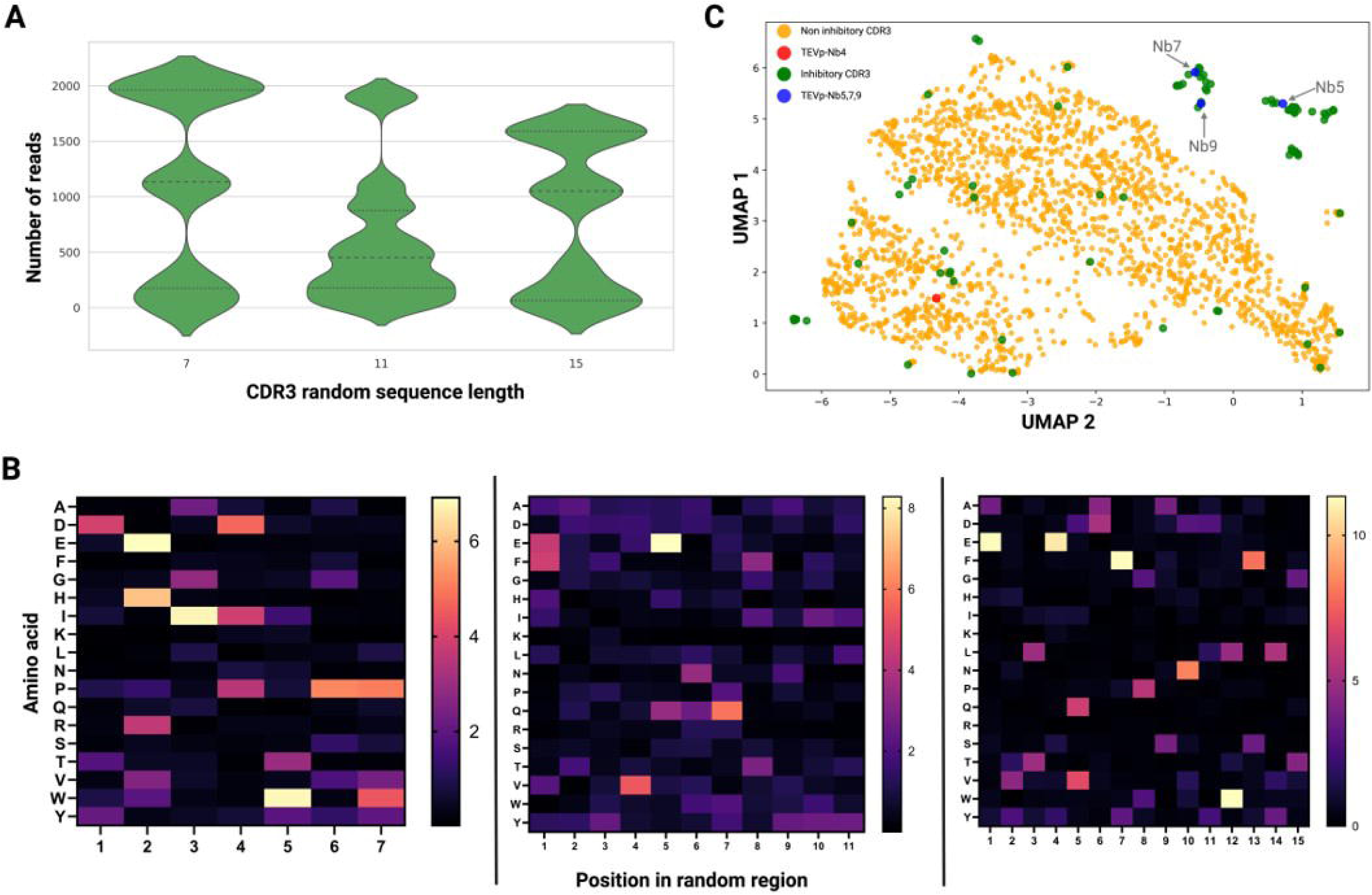
NGS analysis of NbLibrary_NE_TEVp. (A) The distribution of CDR3 sequences based on the number of reads after round 4. (B) UMAP Supervised dimensionality reduction of CDR3 sequences. ESM-2 embeddings show distinct clusters of inhibitory versus non-inhibitory sequences. TEV-Nb4’s CDR3 appears far away from TEV-Nb5, 7, and 9, reinforcing that TEV-Nb4 binds to a different epitope. (C) Heatmap of amino acid frequency enrichment (Round 4 / Naive) for CDR3 sequences.

To identify key sequence features of TEVp Nbs, we analyzed amino acid frequency enrichments at each position along the CDR3 region by comparing the final sorted library to the original 1-billion-member naïve synthesized library. This analysis revealed similarities and stark differences in the amino acid composition of inhibitory CDR3s depending on their variable lengths (**Figure 4B**). CDR-11 and CDR-14 sequences displayed similar patterns, with notable enrichments of Glu at positions 1 (P1) and 5 (P5), and Phe at P7. Co-variance analysis of CDR-11 sequences **(Supplemental Figure 15)** identified a P5-P7 ENQ motif resembling a TEVp substrate fragment **(Dataset, Supplemental Table 5)**. In contrast to CDR-11 and 14, CDR-7 sequences exhibited high variability across all positions in their variable region. Notable enrichments included Asp at P1 and P4, Glu, His, Arg at P2, Ile at P3, and Pro at P4, P6, and P7. Co-variance analysis of highly abundant sequences identified a P3-P7 EIDW inhibitory motif, which is highly enriched in a dominant CDR3 sequence **(Supplemental Figure 15)**. One particularly intriguing observation was the enrichment of prolines in CDR-7 sequences. While prolines are rarely found in Nb-binding interfaces^63, 64^, their presence—especially at the borders of CDR3s^65^—limits CDR conformational flexibility and is known to enhance the kinetics and affinity of immunoglobulin interactions^66^. Lastly, investigating the enrichment of Arg at P2 revealed that its enrichment was primarily due to its presence in the highly abundant TEV-Nb4. The CDR3 sequence of TEV-Nb4 diverges from all other Nb inhibitory sequences isolated, suggesting that TEV-Nb4 is a unique Nb, likely binding to a distinct TEVp epitope.

Owing to the diversity of inhibitory CDR3s and to study differences between inhibitory and non-inhibitory CDR3 sequences, we used Evolutionary Scale Modeling (ESM-2)^67^ a pre-trained language model for proteins and a supervised dimensionality reduction with Uniform Manifold Approximation and Projection (UMAP)^68^ to embed and visualize Nb sequences in 2D space. Pleasingly, most inhibitory Nbs clustered away from non-inhibitory ones (**Figure 4C**). More importantly, TEV-Nb4 is positioned far from TEV-Nb5, TEV-Nb7, and TEV-Nb9, which bind to the active site, further supporting the hypothesis that Nb4 targets a different TEVp epitope.

### HARP can isolate selective kallikrein 6 inhibitory Nbs for the first time

Establishing HARP with hMMP8 demonstrates that eukaryotic proteases active in the yeast ER can be targeted for inhibitory Nb development. To further explore HARP’s potential, we aimed to isolate selective inhibitors of human kallikrein 6 (hK6). Human kallikreins (hKs) play key roles in cancer pathophysiology, but their high sequence homology and structural similarity make selective inhibition difficult^69, 70^. Among hKs, hK6 is overexpressed in several cancers^71^, driving tumor invasion and metastasis, and may be a viable target to combat neurodegeneration^72, 73^. Currently, few selective small molecule hK6 inhibitors exist^74, 75^ and hK6 is only mildly inhibited by endogenous serine protease inhibitors^76^. Moreover, reengineering inhibitors based on the amyloid precursor protein Kunitz protease inhibitor (APPI) domain results in potent but non-selective clones^77^. Note that an hK6 inhibitory monoclonal antibody is reported but not well-characterized^78^.

We transformed the Nb library with no ERS into an hK6 yeast reporter strain, as we did for TEVp. After four rounds of FACS, library enrichment occurred similar to the TEVp campaign **(Supplemental Figure 16)**. Confident that hK6 Nbs are inhibitory, we only characterized three hK6-Nb clones. All three hK6-Nbs bound to hK6 with *K_D_s* between 200 and 850 nM (**Table 3, Supplemental Table 6)**. Interestingly, among the three hK6-Nbs, only hK6-Nb1 was inhibitory. The unexpected discovery of false positives unveiled a rare failure mode of the HARP approach. We hypothesize that an otherwise non-inhibitory Nb can bind to a protease in a manner that prevents it from binding to the ER receptor^52^. Such an interaction would decrease the enzyme’s activity in the ER and could appear as “inhibitory” on flow cytometry. However, we believe such an outcome may not be common and can be mitigated by extending the linker length (i.e., (G_4_S)_3_ versus only GS in our case) between the protease C-terminus and the ERS. Another possibility is for an Nb to bind the Erd2 receptor and block protease and substrate Erd2 receptor binding. Since hK6_Nb2 and 3 bind to hK6, this latter possibility has not yet been observed.

**Table 3.**
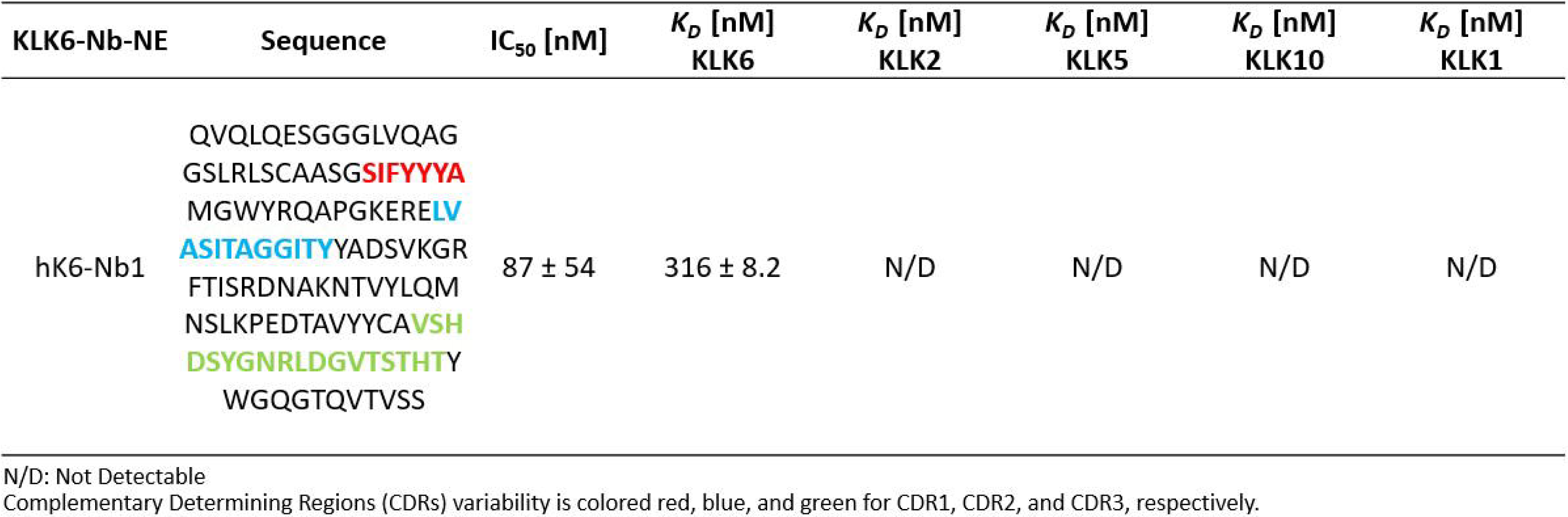
Biochemical characterization of hK6-Nb1.

| KLK6-Nb-NE | Sequence | IC<sub>50</sub> [nM] | K<sub>D</sub> [nM]<br>KLK6 | K<sub>D</sub> [nM]<br>KLK2 | K<sub>D</sub> [nM]<br>KLK5 | K<sub>D</sub> [nM]<br>KLK10 | K<sub>D</sub> [nM]<br>KLK1 |
| --- | --- | --- | --- | --- | --- | --- | --- |
| hK6-Nb1 | QVQLQESGGGLVQAG<br>GSLRLSCAASG**SIFYYYA**<br>MGWYRQAPGKEREL**V**<br>**ASITAGGITY**YADSVKGR<br>FTISRDNAKNTVYLQM<br>NSLKPEDTAVYYCA**VSH**<br>**DSYGNRLDGV****TSTHTY**<br>WGQGTQVTVSS | 87 ± 54 | 316 ± 8.2 | N/D | N/D | N/D | N/D |

N/D: Not Detectable
Complementary Determining Regions (CDRs) variability is colored red, blue, and green for CDR1, CDR2, and CDR3, respectively.

HK6-Nb1 is a potent and highly selective inhibitor of hK6, with an IC_50_ of 80 nM. Notably, it fails to bind a panel of other hKs with high sequence similarity (**Table 3**). Structural analysis based on a highly confident AlphaFold3 model of hK6_Nb1 bound to hK6 (ipTM = 0.8, pTM = 0.82) indicates that hK6-Nb1 functions as an active site-blocking Nb. Its 18-amino-acid CDR3 region is rich in charged and polar residues, including His99, His113, Asp100, Asp107, Asn104, Arg105, Ser97, Ser101, Ser111, Thr110, Thr112, and Thr114. These residues form hydrogen bonds and hydrophobic interactions with the S3, S2, and S1 subsite residues of hK6 (**Figure 5**). Notably, residues 99–104 of the CDR3 form a protruding loop (a convex-shaped paratope^25^) that directly engages the S1 pocket of hK6, while the remaining residues thread along one side of the substrate-binding cleft. This arrangement likely provides structural support and specificity to the highly flexible CDR3 loop. A critical residue in hK6_Nb1 CDR3 is Tyr102, the only tyrosine residue in its CDR3. Tyr102 fits snugly within the S1 pocket of hK6 (residues 189–195, 214–220, and 224–228, along with the catalytic triad). It forms a hydrogen bond with the backbone carboxyl group of Ser190 and engages in hydrophobic interactions with Val208 and Gln192. The positioning of Tyr102 in the S1 pocket resembles that of the broad-spectrum active site inhibitor benzamide **(Supplemental Figure 17)**, blocking the active site while being uncleavable. Our model is further validated by a co-crystal structure of hK6 with a peptide-based activity probe, whose selectivity for hK6 is accentuated by interactions with the S1, S2, and S3 pocket of hK6. Interestingly, in our case, these same pockets are engaged by the CDR3 of hK6-Nb1, although the interacting CDR3 residues are of opposite polarity. Furthermore, these binding mechanisms are markedly different from that of non-selective APPI inhibitors, which targets the S1’-S3’ subsites of hK6^77^.

**Figure 5.**
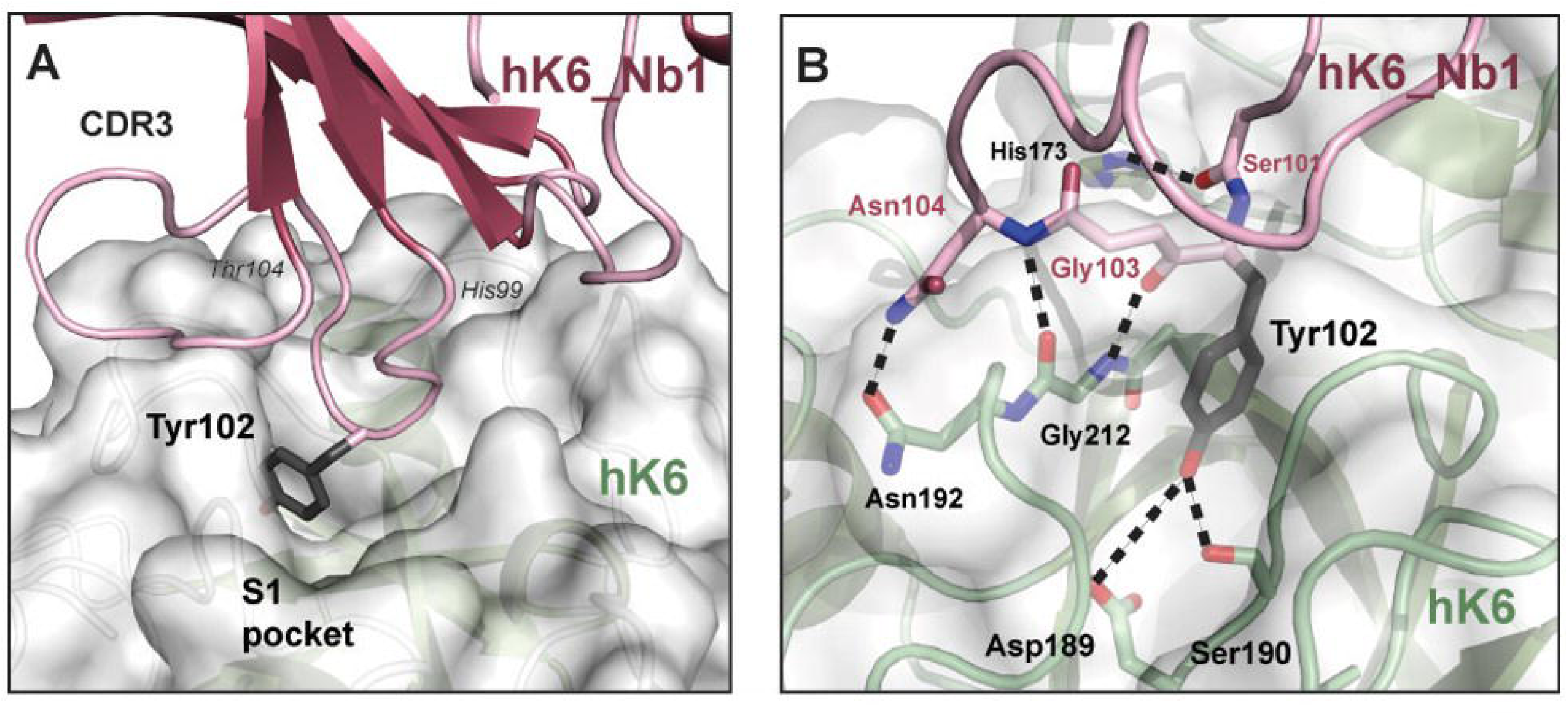
A model of hK6 inhibited by hK6-Nb1. (A) Interaction of hK6 (surface/green) and HK6-Nb1 (magenta). The CDR3 loop (His99 to Thr103) binds the surface around the substrate binding cleft and Tyr1) Detailed interactions between HK6-Nb1 (magenta) and hK6 (green). Residues 101-104 of the CDR3 loop form specific interactions in the S1 pocket, including backbone/sidechain hydrogen bonds with Tyr102 and a parallel β-sheet arrangement of Gly103-Asn104 with the protease backbone.

## Conclusion

In this study, we developed HARP, a high-throughput functional screening platform for reprogramming proteases by discovering protease inhibitory nanobodies. HARP directly reports Nb-mediated inhibition of an ER-sequestered protease via a sensitive and precise output on the yeast cell surface. This output is not mediated by signal amplification^79–82^ (transcription activation) or the action of a secondary enzymatic process (antibiotic resistance or nutritional marker enzymes)^42, 43, 83, 84^. This means that HARP’s dynamic range is only dictated by the activity of its interacting constituents (protease, inhibitor, substrate) and the steady state accumulation of the ER-transiting (un)modified substrate polypeptide on the cell surface. We estimate that HARP exhibits a linear response over a two-order-of-magnitude dynamic range for a highly ER-active protease. Therefore, optimizing HARP for a specific enzyme only requires maximizing enzyme activity and minimizing non-productive enzyme modulator interactions. Our first version of HARP uses strong galactose-inducible promoters and ERS manipulations to achieve this goal. Yet, one can imagine introducing titratable promoters on the protease and inhibitor gene cassettes^85–87^ to optimize the window and stringency of inhibitor discovery.

Through HARP, we isolated TEVp and hK6 inhibitory Nbs exhibiting high selectivity and with *K_I_s* in the low nanomolar to less than 100 nM. TEVp was chosen as proof-of-concept since it behaves predictably in the yeast ER and has been the subject of several engineering campaigns^48, 88^. Besides, these first time isolated TEVp inhibitory Nbs can be integrated into TEVp-powered genetic and protein circuits as a regulatory module^89, 90^. In the case of TEVp, all tested Nbs were inhibitory, whereas only one out of the three hK6-Nbs selected were effective when tested *in vitro*. While false positives may occur occasionally, the hit rate for HARP far surpasses binding-first platforms, in which less than 5% of binding events lead to enzyme inhibition^91^.

The design principles of HARP provided opportunities to isolate inhibitors that conventional methods may not have found, including uncompetitive inhibitors and potent inhibitors with moderate binding affinity. For example, animal immunization and selections based on binding affinity would require the presentation of a dTEV-substrate complex to isolate TEV-Nb4. Furthermore, repeated rounds on stringent binding affinity selections would have likely overlooked hK6-Nb1, a moderate binder with strong inhibitory potency^42, 43^. NGS analysis of the TEVp inhibitory Nbs for the TEVp discovery campaign confirms the system’s high dynamic range, showing a positive correlation between sequence abundance and inhibitory potency. Furthermore, ESM-2 embedding and UMAP visualization support the distinct molecular features of inhibitory versus non-inhibitory CDR3s. However, we should note that deep sequencing analysis may contain potential false positives. For instance, the top Nb sequence from Next-Gen Sequencing, NGS-Nb1, was conspicuously absent from single-clone sequencing and ultimately showed no inhibition when tested in yeast or *in vitro*.

HARP, as presented here, showcases its ability to discover protease-inhibitory nanobodies. Yet, one can imagine this concept being straightforwardly extended to other scaffolds (endogenous inhibitors, ScFvs, DARPins, Fabs, cyclic peptides), other protease targets, and other post-translational modification enzymes (kinases, acetyltransferases). Pleasingly, these binder scaffolds are already employed in traditional yeast surface display strategies^92, 93^. Additionally, the ER sequestration strategy has so far encompassed several new proteases^52, 94, 95^, kinases^96, 97^ and acetyltransferases^98^, increasing the feasibility and promise of inhibitor discovery. These precedents motivate us to expand our technology in these directions, with a focus on ER engineering to include additional enzyme targets, transcriptional control modules, and continuous evolution for inhibitor discovery and maturation.

## Acknowledgments

The authors would like to acknowledge Hunter Vu and Emily Prins for their contributions to finishing the final characterizations necessary for the manuscript.

## References

1. López-Otín, C. & Bond, J.S. Proteases: multifunctional enzymes in life and disease. J Biol Chem 283, 30433–30437 (2008).

2. Quirós, P.M., Langer, T. & López-Otín, C. New roles for mitochondrial proteases in health, ageing and disease. Nature Reviews Molecular Cell Biology 16, 345–359 (2015).

3. Reiser, J., Adair, B. & Reinheckel, T. Specialized roles for cysteine cathepsins in health and disease. The Journal of Clinical Investigation 120, 3421–3431 (2010).

4. López-Otín, C. & Matrisian, L.M. Emerging roles of proteases in tumour suppression. Nature Reviews Cancer 7, 800–808 (2007).

5. Pham, C.T.N. Neutrophil serine proteases: specific regulators of inflammation. Nature Reviews Immunology 6, 541–550 (2006).

6. Pickart, C.M. & Cohen, R.E. Proteasomes and their kin: proteases in the machine age. Nature Reviews Molecular Cell Biology 5, 177–187 (2004).

7. Drag, M. & Salvesen, G.S. Emerging principles in protease-based drug discovery. Nature Reviews Drug Discovery 9, 690–701 (2010).

8. Powers, J.C., Asgian, J.L., Ekici, O.D. & James, K.E. Irreversible inhibitors of serine, cysteine, and threonine proteases. Chem Rev 102, 4639–4750 (2002).

9. Hu, J., Van den Steen, P.E., Sang, Q.X. & Opdenakker, G. Matrix metalloproteinase inhibitors as therapy for inflammatory and vascular diseases. Nat Rev Drug Discov 6, 480–498 (2007).

10. Turk, B. Targeting proteases: successes, failures and future prospects. Nat Rev Drug Discov 5, 785–799 (2006).

11. Leung, D., Abbenante, G. & Fairlie, D.P. Protease Inhibitors:[] Current Status and Future Prospects. Journal of Medicinal Chemistry 43, 305–341 (2000).

12. Anderson, J., Schiffer, C., Lee, S.K. & Swanstrom, R. Viral protease inhibitors. Handb Exp Pharmacol 189, 85–110 (2009).

13. Manasanch, E.E. & Orlowski, R.Z. Proteasome inhibitors in cancer therapy. Nat Rev Clin Oncol 14, 417–433 (2017).

14. Xie, L., Xie, L. & Bourne, P.E. Structure-based systems biology for analyzing off-target binding. Curr Opin Struct Biol 21, 189–199 (2011).

15. Deu, E., Verdoes, M. & Bogyo, M. New approaches for dissecting protease functions to improve probe development and drug discovery. Nat Struct Mol Biol 19, 9–16 (2012).

16. Vandenbroucke, R.E. & Libert, C. Is there new hope for therapeutic matrix metalloproteinase inhibition? Nat Rev Drug Discov 13, 904–927 (2014).

17. Moussa-Pacha, N.M., Abdin, S.M., Omar, H.A., Alniss, H. & Al-Tel, T.H. BACE1 inhibitors: Current status and future directions in treating Alzheimer’s disease. Med Res Rev 40, 339–384 (2020).

18. Govindaraj, R.G., Thangapandian, S., Schauperl, M., Denny, R.A. & Diller, D.J. Recent applications of computational methods to allosteric drug discovery. Frontiers in Molecular Biosciences 9 (2023).

19. Niphakis, M.J. & Cravatt, B.F. Enzyme inhibitor discovery by activity-based protein profiling. Annu Rev Biochem 83, 341–377 (2014).

20. Merdanovic, M., Mönig, T., Ehrmann, M. & Kaiser, M. Diversity of allosteric regulation in proteases. ACS Chem Biol 8, 19–26 (2013).

21. Tee, W.-V. & Berezovsky, I.N. Allosteric drugs: New principles and design approaches. Current Opinion in Structural Biology 84, 102758 (2024).

22. Farady, C.J. & Craik, C.S. Mechanisms of macromolecular protease inhibitors. Chembiochem 11, 2341–2346 (2010).

23. Stoop, A.A. & Craik, C.S. Engineering of a macromolecular scaffold to develop specific protease inhibitors. Nature Biotechnology 21, 1063–1068 (2003).

24. Scott, C.J. & Taggart, C.C. Biologic protease inhibitors as novel therapeutic agents. Biochimie 92, 1681–1688 (2010).

25. Nam, D.H., Rodriguez, C., Remacle, A.G., Strongin, A.Y. & Ge, X. Active-site MMP-selective antibody inhibitors discovered from convex paratope synthetic libraries. Proceedings of the National Academy of Sciences 113, 14970–14975 (2016).

26. Gerhardy, S. et al. Allosteric inhibition of HTRA1 activity by a conformational lock mechanism to treat age-related macular degeneration. Nature Communications 13, 5222 (2022).

27. El Debs, B., Utharala, R., Balyasnikova, I.V., Griffiths, A.D. & Merten, C.A. Functional single-cell hybridoma screening using droplet-based microfluidics. Proc Natl Acad Sci U S A 109, 11570–11575 (2012).

28. Chestukhin, A. & DeCaprio, J.A. Western blot screening for monoclonal antibodies against human separase. Journal of Immunological Methods 274, 105–113 (2003).

29. Lim, C.C., Woo, P.C.Y. & Lim, T.S. Development of a Phage Display Panning Strategy Utilizing Crude Antigens: Isolation of MERS-CoV Nucleoprotein human antibodies. Scientific Reports 9, 6088 (2019).

30. Chen, S. et al. Identification of highly selective covalent inhibitors by phage display. Nat Biotechnol 39, 490–498 (2021).

31. Hawinkels, L.J.A.C. et al. Efficient degradation-aided selection of protease inhibitors by phage display. Biochemical and Biophysical Research Communications 364, 549–555 (2007).

32. Wang, C.I., Yang, Q. & Craik, C.S. Phage display of proteases and macromolecular inhibitors. Methods Enzymol 267, 52–68 (1996).

33. Markland, W., Roberts, B.L. & Ladner, R.C. in Methods in Enzymology, Vol. 267 28–51 (Academic Press, 1996).

34. Aoki, W. et al. High-throughput screening of improved protease inhibitors using a yeast cell surface display system and a yeast cell chip. J Biosci Bioeng 111, 16–18 (2011).

35. Glotzbach, B. et al. Combinatorial optimization of cystine-knot peptides towards high-affinity inhibitors of human matriptase-1. PLoS One 8, e76956 (2013).

36. Lopez-Morales, J. et al. Protein Engineering and High-Throughput Screening by Yeast Surface Display: Survey of Current Methods. Small Science 3, 2300095 (2023).

37. Diamante, L., Gatti-Lafranconi, P., Schaerli, Y. & Hollfelder, F. In vitro affinity screening of protein and peptide binders by megavalent bead surface display. Protein Eng Des Sel 26, 713–724 (2013).

38. Machon, U. et al. On-bead screening of a combinatorial fumaric acid derived peptide library yields antiplasmodial cysteine protease inhibitors with unusual peptide sequences. J Med Chem 52, 5662–5672 (2009).

39. Bacon, K., Burroughs, M., Blain, A., Menegatti, S. & Rao, B.M. Screening Yeast Display Libraries against Magnetized Yeast Cell Targets Enables Efficient Isolation of Membrane Protein Binders. ACS Combinatorial Science 21, 817–832 (2019).

40. Chavarria-Smith, J. et al. Dual antibody inhibition of KLK5 and KLK7 for Netherton syndrome and atopic dermatitis. Sci Transl Med 14, eabp9159 (2022).

41. Demeestere, D. et al. Development and Validation of a Small Single-domain Antibody That Effectively Inhibits Matrix Metalloproteinase 8. Mol Ther 24, 890–902 (2016).

42. Gal-Tanamy, M. et al. HCV NS3 serine protease-neutralizing single-chain antibodies isolated by a novel genetic screen. J Mol Biol 347, 991–1003 (2005).

43. Lopez, T. et al. Functional selection of protease inhibitory antibodies. Proceedings of the National Academy of Sciences 116, 16314–16319 (2019).

44. Bond, J.S. Proteases: History, discovery, and roles in health and disease. Journal of Biological Chemistry 294, 1643–1651 (2019).

45. Yi, L. et al. Engineering of TEV protease variants by yeast ER sequestration screening (YESS) of combinatorial libraries. Proc Natl Acad Sci U S A 110, 7229–7234 (2013).

46. Li, Q. et al. Profiling Protease Specificity: Combining Yeast ER Sequestration Screening (YESS) with Next Generation Sequencing. ACS Chemical Biology 12, 510–518 (2017).

47. Denard, C.A. et al. YESS 2.0, a Tunable Platform for Enzyme Evolution, Yields Highly Active TEV Protease Variants. ACS Synthetic Biology 10, 63–71 (2021).

48. Martinusen, S.G. et al. Modular and integrative activity reporters enhance biochemical studies in the yeast ER. Protein Engineering, Design and Selection 37 (2024).

49. Alcala-Torano, R. et al. Yeast Display Enables Identification of Covalent Single-Domain Antibodies against Botulinum Neurotoxin Light Chain A. ACS Chemical Biology 17, 3435–3449 (2022).

50. Tom, I. et al. Development of a therapeutic anti-HtrA1 antibody and the identification of DKK3 as a pharmacodynamic biomarker in geographic atrophy. Proceedings of the National Academy of Sciences 117, 9952–9963 (2020).

51. Atwal, J.K. et al. A Therapeutic Antibody Targeting BACE1 Inhibits Amyloid-β Production in Vivo. Science Translational Medicine 3, 84ra43–84ra43 (2011).

52. Mei, M. et al. Characterization of aromatic residue-controlled protein retention in the endoplasmic reticulum of Saccharomyces cerevisiae. J Biol Chem 292, 20707–20719 (2017).

53. Armstrong, L.A. et al. Biochemical characterization of protease activity of Nsp3 from SARS-CoV-2 and its inhibition by nanobodies. PLoS One 16, e0253364 (2021).

54. Uchański, T., Pardon, E. & Steyaert, J. Nanobodies to study protein conformational states. Current Opinion in Structural Biology 60, 117–123 (2020).

55. Lan, T. et al. Discovery of Thioether-Cyclized Macrocyclic Covalent Inhibitors by mRNA Display. Journal of the American Chemical Society 146, 24053–24060 (2024).

56. Gutierrez-Campos, R., Torres-Acosta, J.A., Saucedo-Arias, L.J. & Gomez-Lim, M.A. The use of cysteine proteinase inhibitors to engineer resistance against potyviruses in transgenic tobacco plants. Nature Biotechnology 17, 1223–1226 (1999).

57. Murry, L.E. et al. Transgenic Corn Plants Expressing MDMV Strain B Coat Protein are Resistant to Mixed Infections of Maize Dwarf Mosaic Virus and Maize Chlorotic Mottle Virus. Bio/Technology 11, 1559–1564 (1993).

58. McMahon, C. et al. Yeast surface display platform for rapid discovery of conformationally selective nanobodies. Nature Structural & Molecular Biology 25, 289–296 (2018).

59. Phan, J. et al. Structural basis for the substrate specificity of tobacco etch virus protease. J Biol Chem 277, 50564–50572 (2002).

60. Abramson, J. et al. Accurate structure prediction of biomolecular interactions with AlphaFold 3. Nature 630, 493–500 (2024).

61. Yuan, S. et al. Structure of the HRV-C 3C-Rupintrivir Complex Provides New Insights for Inhibitor Design. Virol Sin 35, 445–454 (2020).

62. Li, F. et al. Procleave: Predicting Protease-specific Substrate Cleavage Sites by Combining Sequence and Structural Information. Genomics, Proteomics & Bioinformatics 18, 52–64 (2020).

63. Reis, P.B.P.S. et al. Antibody-Antigen Binding Interface Analysis in the Big Data Era. Frontiers in Molecular Biosciences 9 (2022).

64. Deng, J. et al. Nanobody–antigen interaction prediction with ensemble deep learning and prompt-based protein language models. Nature Machine Intelligence 6, 1594–1604 (2024).

65. Adib-Conquy, M., Gilbert, M. & Avrameas, S. Effect of amino acid substitutions in the heavy chain CDR3 of an autoantibody on its reactivity. International Immunology 10, 341–346 (1998).

66. Haidar, J.N. et al. Backbone Flexibility of CDR3 and Immune Recognition of Antigens. Journal of Molecular Biology 426, 1583–1599 (2014).

67. Rives, A. et al. Biological structure and function emerge from scaling unsupervised learning to 250 million protein sequences. Proceedings of the National Academy of Sciences 118, e2016239118 (2021).

68. Healy, J. & McInnes, L. Uniform manifold approximation and projection. Nature Reviews Methods Primers 4, 82 (2024).

69. Paliouras, M., Borgono, C. & Diamandis, E.P. Human tissue kallikreins: The cancer biomarker family. Cancer Letters 249, 61–79 (2007).

70. Borgoño, C.A. & Diamandis, E.P. The emerging roles of human tissue kallikreins in cancer. Nat Rev Cancer 4, 876–890 (2004).

71. Radisky, E.S. Extracellular proteolysis in cancer: Proteases, substrates, and mechanisms in tumor progression and metastasis. J Biol Chem 300, 107347 (2024).

72. Yoon, H. et al. Kallikrein 6 signals through PAR1 and PAR2 to promote neuron injury and exacerbate glutamate neurotoxicity. J Neurochem 127, 283–298 (2013).

73. Mella, C., Figueroa, C.D., Otth, C. & Ehrenfeld, P. Involvement of Kallikrein-Related Peptidases in Nervous System Disorders. Frontiers in Cellular Neuroscience 14 (2020).

74. Aït Amiri, S., et al. Identification of First-in-Class Inhibitors of Kallikrein-Related Peptidase 6 That Promote Oligodendrocyte Differentiation. J Med Chem 64, 5667–5688 (2021).

75. Zhang, L. et al. A KLK6 Activity-Based Probe Reveals a Role for KLK6 Activity in Pancreatic Cancer Cell Invasion. Journal of the American Chemical Society 144, 22493–22504 (2022).

76. Magklara, A. et al. Characterization of the enzymatic activity of human kallikrein 6: autoactivation, substrate specificity, and regulation by inhibitors. Biochemical and Biophysical Research Communications 307, 948–955 (2003).

77. Sananes, A. et al. A potent, proteolysis-resistant inhibitor of kallikrein-related peptidase 6 (KLK6) for cancer therapy, developed by combinatorial engineering. J Biol Chem 293, 12663–12680 (2018).

78. Ghosh, M.C., Grass, L., Soosaipillai, A., Sotiropoulou, G. & Diamandis, E.P. Human Kallikrein 6 Degrades Extracellular Matrix Proteins and May Enhance the Metastatic Potential of Tumour Cells. Tumor Biology 25, 193–199 (2004).

79. Wang, W. et al. A light- and calcium-gated transcription factor for imaging and manipulating activated neurons. Nature Biotechnology 35, 864–871 (2017).

80. Algar, W.R., Hildebrandt, N., Vogel, S.S. & Medintz, I.L. FRET as a biomolecular research tool — understanding its potential while avoiding pitfalls. Nature Methods 16, 815–829 (2019).

81. Tracy, B.P., Gaida, S.M. & Papoutsakis, E.T. Flow cytometry for bacteria: enabling metabolic engineering, synthetic biology and the elucidation of complex phenotypes. Current Opinion in Biotechnology 21, 85–99 (2010).

82. Sanchez, M.I. & Ting, A.Y. Directed evolution improves the catalytic efficiency of TEV protease. Nat Methods 17, 167–174 (2020).

83. Carrico, Z.M., Strobel, K.L., Atreya, M.E., Clark, D.S. & Francis, M.B. Simultaneous selection and counter-selection for the directed evolution of proteases in E. coli using a cytoplasmic anchoring strategy. Biotechnology and Bioengineering 113, 1187–1193 (2016).

84. Cottier, V., Barberis, A. & Lüthi, U. Novel yeast cell-based assay to screen for inhibitors of human cytomegalovirus protease in a high-throughput format. Antimicrob Agents Chemother 50, 565–571 (2006).

85. Shaw, W.M., Khalil, A.S. & Ellis, T. A Multiplex MoClo Toolkit for Extensive and Flexible Engineering of Saccharomyces cerevisiae. ACS Synth Biol 12, 3393–3405 (2023).

86. Lee, M.E., DeLoache, W.C., Cervantes, B. & Dueber, J.E. A Highly Characterized Yeast Toolkit for Modular, Multipart Assembly. ACS Synth Biol 4, 975–986 (2015).

87. Ottoz, D.S., Rudolf, F. & Stelling, J. Inducible, tightly regulated and growth condition-independent transcription factor in Saccharomyces cerevisiae. Nucleic Acids Res 42, e130 (2014).

88. Denard, C.A. et al. YESS 2.0, a Tunable Platform for Enzyme Evolution, Yields Highly Active TEV Protease Variants. ACS Synth Biol 10, 63–71 (2021).

89. Chen, Z. & Elowitz, M.B. Programmable protein circuit design. Cell 184, 2284–2301 (2021).

90. Fernandez-Rodriguez, J. & Voigt, C.A. Post-translational control of genetic circuits using Potyvirus proteases. Nucleic Acids Res 44, 6493–6502 (2016).

91. Chavarria-Smith, J. et al. Dual antibody inhibition of KLK5 and KLK7 for Netherton syndrome and atopic dermatitis. Science Translational Medicine 14, eabp9159 (2022).

92. Li, Y., Wang, X., Zhou, N.-Y. & Ding, J. Yeast surface display technology: Mechanisms, applications, and perspectives. Biotechnology Advances 76, 108422 (2024).

93. Raeeszadeh-Sarmazdeh, M. & Boder, E.T. in Yeast Surface Display. (ed. M.W. Traxlmayr) 3-25 (Springer US, New York, NY; 2022).

94. Lubin, J.H., et al. A comprehensive survey of coronaviral main protease active site diversity in 3D: Identifying and analyzing drug discovery targets in search of broad specificity inhibitors for the next coronavirus pandemic. bioRxiv (2023).

95. Yaghi, R.M., Andrews, C.L., Wylie, D.C. & Iverson, B.L. High-Resolution Substrate Specificity Profiling of SARS-CoV-2 Mpro; Comparison to SARS-CoV Mpro. ACS Chemical Biology 19, 1474–1483 (2024).

96. Ezagui, J., Russell, B., Mairena, Y. & Stern, L.A. Endoplasmic reticulum sequestration empowers phosphorylation profiling on the yeast surface. AIChE Journal 68, e17931 (2022).

97. Taft, J.M. et al. Rapid Screen for Tyrosine Kinase Inhibitor Resistance Mutations and Substrate Specificity. ACS Chemical Biology 14, 1888–1895 (2019).

98. Martinusen, S.G. & Denard, C.A. Leveraging yeast sequestration to study and engineer posttranslational modification enzymes. Biotechnol Bioeng 121, 903–914 (2024).

