## Supplemental File for "High-throughput Activity Reprogramming of Proteases (HARP)"

**Running title: A Yeast Platform for the Discovery of Protease Inhibitory Macromolecules**

Samantha G Martinusen^1^, Ethan W Slaton^1^, Seyednima Ajayebi^1^, Marian A Pulgar^1^, Cassidy F Simas^2^, Sage E Nelson^1^, Julia T Besu^3^, Amit Dutta^4^, Steven Bruner^4^, Carl A Denard^1,5^ *

^1^Department of Chemical Engineering, University of Florida, Gainesville, 32611, USA

^2^J. Crayton Pruitt Family Department of Biomedical Engineering, University of Florida, Gainesville, 32611, USA

^3^Department of Biology, University of Florida, Gainesville, 32611, USA

^4^Department of Chemistry, University of Florida, Gainesville, 32611, USA

^5^UF Health Cancer Center, University of Florida, Gainesville, 32611, USA

*Carl A Denard:

**Supplemental Figures**

Supplemental Figure 1. MMP8 inhibition based on ER retention.

Supplemental Figure 2. FACS plots representing MMP8 inhibition based on ER retention.

Supplemental Figure 3. Gating strategy for MMP8 inhibition mock sort.

Supplemental Figure 4. Flow plots for each FACS round of the MMP8 inhibition mock sort.

Supplemental Figure 5. Sequence alignment of the isolated Nb constructs from the MMP8 mock sort.

Supplemental Figure 6. NbLibrary_TEVp FACS enrichment.

Supplemental Figure 7. Impact of ERS on inhibitory phenotype observed in yeast assay.

Supplemental Figure 8. Gating strategy for the TEVp Nb Library sort.

Supplemental Figure 9. Enrichment gating strategy for the TEVp Nb Library sort.

Supplemental Figure 10. IC50 curves were determined via FRET assays for isolated TEVp-Nbs.

Supplemental Figure 11. BLI curves for TEVp-Nb v dTEVp.

Supplemental Figure 12. BLI curves for TEVp-Nb v TEVp.

Supplemental Figure 13. Alphafold2 models of TEVp-Nbs.

Supplemental Figure 14. TEVp-Nb4 complex structure predicted by AlphaFold3.

Supplemental Figure 15. Frequency of amino acids in the CDR3 for different fixed amino acids in the final enrichment population of NbLibrary_NE_TEVp.

Supplemental Figure 16. KLK6-NbLibrary-NE FACS enrichment.

Supplemental Figure 17. Alphafold prediction of hK6_Nb1 interaction overlayed on hK6 Active Form with benzamidine inhibitor (PBD: 1L2E).

Supplemental Figure 18. SDS-PAGE gel of purified Nbs from NbLibrary_NE_TEVp.

Supplemental Figure 19. SDS-PAGE of purified TEVp for kinetic assays.

Supplemental Figure 20. FRET kinetic assay comparison of purified TEVp to purchased TEVp.

Supplemental Figure 21. Kinetic characterization of TEVp.

Supplemental Figure 22. BLI curve for Nb.b201 v HSA.

Supplemental Figure 23. BLI curves for TEVp-Nb v TVMVp assays.

Supplemental Figure 24. BLI curves for hK6-Nbs v KLK6.

Supplemental Figure 25. IC50 curve determined via FRET assays for the hK6-Nb1 against KLK6.

Supplemental Figure 27. BLI curves for hK6-Nb1 v KLK variants.

Supplemental Figure 28. Properties of CDR3 regions in the final enrichment against naïve library.

**Supplemental Tables**

Supplemental Table 1. Plasmids referenced in the manuscript.

Supplemental Table 2. CDR3 enrichment of isolated TEVp-Nbs.

Supplemental Table 3. Read counts of the top fifty TEVp-Nb sequences in ENR4.

Supplemental Table 4. Binding characterization of the isolated TEVp-Nbs.

Supplemental Table 5. CDR3 sequences in round 4 containing ENQ motif.

Supplemental Table 6. Potency characterization of the isolated hK6-Nbs against KLK6.

Supplemental Table 7. Primer sequences for the assemblies referenced in the manuscript.

Supplemental Table 8. Results of control BLI assay, exploring the influence of trace amounts of glycerol on resulting disassociation constant.

**Supplemental Text**

Methods and Materials

Supplemental References

Supplemental Figures

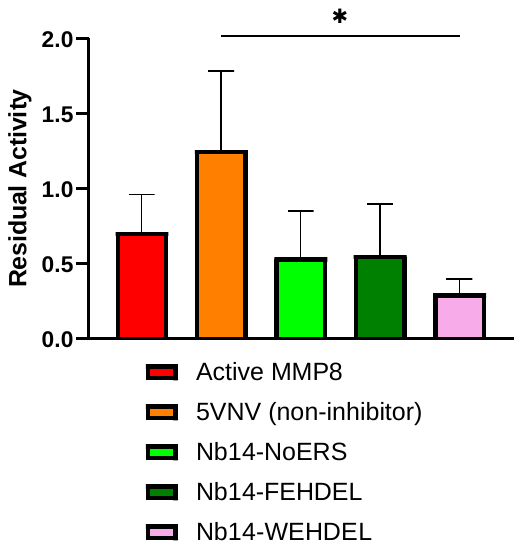

**Supplemental Figure 1. MMP8 inhibition based on ER retention.** Quantification of observed inhibition of MMP8 as a function of Nb14 ER retention. Data normalized to active MMP8 (anti-FLAG/anti-HA).

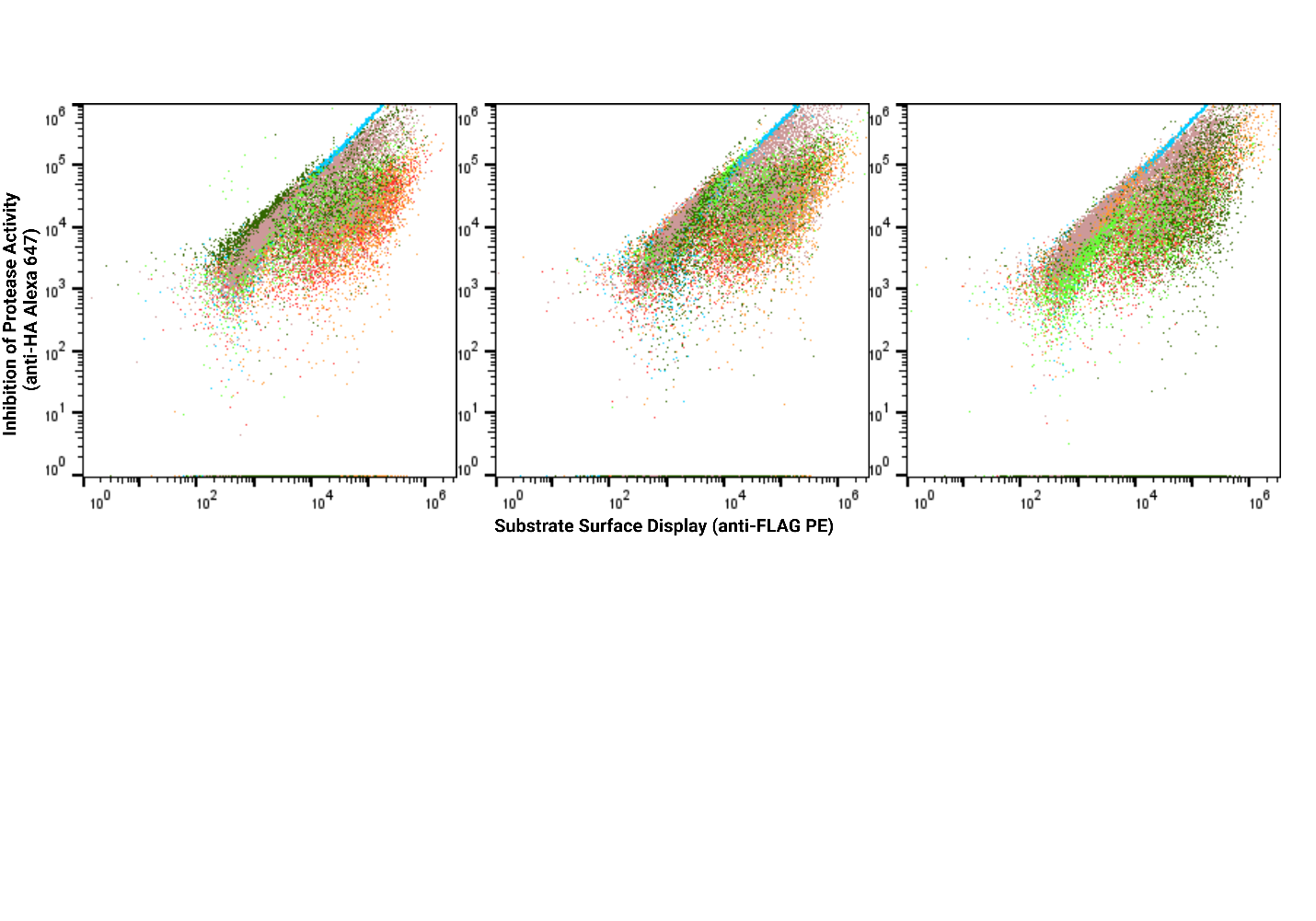

**Supplemental Figure 2. FACS plots representing MMP8 inhibition based on ER retention.** Plots representing each sample (N=3) in the MMP8 inhibition based on ER retention experiment. The colors of populations are as follows: MMP8 Off (blue), MMP8 Active (red), 5VNV (orange), Nb14-NoERS (light green), Nb14-FEHDEL (dark green), and Nb14-WEHDEL (pink). Dot plots are represented here on a log scale.

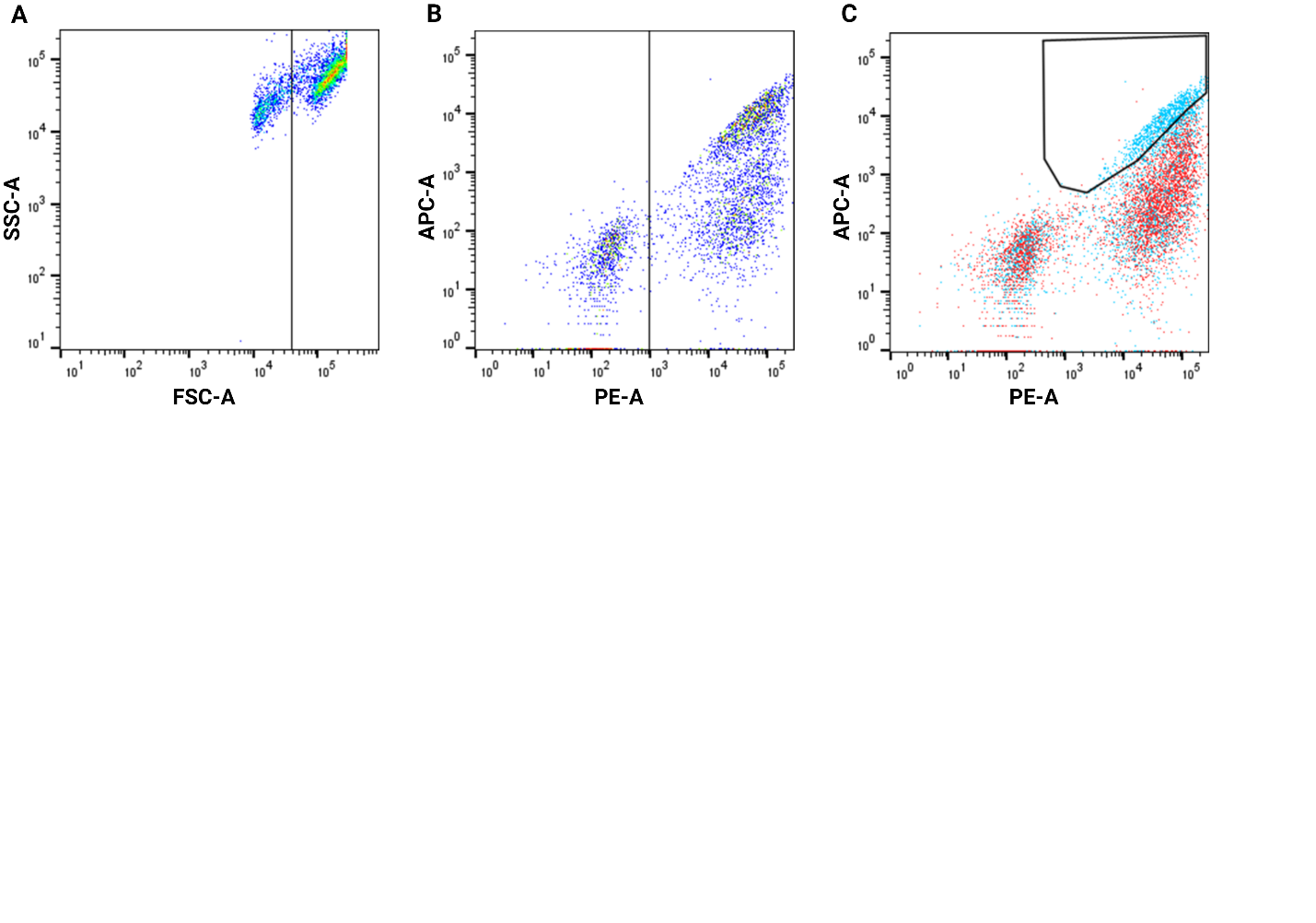

**Supplemental Figure 3. Gating strategy for MMP8 inhibition mock sort.** (A) The yeast cell gate captures all living cells and displays them. The cell population is one sample of the MMP8-Nb14 cells. (B) Displaying cell gate capturing cells displaying peptides on the surface of the cell. The cell population is the same sample of the MMP8-Nb14 cells as in (A). (C) Inhibition gate capturing cells displaying fully intact substrate cassette, displaying high levels of anti-FLAG and anti-HA fluorescence. The light blue cell population is the same sample of the MMP8-Nb14 cells as in (A), and the red cell population is one sample of the MMP8-5vnv cells. Dot plots are represented here on a log scale with PE-A corresponding to anti-FLAG labeling and APC-A corresponding to anti-HA labeling.

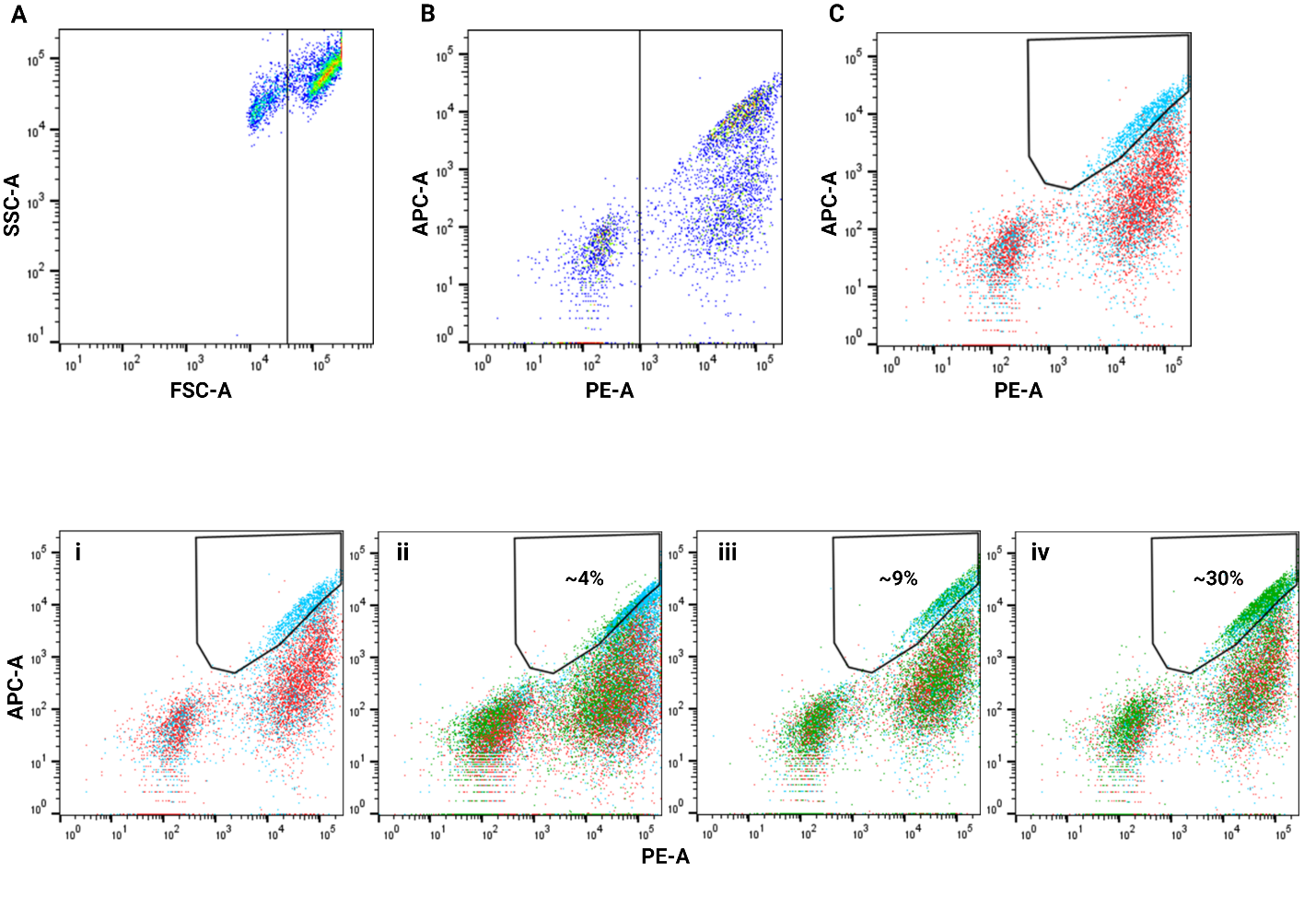

**Supplemental Figure 4. Flow plots for each FACS round of the MMP8 inhibition mock sort.** (i) Examples of the control populations that were run each round: MMP8-Nb14 (positive control) population represented in light blue and MMP8-5vnv (negative control) represented in red. (ii) Initial mock library (represented in dark green) overlaying the control populations, with approximately 4% of cells falling within the sort gate. (iii) Enrichment 1 (ENR1) of the mock library (represented in dark green) overlays the control populations, with approximately 9% of cells falling within the sort gate. (iv) Enrichment 2 (ENR2) of the mock library (represented in dark green) overlays the control populations, with approximately 30% of cells falling within the sort gate. Dot plots are represented here on a log scale with PE-A corresponding to anti-FLAG labeling and APC-A corresponding to anti-HA labeling.

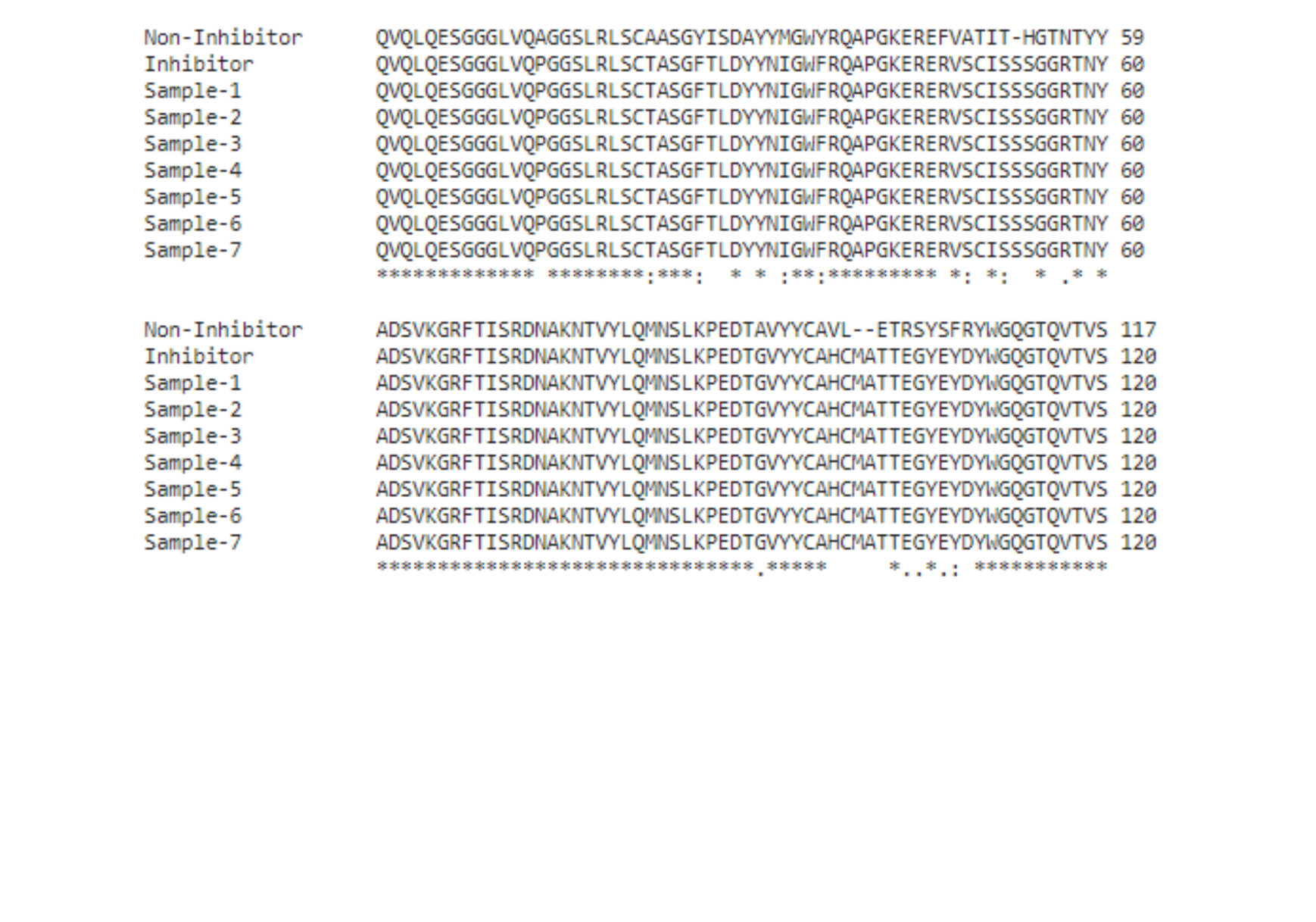

**Supplemental Figure 5. Sequence alignment of the isolated Nb constructs from the MMP8 mock sort.** Amino acid alignment of isolated Nb plasmids from MMP8 Mock Sort RND2 enriched population illustrates all isolated Nbs are identical to the strong inhibitor (Inhibitor sequence, line two).

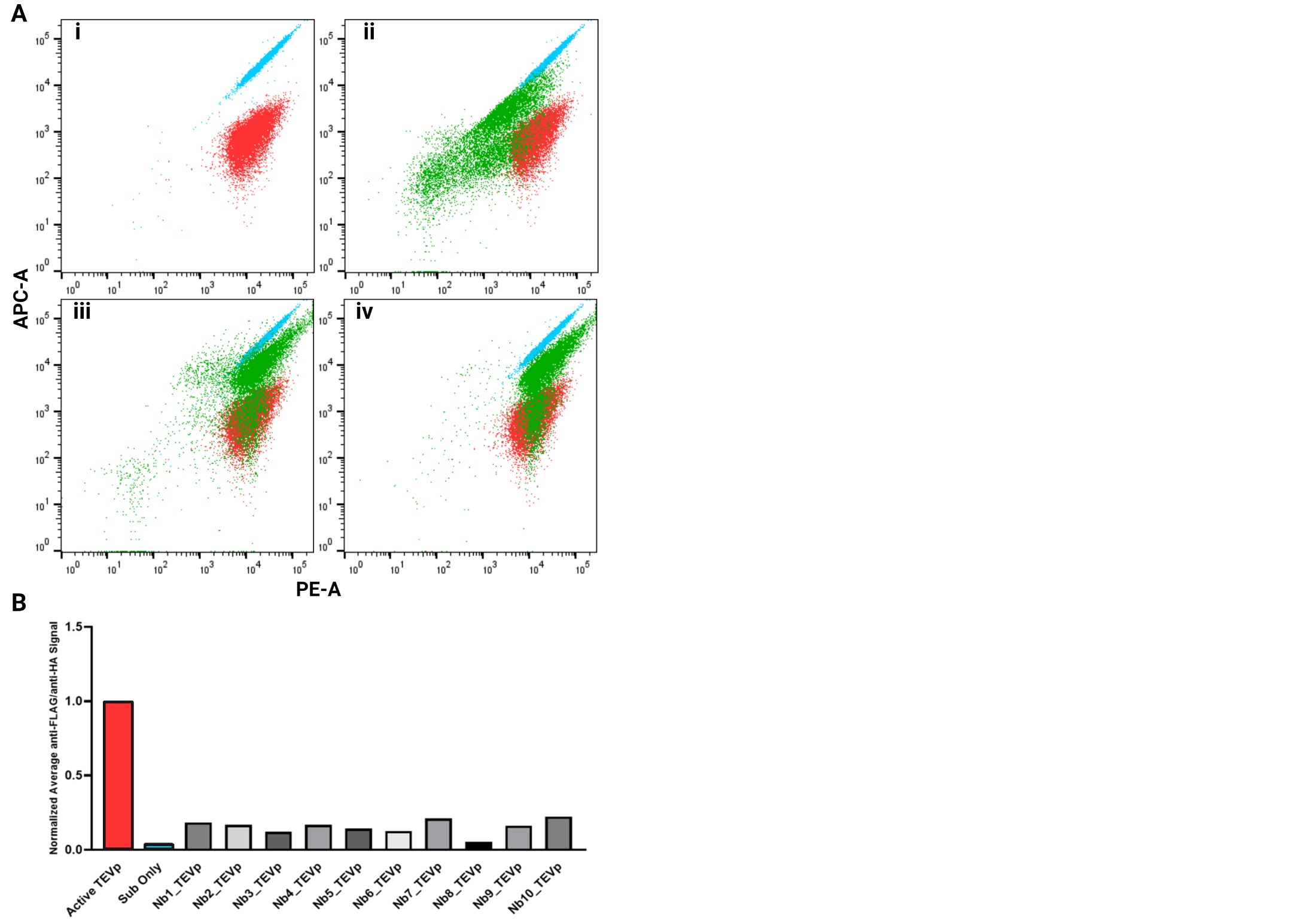

**Supplemental Figure 6. NbLibrary_TEVp FACS enrichment.** (A) Flow plots for each FACS round of the NbLibrary_TEVp sort. (Ai) Illustrations of the positive (blue) and negative (red) controls. (Aii-iv) Corresponding flow plots of the (ii) Initial Library, (iii) Enrichment 1, and (iv) Enrichment 2, with NbLibrary_TEVp represented in green. (B) Representation of the anti-FLAG/anti-HA ratio (normalized to active TEVp protease) of the selected constructs isolated from NbLibrary_TEVp. Dot plots are represented here on a log scale, with PE-A corresponding to anti-FLAG labeling and APC-A corresponding to anti-HA labeling.

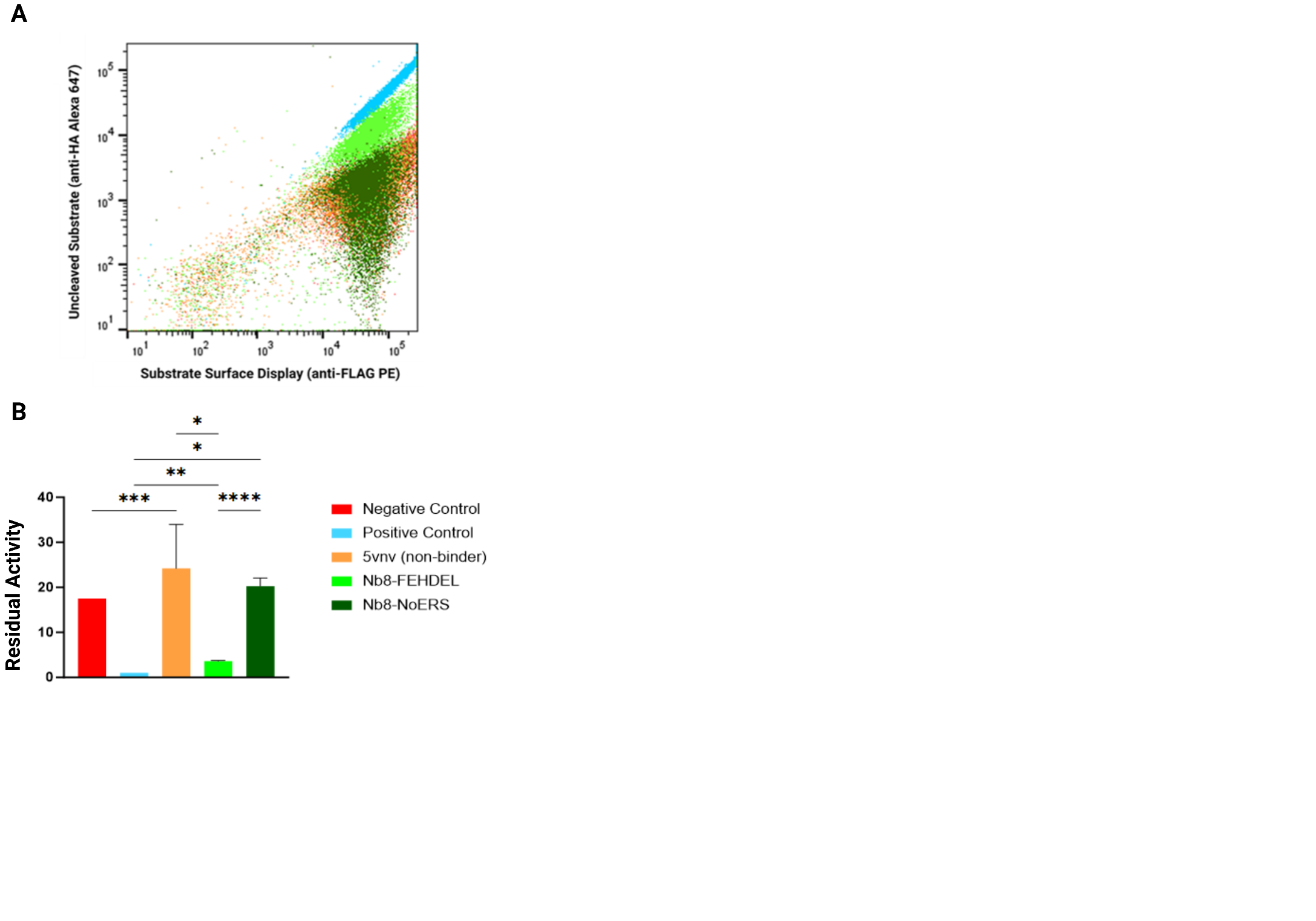

**Supplemental Figure 7. Impact of ERS on inhibitory phenotype observed in yeast assay. (**A) Flow dot plot representation of our positive control (TEVp substrate integrated, blue), negative control (TEVp active protease integrated, red), non-inhibitory Nb (orange), Nb8-TEVp (with the FEHDEL ERS, light green), and Nb8-TEVp-STOP (no ERS, dark green). (B) Fold-change quantification of the inhibition ratio (anti-FLAG/anti-HA normalized to active TEVp). Colors correspond with the dot plot in (A). Dot plots are represented in a log scale with PE-A corresponding with anti-FLAG labeling and APC-A corresponding with anti-HA labeling.

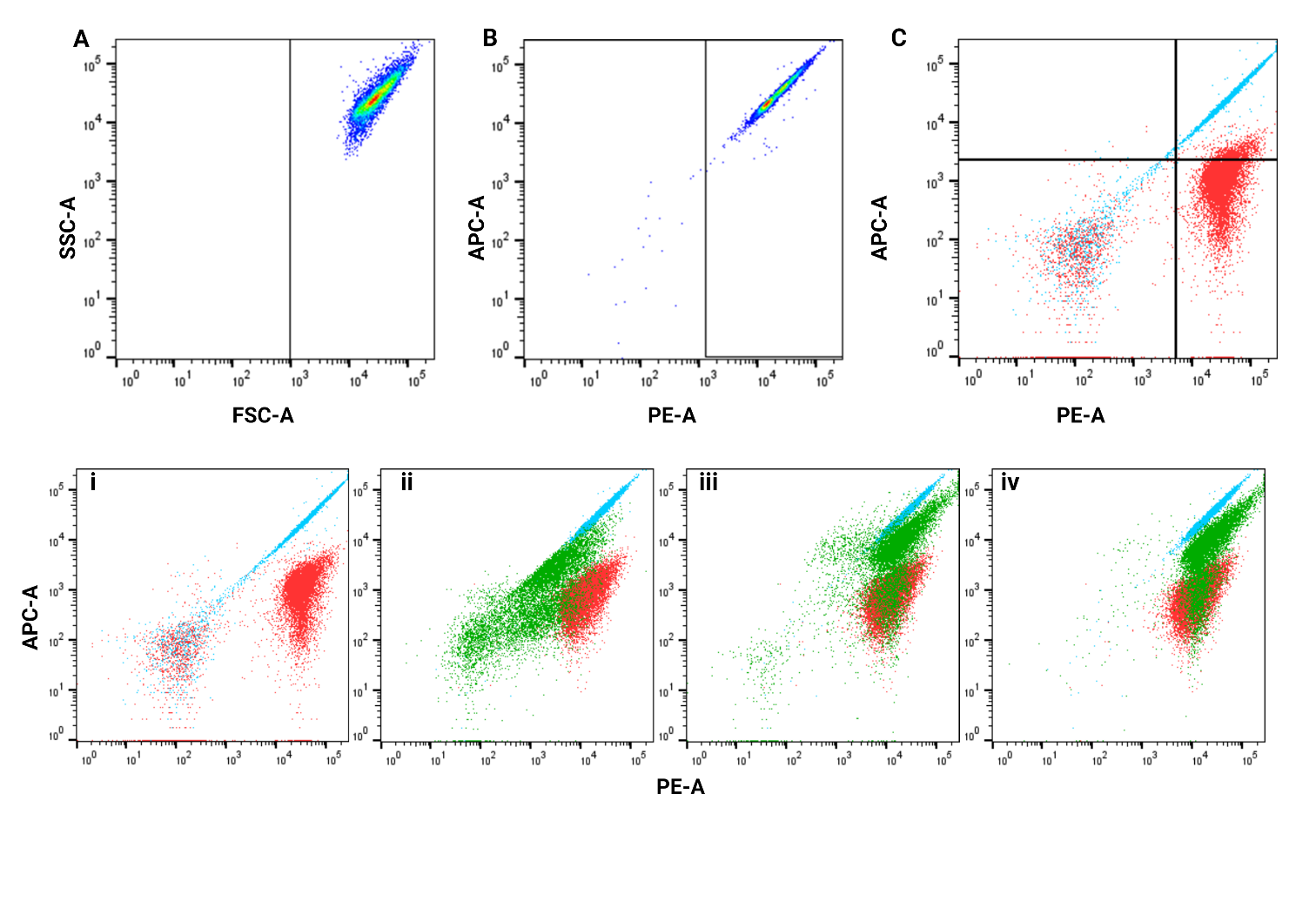

**Supplemental Figure 8. Gating strategy for the TEVp Nb Library sort.** (A) The yeast cell gate captures all living cells and displays them. The cell population is the TEVs control (positive control) population. (B) Displaying cell gate capturing cells displaying peptides on the surface of the cell. The cell population is the same as the TEVs control (positive control) population sample in (A). (C) Inhibition gate capturing cells displaying fully intact substrate cassette, displaying high levels of anti-FLAG and anti-HA fluorescence. The blue cell population is the same sample as the TEV control (positive control) population in (A), and the red cell population is the TEVp control (negative control) population. Dot plots are represented here on a log scale with PE-A corresponding to anti-FLAG labeling and APC-A corresponding to anti-HA labeling.

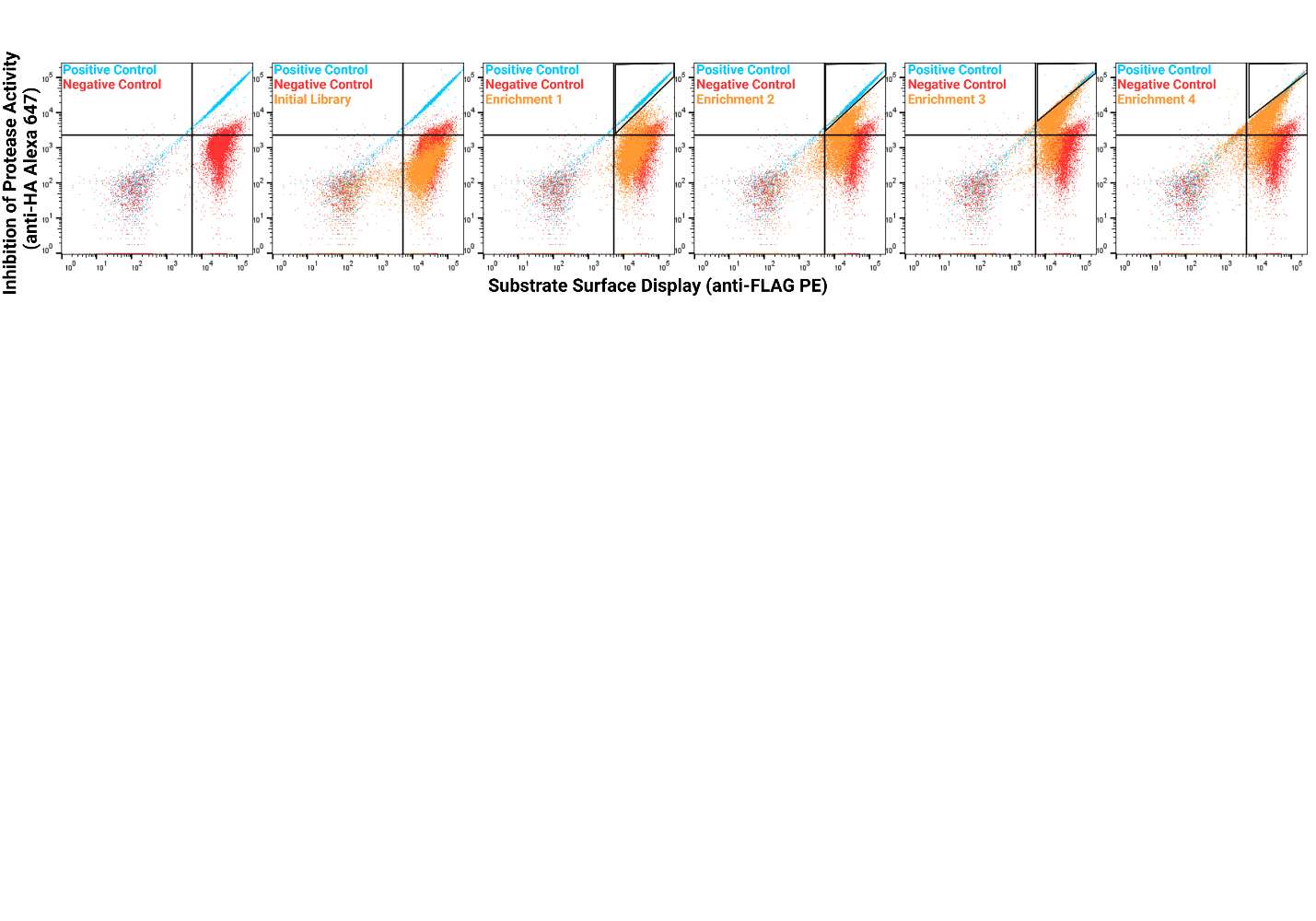

**Supplemental Figure 9. Enrichment gating strategy for the TEVp Nb Library sort.** Gating strategy for approximately 1% of cells (orange population) to enrich the inhibitory phenotype. Dot plots are represented here on a log scale.

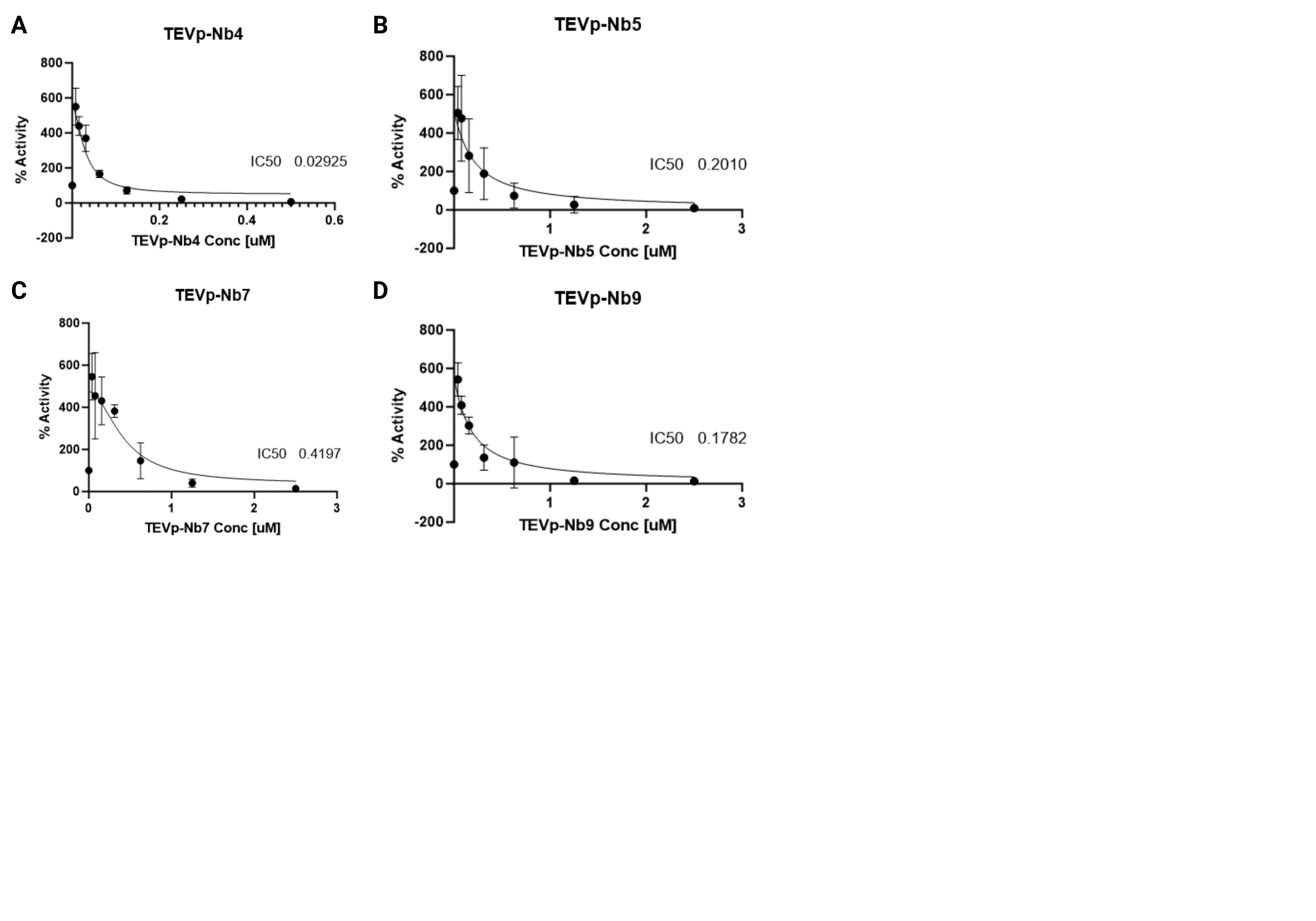

**Supplemental Figure 10. IC50 curves were determined via FRET assays for the isolated TEVp-Nbs.** All inhibition assays were conducted using a set concentration of TEVp [2 uM] and Abz-DNP substrate [20 uM]. Triplicates were measured for each TEVp-Nb, the average IC50 represented on each plot, and the errors were propagated by taking the standard deviation of the IC50 calculated for each sample. Nb-NE identifies are as follows: (A) TEVp-Nb4, (B) TEVp-Nb5, (C) TEVp-Nb7, and (D) TEVp-Nb9.

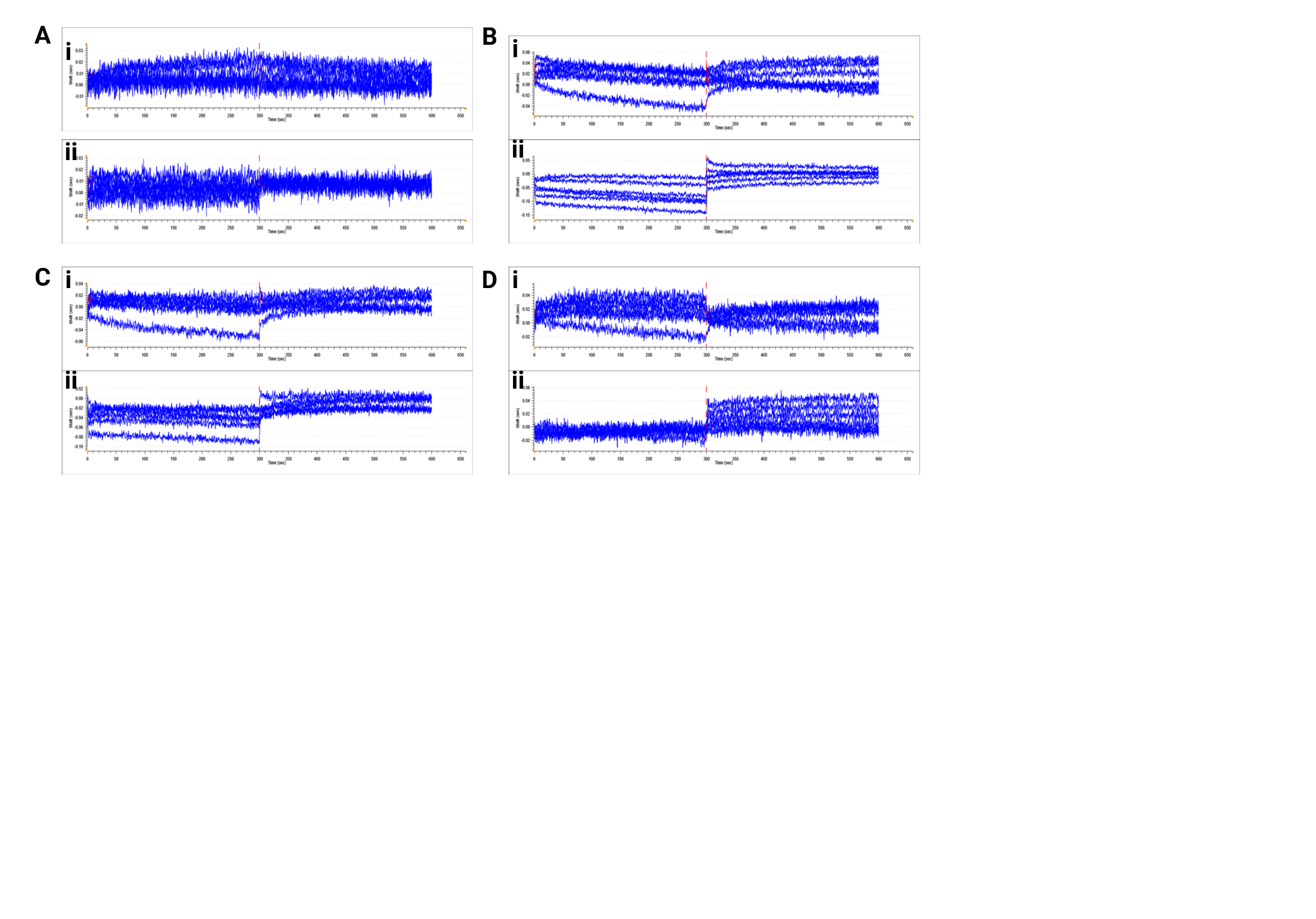

**Supplemental Figure 11. BLI Curves for TEVp-Nb v. dTEVp.** Association and disassociation curves for all TEVp-Nb v. dTEVp BLI assays. (A-D) corresponds to TEVp-Nb4, TEVp-Nb5, TEVp-Nb7, and TEVp-Nb9, respectively. Binding to dTEVp and dTEVp+Sub (incubated before binding assay for 60 minutes) correspond to (i) and (ii), respectively.

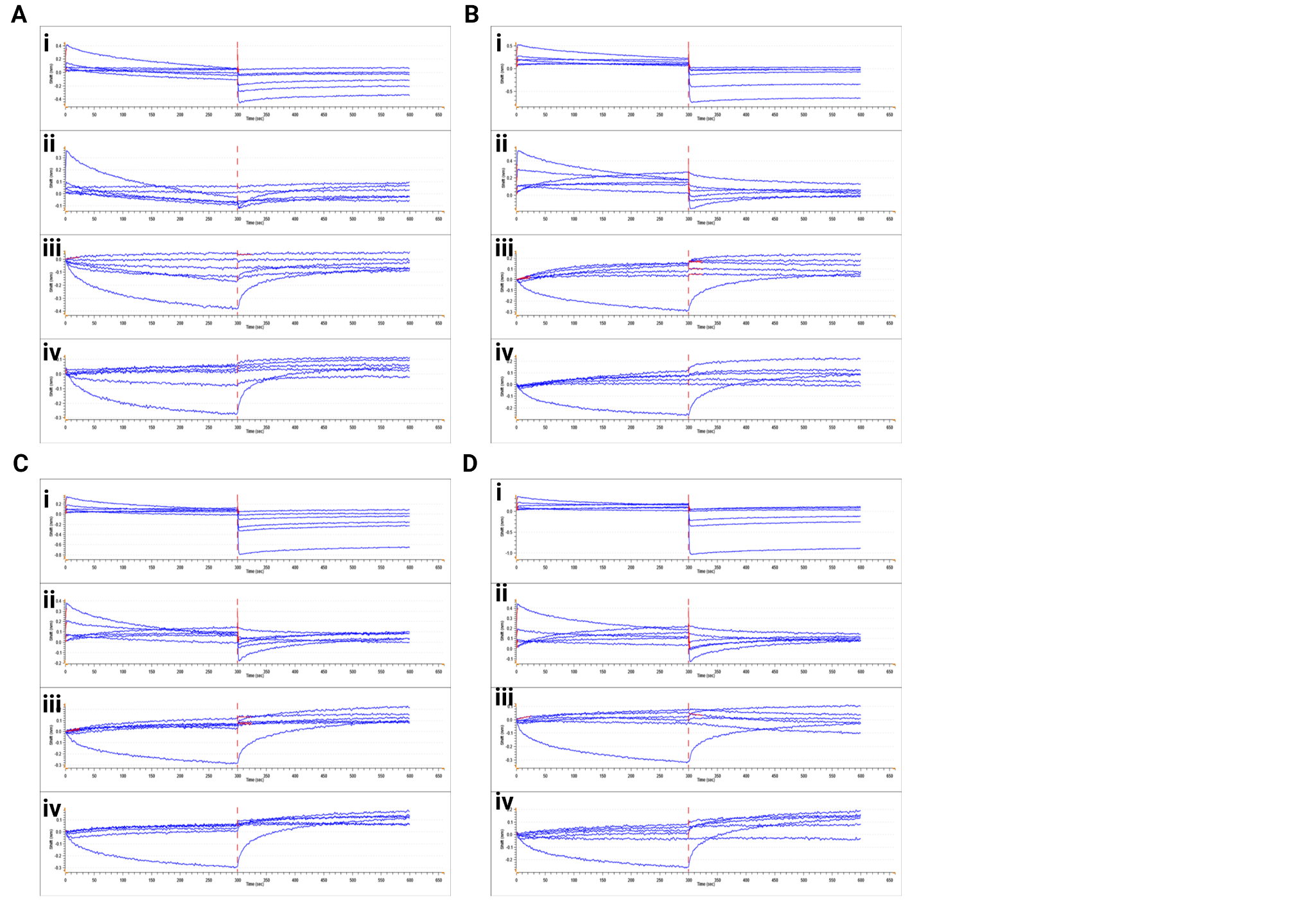

**Supplemental Figure 12. BLI binding curves for TEVp-Nb v. TEVp.** Association and disassociation curves for all TEVp-Nb v. TEVp BLI assays. (A-D) corresponds to TEVp-Nb4, TEVp-Nb5, TEVp-Nb7, and TEVp-Nb9, respectively. Binding assays to TEVp at different concentration titrations correspond to (i-iv).

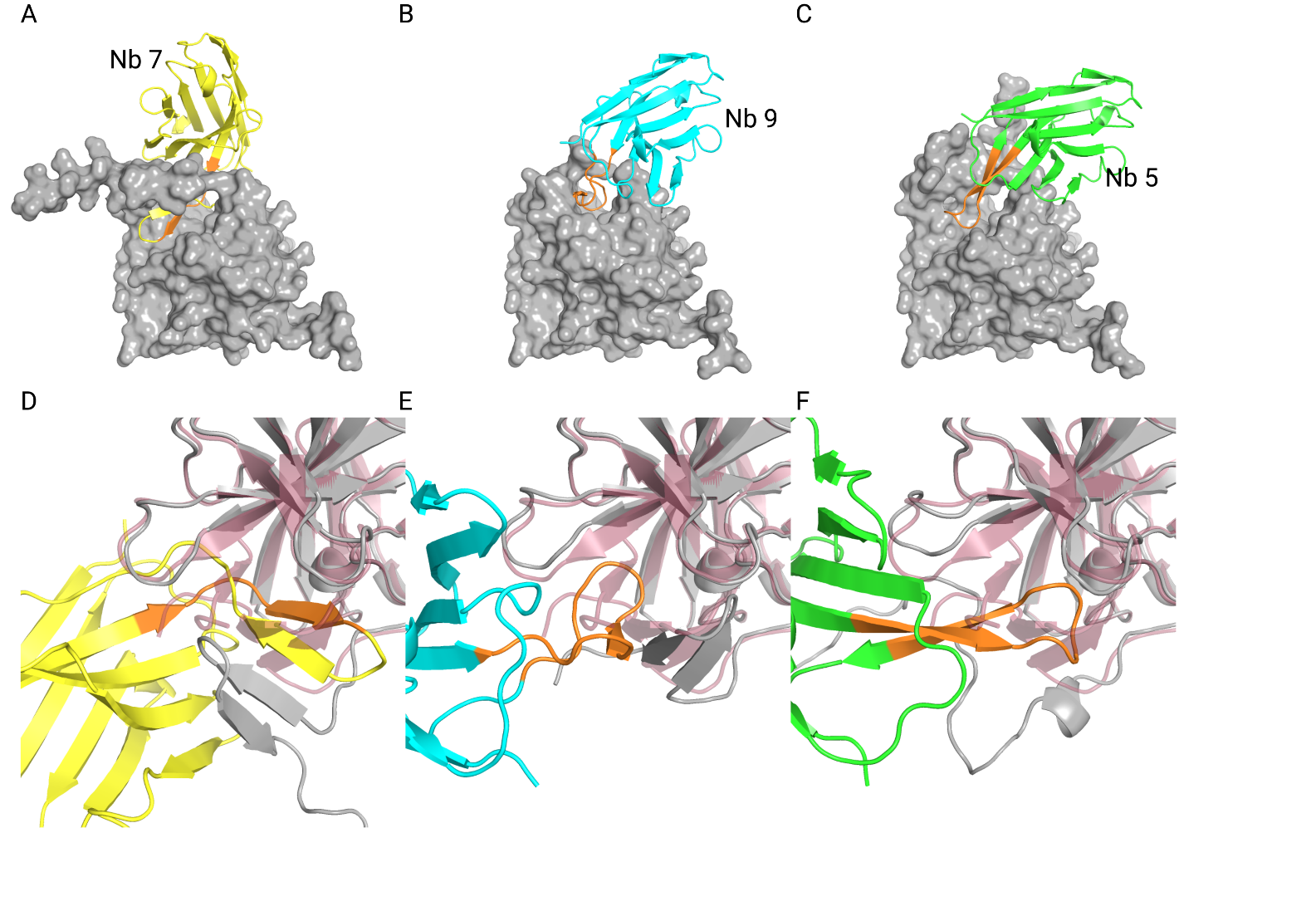

**Supplemental Figure 13. AlphaFold3 models of TEVp-Nbs**. (A–C) AlphaFold3 predictions of TEV-Nb7, TEV-Nb9, and TEV-Nb5 in complex with dTEVp show that their CDR3 loops interact with dTEVp’s active site pocket and disturb its overall structure by forming new interactions. (D–F) Complex structures overlaid with the solved structure of dTEVp (PDB: 1LVB) illustrate the disturbed active site pocket upon binding of the Nb CDR3 loops. AF3 predictions suggest that TEV-Nb7 binds to the active site by forming a parallel beta-sheet interaction.

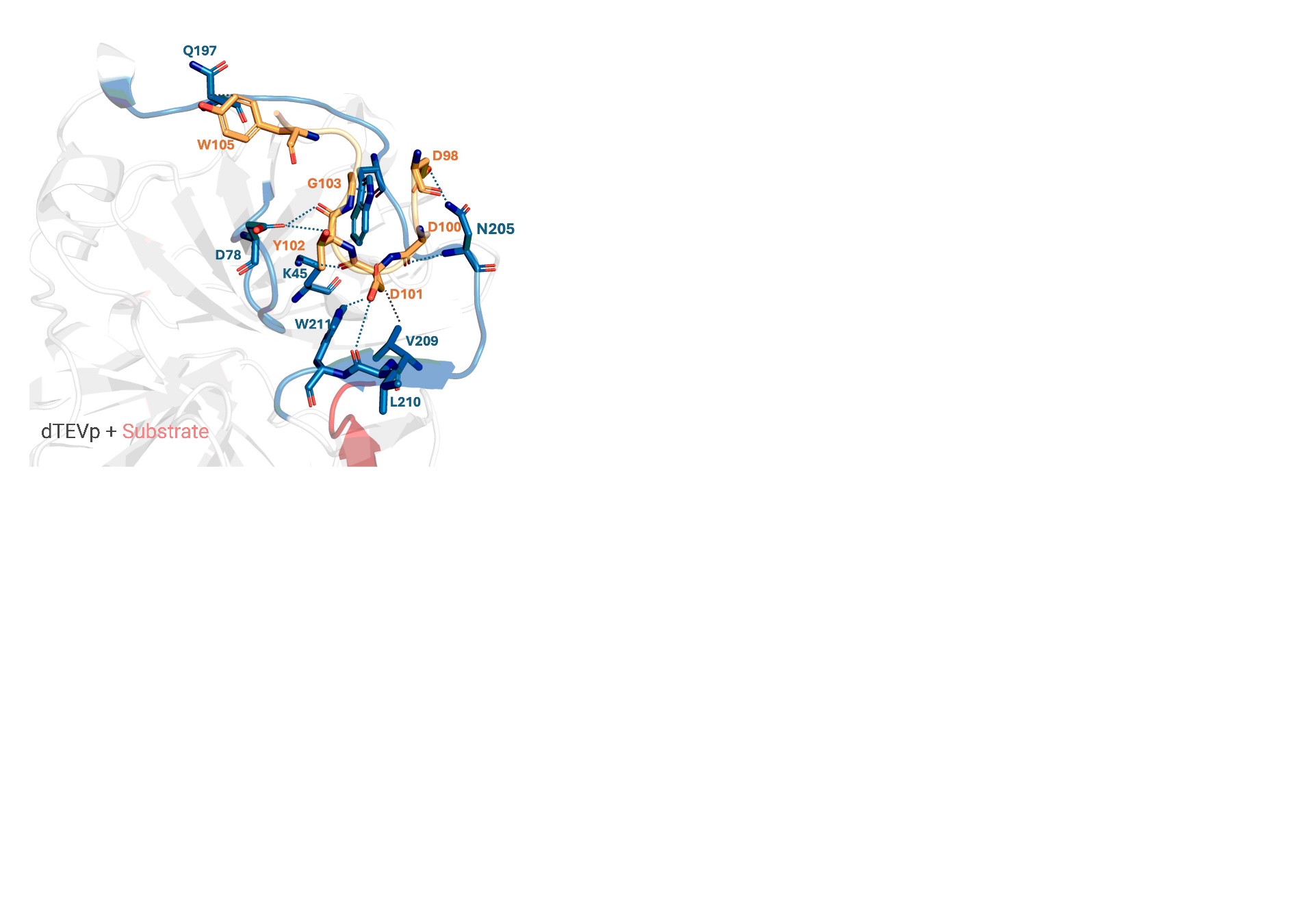

**Supplemental Figure 14. TEVp-Nb4 complex structure predicted by AlphaFold3**. A dTEVp loop, starting at the top of the active site, surrounds the Nb4 CDR3 through a matrix of hydrogen bonds and hydrophobic interactions.

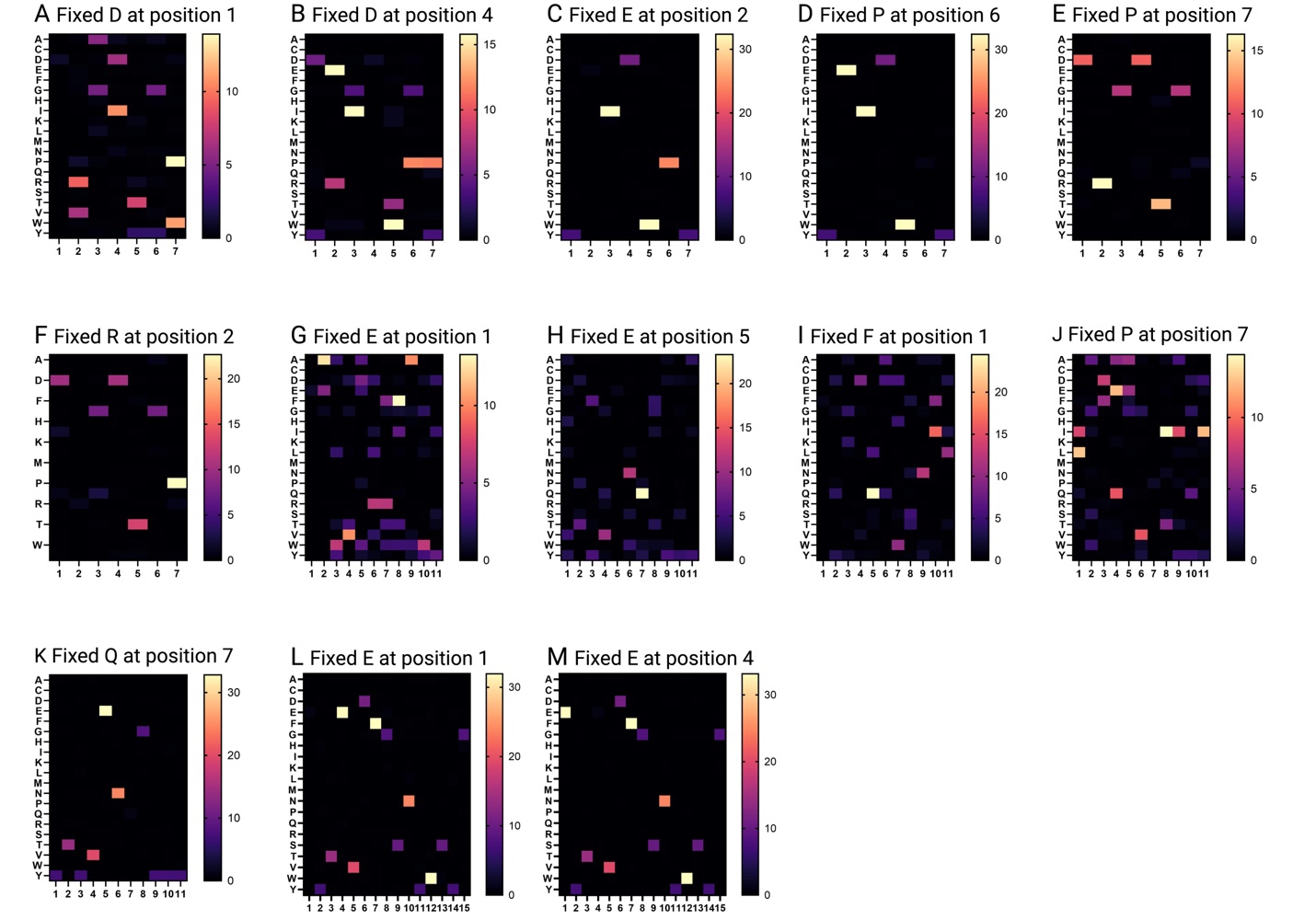

**Supplemental Figure 15. Frequency of amino acids in the CDR3 variable region for different fixed amino acids in the final enrichment population of NbLibrary_NE_TEVp.**

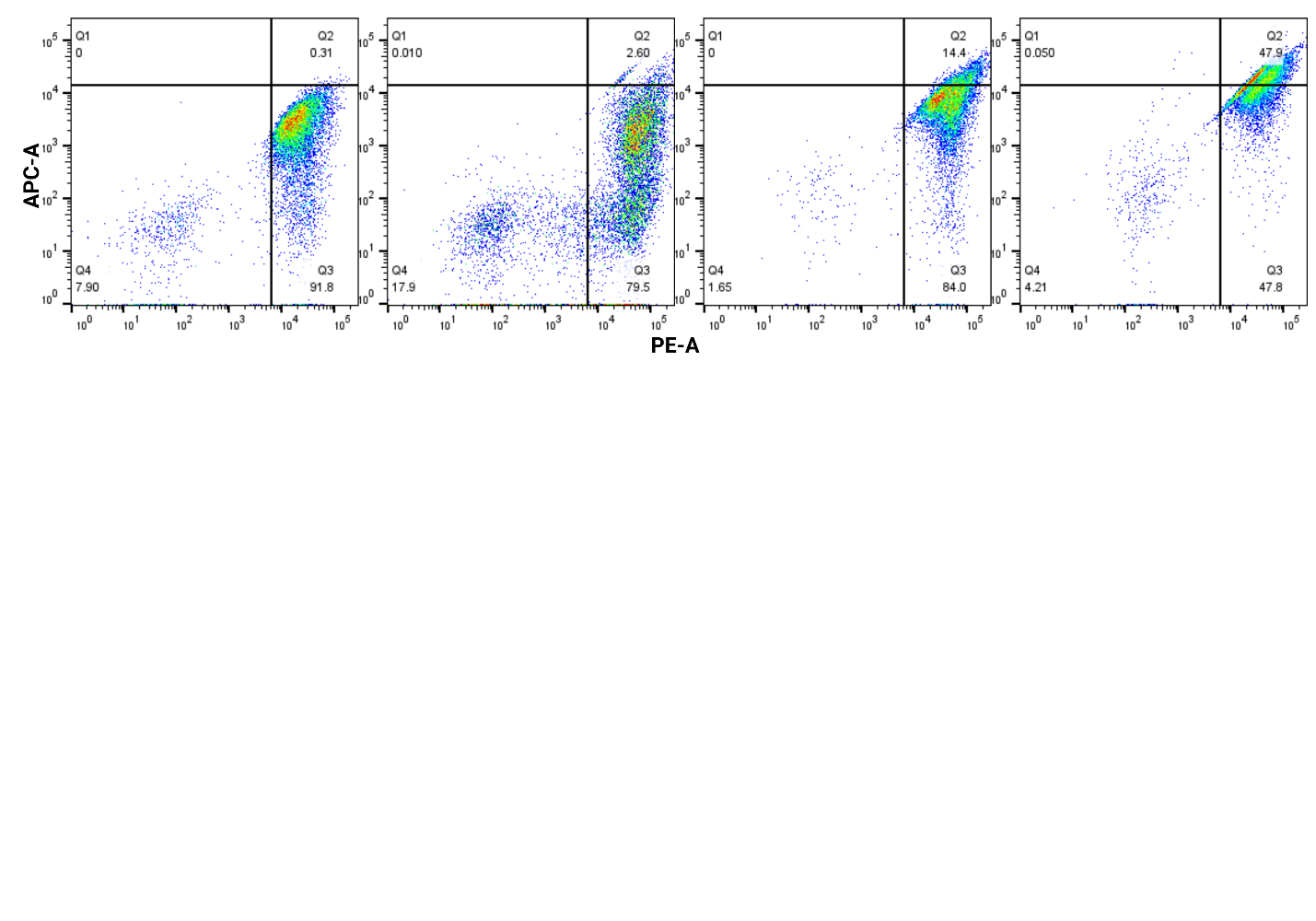

**Supplemental Figure 16. KLK6-NbLibrary-NE FACS enrichment.** Flow plots of each FACS round depicting (from left to right) the initial library, Enrichment 1, Enrichment 2, and Enrichment 3. Dot plots are represented here on a log scale, with PE-A corresponding to anti-FLAG labeling and APC-A corresponding to anti-HA labeling.

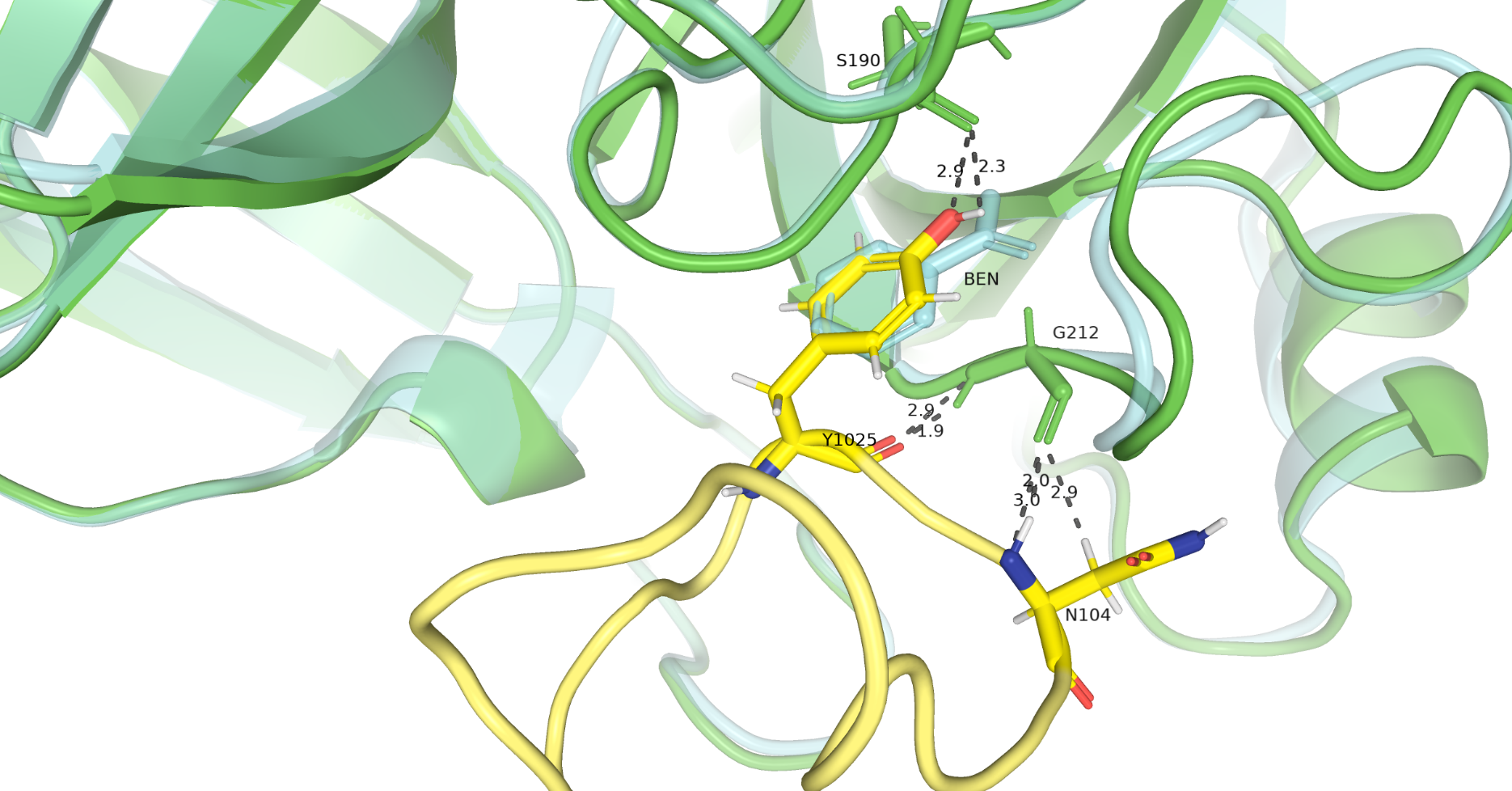

**Supplemental Figure 17. AlphaFold prediction of hK6_Nb1 interaction overlayed on active hK6 with benzamidine inhibitor (PDB: 1L2E).** Overlaying the predicted structure of the hK6_Nb1 complex on the solved structure of hK6 inhibited by benzamidine (in cyan) shows Tyr102 benzene ring (in yellow) placed in a similar location of benzamidine in the S2 pocket.

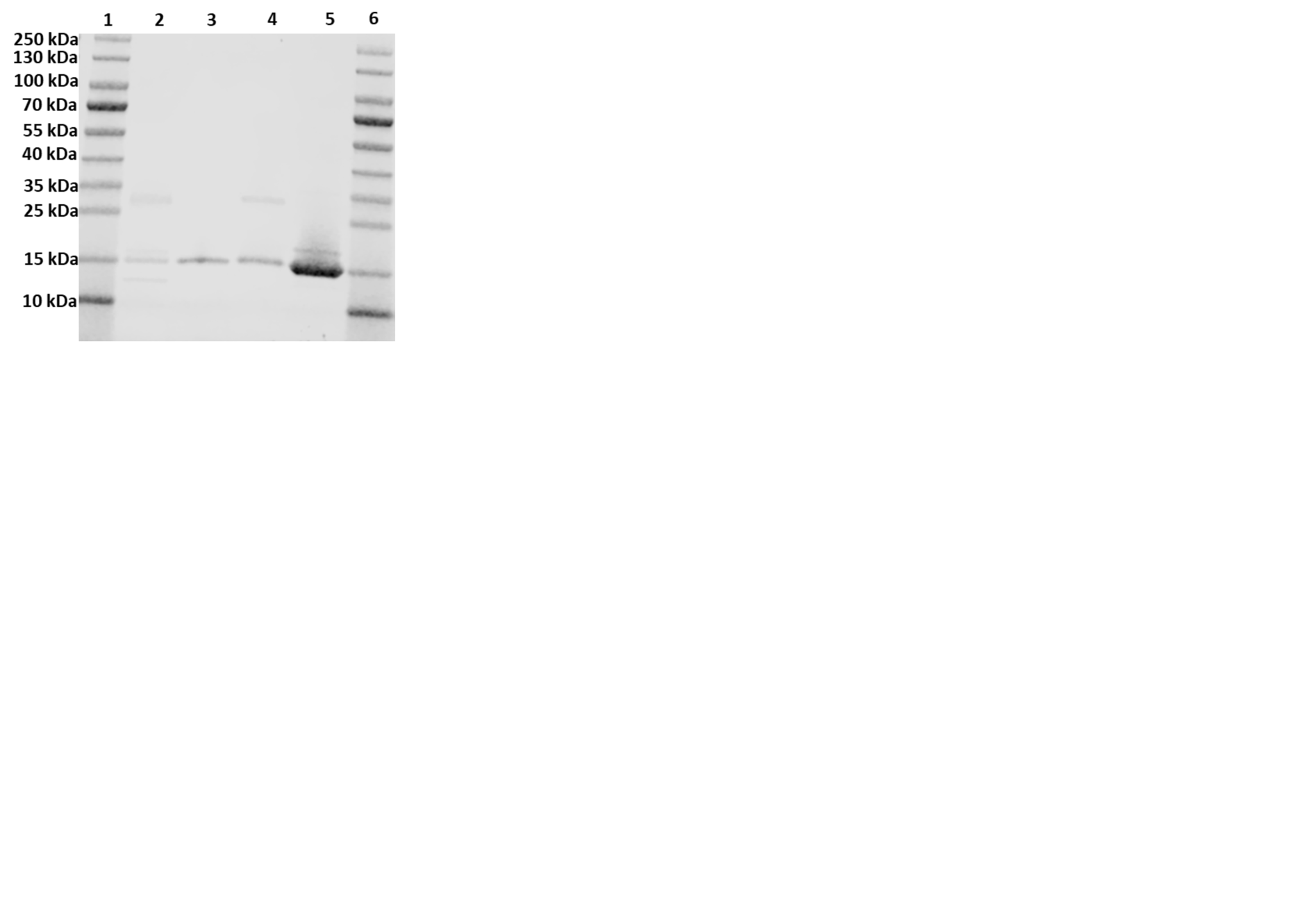

**Supplemental Figure 18. SDS-PAGE gel of purified Nbs from NbLibrary_NE_TEVp**: (Lane 2) Nb4-NE, (Lane 3) Nb5-NE, (Lane 4) Nb7-NE, and (Lane 5) Nb9-NE. Nbs were isolated via Ni-NTA purification.

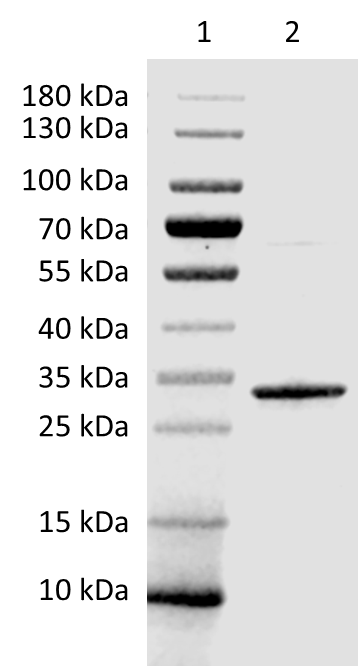

**Supplemental Figure 19. SDS-PAGE of purified TEVp for kinetic assays.**

 
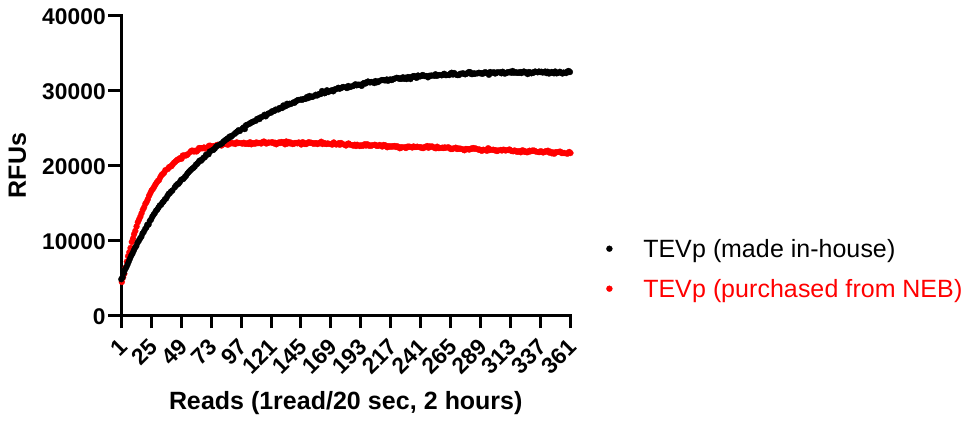

**Supplemental Figure 20. FRET kinetic assay comparison of purified TEVp to purchased TEVp.** Activity assays were conducted at an enzyme concentration of 0.08 uM and a substrate concentration of 20 uM.

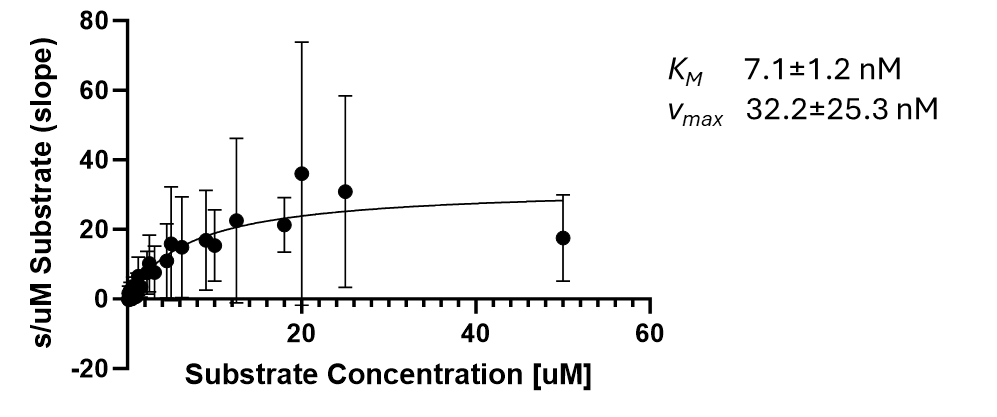

**Supplemental Figure 21. Kinetic characterization of TEVp on FRET peptide.**

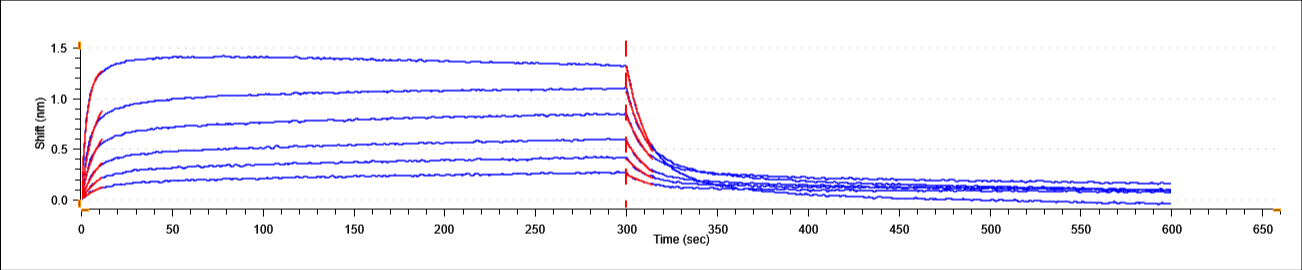

**Supplemental Figure 22. BLI curves for Nb.b201 v. HSA Assay.** Association and disassociation curves for all Nb.b201 v. HSA BLI assay.

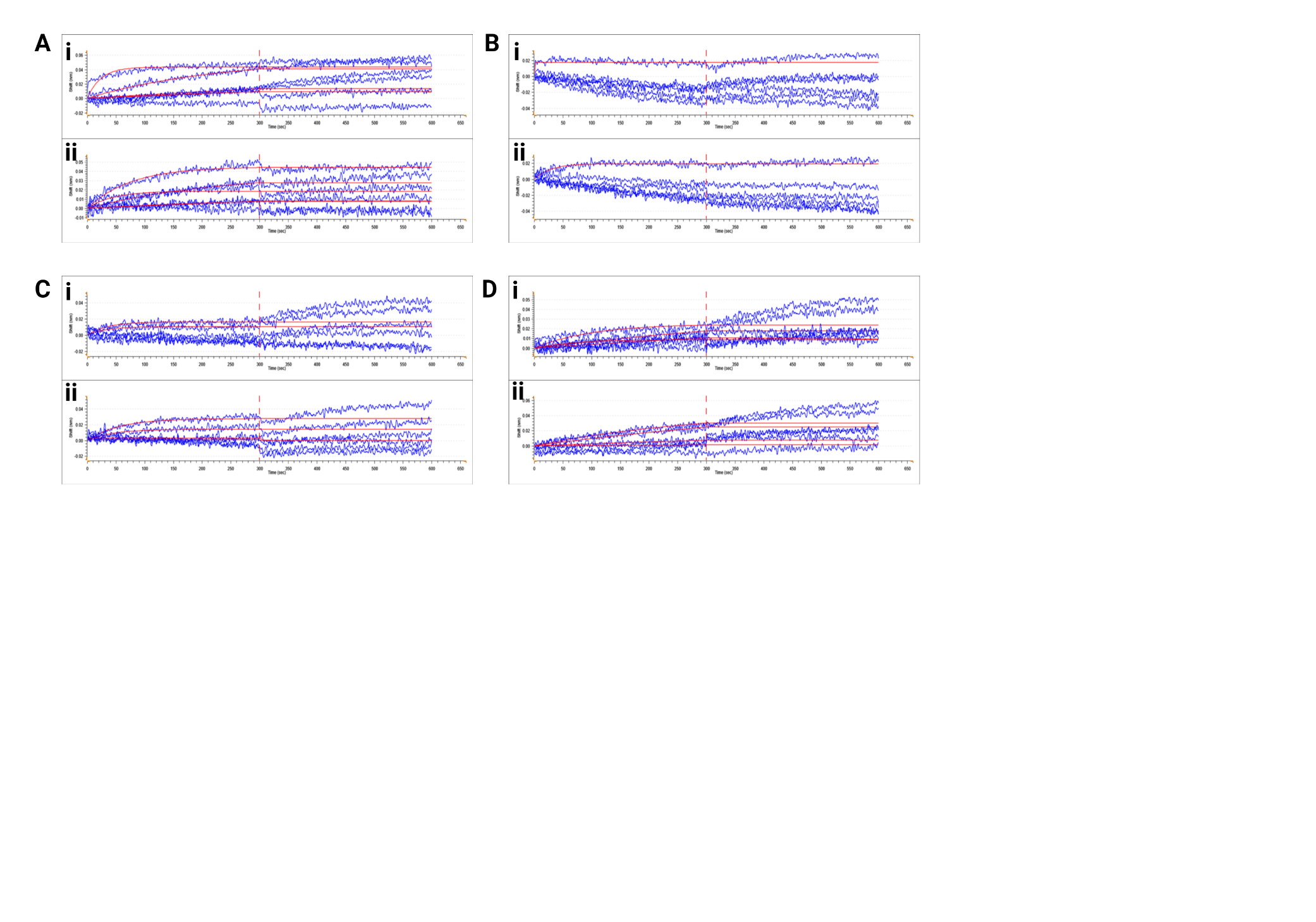

**Supplemental Figure 23. BLI curves for TEVp-Nb v. TVMVp Assays.** Association and disassociation curves for all TEVp-Nb v. TVMVp BLI assays. (A-D) corresponds to TEVp-Nb4, TEVp-Nb5, TEVp-Nb7, and TEVp-Nb9, respectively. Binding assays to TVMVp at different concentration titrations correspond to (i-iv).

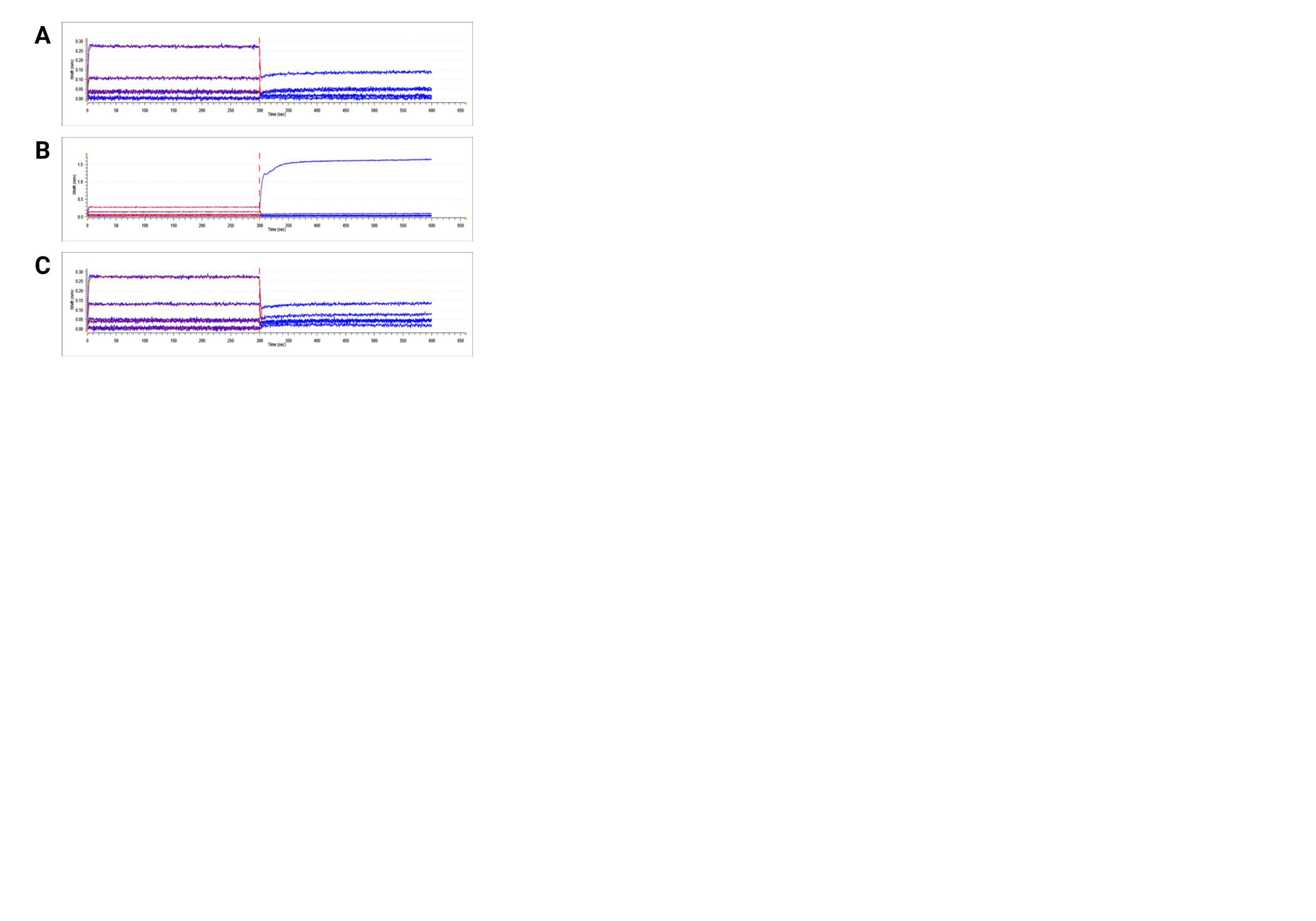

**Supplemental Figure 24. BLI curves for hK6-Nb1 v. KLK6.** Association and disassociation curves for all hK6-Nb v. KLK6 BLI assays. (A-D) corresponds to hK6-Nb1, hK6-Nb3, and hK6-Nb7, respectively.

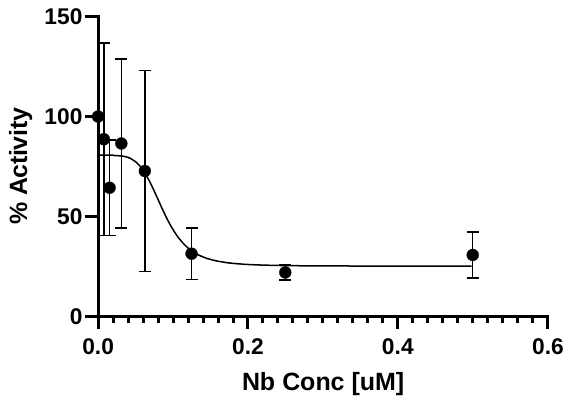

**Supplemental Figure 25. IC50 curve determined via FRET assays for the hK6-Nb1 against KLK6.** All inhibition assays were conducted using a set concentration of KLK6 [0.07 uM] and DABCYL-EDANS substrate [20 uM]. Triplicates were measured, the average IC50 represented on the plot, and the error was propagated by taking the standard deviation of the IC50 calculated for the sample.

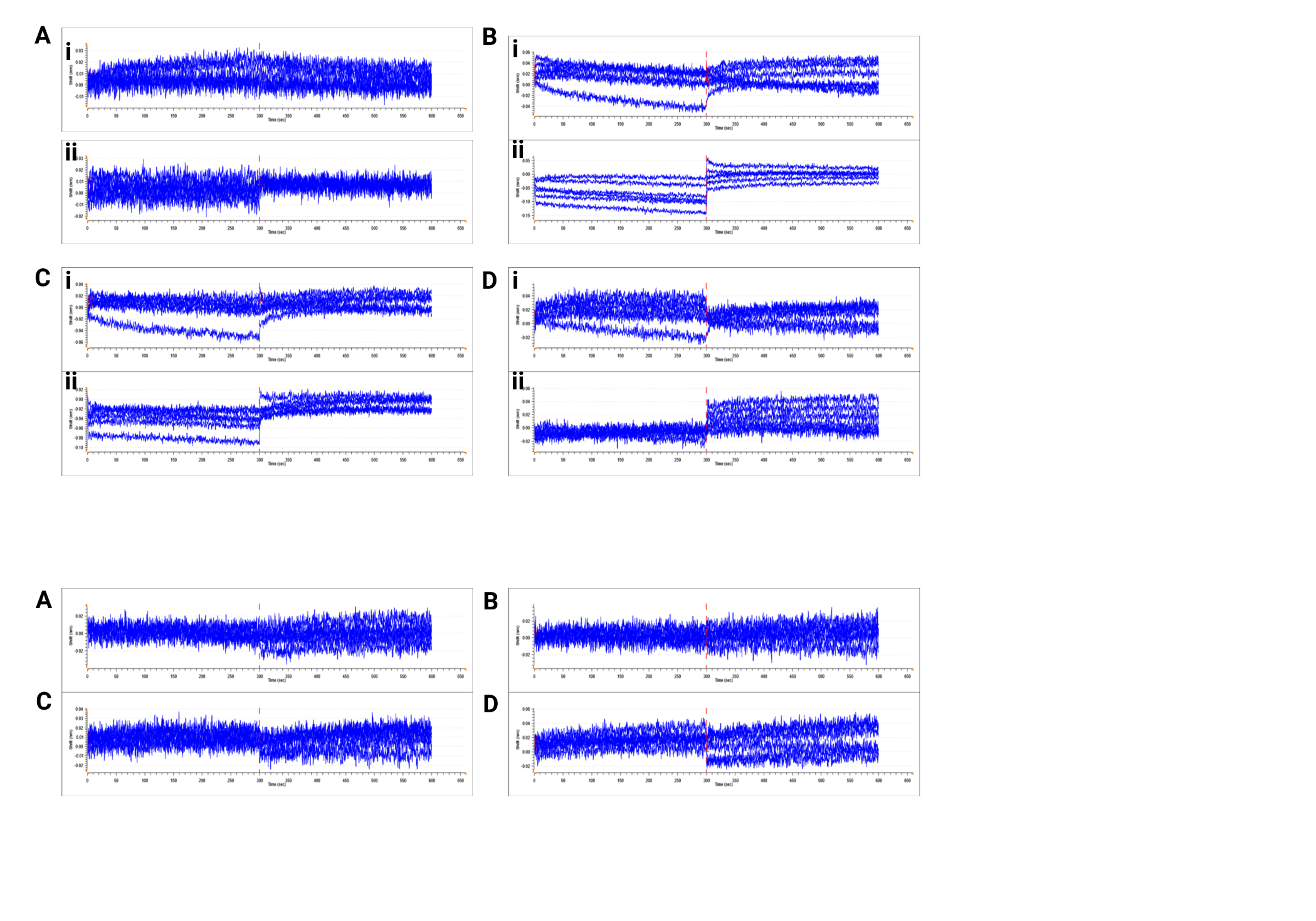

**Supplemental Figure 26. BLI curves for hK6-Nb1 v. KLK Variants.** Association and disassociation curves for hK6-Nb1 v. KLK Variant BLI assays. (A-D) corresponding to KLK2, KLK5, KLK10, and KLK1, respectively.

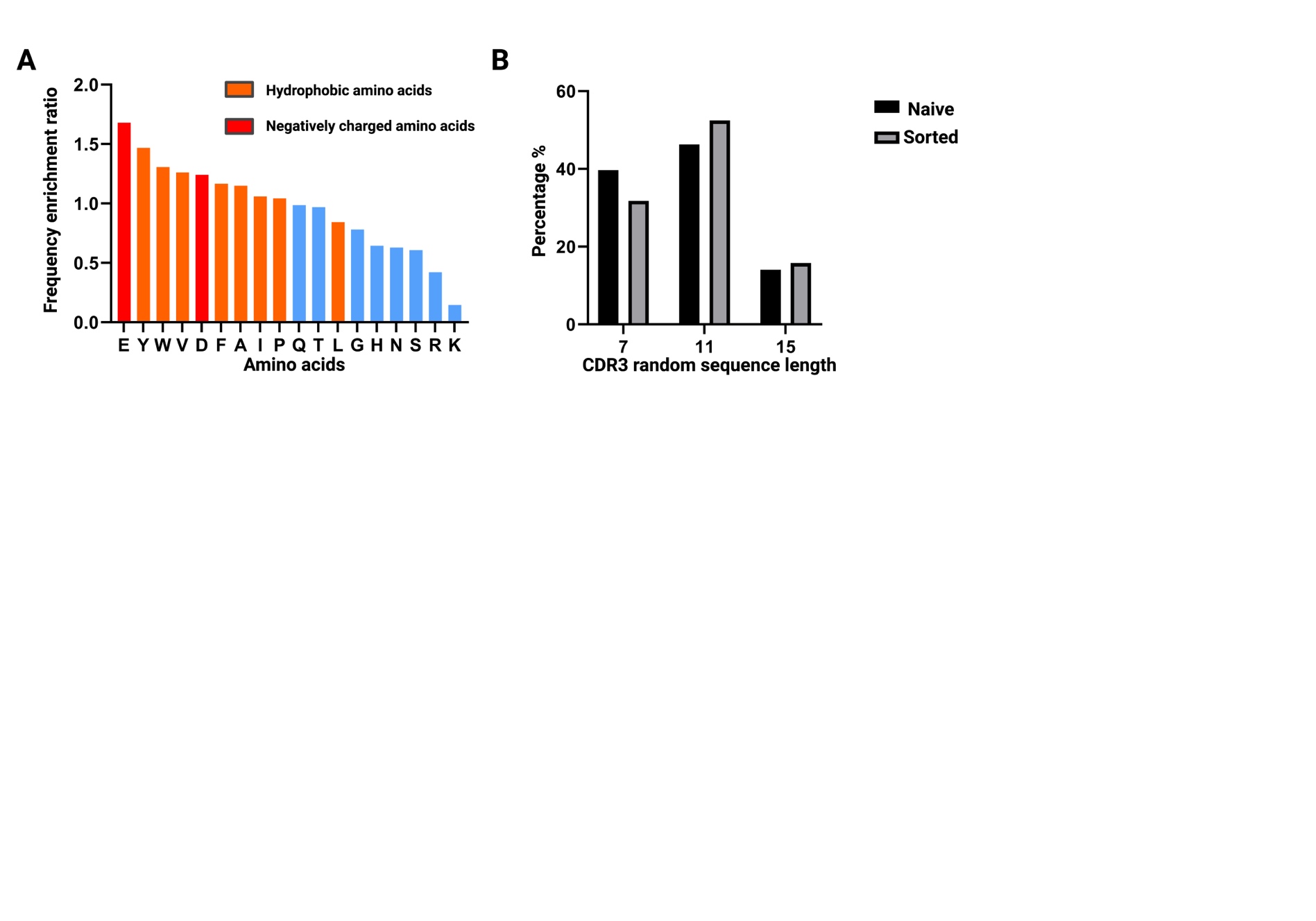

**Supplemental Figure 27. Properties of CDR3 regions in the final enrichment against naïve library.** (A) Frequency of amino acids and their corresponding properties in the CDR3 in the final enrichment population of NbLibrary_NE_TEVp. (B) Length distribution of CDR3 variable region in round 4 vs Naïve.

Supplemental Tables

Supplemental Table 1. Plasmids referenced in the manuscript.

| **Strain or Plasmid** | **Key Characteristics** | **Plasmid Map**  **(if applicable)** |
| --- | --- | --- |
| **Strains** | | |
| LLE | *S. cerevisiae*  MATa AGA1::GAL1-AGA1::URA3 ura3-52 trp1 leu2-delta200::pACT1-LexA-hER-haB112-ADH1t_pTEF1-ble-TEFt his3-delta200 pep4::HIS3 prbd1.6R can1 GAL |  |
| LLE-pY3-LYS-TEVp-FF | Integration Site: LYS2  Yeast Marker: Leu2  Protease Promoter: *pGAL1-10*  Protease: TEVp  Protease ERS: FEHDEL  Substrate Promoter: *pGAL1-10*  Substrate: ENLYFQS  Substrate ERS: FEHDEL |  |
| LLE-pY2-MET-TEVsubstrate-FEH | Integration Site: MET15  Yeast Marker: Leu2  Substrate Promoter: *pGAL1*  Substrate: ENLYFQS  Substrate ERS: FEHDEL |  |
| **Plasmids** | | |
| pN3 | Promoter: *pGAL1*  *E. coli* Marker: AmpR  Yeast Marker: HygroR  Copy Number: Cen6/ARS | <https://benchling.com/s/seq-SFBATiwJUwbjJzeOdINH?m=slm-q7p9tIrfJhJD3K6QqrDD> |
| pY3-LYS-IV | Integration Site: LYS2  Yeast Marker: Leu2  Protease Promoter: *pGAL1-10*  Substrate Promoter: *pGAL1-10* | <https://benchling.com/s/seq-WFK228gqqGFB7y7hjGHe?m=slm-1xOs5ZvtCMjLeJXS8KYl> |
| pN3-Nb14 | Nb Insert: Nb14  Nb Insert ERS: WEHDEL | <https://benchling.com/s/seq-YphVNn5J8iiPtqjBzP2c?m=slm-JUnicPbFpA8SduF15nEK> |
| pN3-Nb14-NE | Nb Insert ERS: NONE |  |
| pN3-5vnv | Nb Insert: 5vnv  Nb Insert ERS: WEHDEL | <https://benchling.com/s/seq-8wY9wFGvlFoqmtvBLX7t?m=slm-uGRNbtxviS6lXqmr7w57> |
| pY3-MMP8-WF | Protease Promoter: *pGAL1-10*  Protease: MMP8p  Protease ERS: WEHDEL  Substrate Promoter: *pGAL1-10*  Substrate: PLGLVA  Substrate ERS: FEHDEL | <https://benchling.com/s/seq-ZaW46SaHH6HTveq6i9Mq?m=slm-aXksuZ9BTknB5wtQm9HX> |
| pY3-NbLibRec | Promoter: *pGAL1*  *E. coli* Marker: AmpR  Yeast Marker: TRP  Copy Number: Cen6/ARS | <https://benchling.com/s/seq-p8oo3ZCT0l1UmuucwoR4?m=slm-3So7SifBZJpazMtZQU9O> |
| pET26b-BsaI |  | <https://benchling.com/s/seq-hefUVTtyLbpSJSYc7kwZ?m=slm-BegZ0dRyaGam8Mm2fie3> |
| pET26b-Nb4-NE | Nb Insert: Nb4-NE | <https://benchling.com/s/seq-7zl4FLWMLIUPMvEc9aTf?m=slm-K22JrYXrkSAnQ71mEOS4> |
| pET26b-Nb5-NE | Nb Insert: Nb5-NE | <https://benchling.com/s/seq-Wu3LgDQDHwfhxGJCDlVo?m=slm-UJptLwFwliojcE3vX2mY> |
| pET26b-Nb7-NE | Nb Insert: Nb7-NE | <https://benchling.com/s/seq-8AR451w0CnrPMzEXzdyu?m=slm-DoOb1D5bZKZr8cYZ6rVY> |
| pET26b-Nb9-NE | Nb Insert: Nb9-NE | <https://benchling.com/s/seq-zpPuiHwL93DXryerlIHL?m=slm-gk2EP3YPBWwDLzEUpwXy> |
| pN4 | Promoter: *pGAL1*  *E. coli* Marker: AmpR  Yeast Marker: TRP  Copy Number: Cen6/ARS | <https://benchling.com/s/seq-TpoHECjJyN5ebG8nfVoH?m=slm-RYtPQP4wU4r9Apk6Sndu> |

Supplemental Table 2. CDR3 enrichment of isolated TEVp-Nbs,

| **Clone** | **CDR3** | **CDR3 Rank** | **Nb Rank** | **Residual Activity** |
| --- | --- | --- | --- | --- |
| TEVp-Nb1 | VFYGDQDDSNILYT | 6 | 5 | 12% |
| TEVp-Nb2 | VVDAYRDSYYTVLL | 42 | 41 | 22% |
| TEVp-Nb3 | AEYTEVDFGSNYWSYGHF | 4 | 2 | 15% |
| TEVp-Nb4 | VDRGDTGPYP | 2 | 3 | 5% |
| TEVp-Nb5 | VEAADYLFGYGYYA | 21 | 25 | 7% |
| TEVp-Nb6 | VIELYAYHDYRYFN | 688 | NF | 21% |
| TEVp-Nb7 | AYTYSQDYWGVLTYVGHG | 18 | 20 | 9% |
| TEVp-Nb8 | AYTYVENQGYYYFA | 3 | 1 | 15% |
| TEVp-Nb9 | VAPFYEWYFDYYYI | 14 | 13 | 10% |
| TEVp-Nb10 | AEYTEVDFGSNYWSYGHF | 4 | 2 | 13% |
| TEVp-Nb11 | Sequencing failed |  |  | 6% |
| TEVp-Nb12 | AEYTEVDFGSNYWSYGHF | 4 | 2 | 14% |
| TEVp-Nb13 | Sequencing failed |  |  | 11% |
| TEVp-Nb14 | VYDYTWTYSWVYLYS | 60 | 79 | 20% |
| TEVp-Nb15 | VFYGDQDDSNILYT | 6 | 5 | 13% |
| TEVp-Nb16 | VEAYVDRRFAYYYN | 10 | 7 | 16% |
| TEVp-Nb17 | AYTYSQDYWGVLTYVGHG | 18 | 20 | 17% |
| TEVp-Nb18 | AFGAYSAHNAAIFF | 32 | 36 | 19% |
| TEVp-Nb19 | AHSYRELYYYGDYP | 30 | 34 | 17% |
| TEVp-Nb20 | AHSYRELYYYGDYP | 30 | 34 | 16% |
| TEVp-Nb21 | AYTYVENQGYYYFA | 3 | 107 | 13% |
| TEVp-Nb22 | VFYGDQDDSNILYT | 6 | 5 | 14% |
| TEVp-Nb23 | AIADAGGPIIGDFY | 19 | 32 | 17% |
| TEVp-Nb24 | AYTYVENQGYYYFA | 3 | 1 | 20% |

Supplemental Table 3. Read counts of the top fifty TEVp-Nb sequences in ENR4.

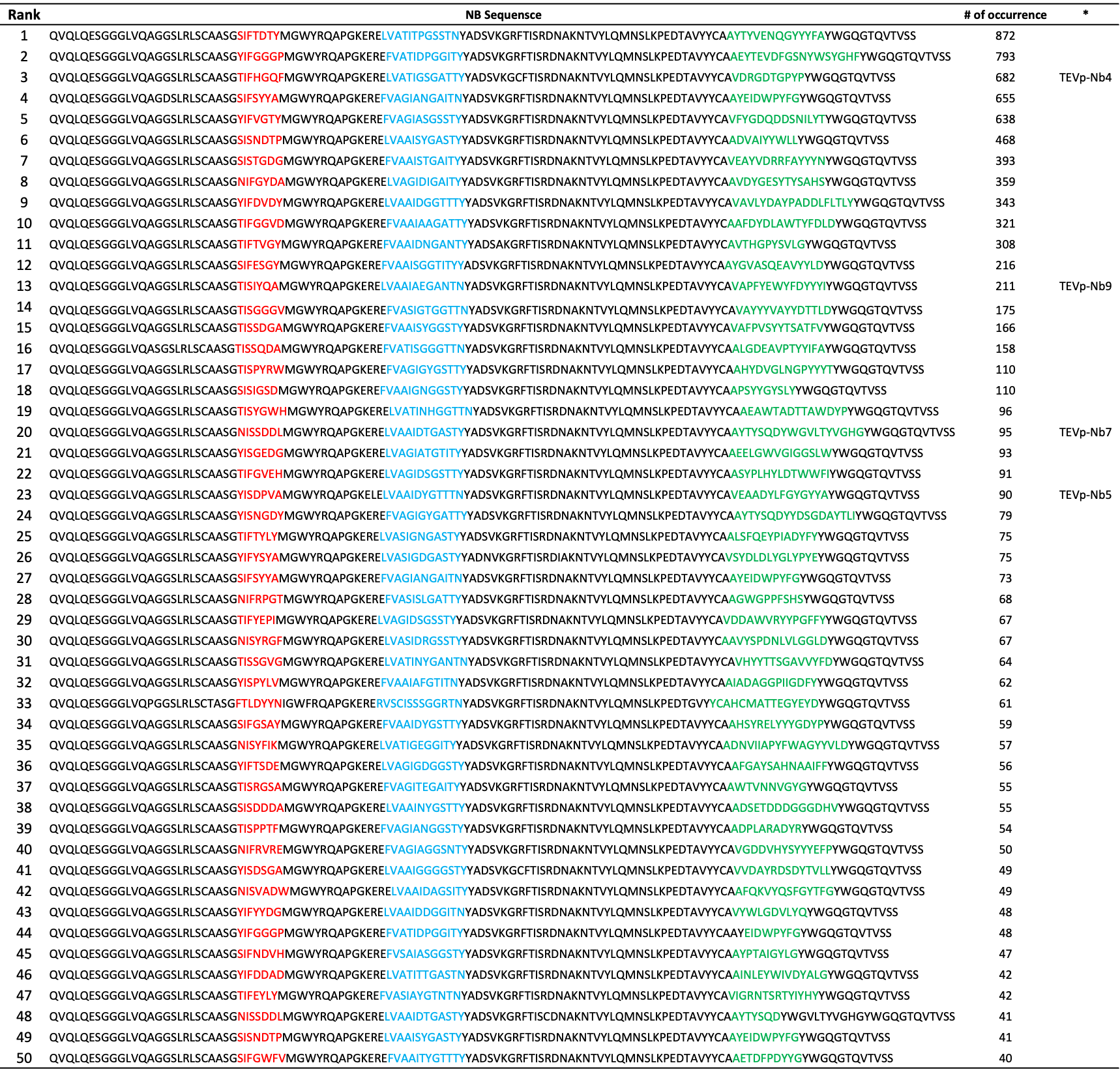

Supplemental Table 4. Binding characterization of the isolated TEVp-Nbs.

| **Nb** | ***K_D_*** **[nM]**  **TEVp** |
| --- | --- |
| **TEVp-Nb4** | 348 ± 14.5 |
| **TEVp-Nb9** | 782 ± 37.2 |
| **TEVp-Nb5** | 411.7 ± 3.9 |
| **TEVp-Nb7** | 2980 ± 106 |

Supplemental Table 5. CDR3 sequences in round 4 containing ENQ motif.

| CDR3 | # of Reads |
| --- | --- |
| AYTYVENQGYYYFA | 1899 |
| AYTYVENQGYYYLA | 2 |
| AYTYVENQGYYYFV | 2 |
| AYTYVENQDYYYFA | 2 |
| AYTYVENQGYYEFP | 2 |
| VYTYVENQGYYYFA | 1 |
| AYSYVENQGYYYFA | 1 |
| AYTNVENQGYYYFA | 1 |
| AYAYVENQGYYYFA | 1 |
| SYTYVENQGYYYFA | 1 |
| AYTYVENQVYYYFD | 1 |
| AYTYVENQGYFDYYYI | 1 |
| AYTYVENQGYHYFA | 1 |
| AYTYVENQGYSYFA | 1 |
| AYTYVENQGYYA | 1 |
| AYTYVENQGYYYFD | 1 |

Supplemental Table 6. Characterization of the isolated hK6-Nbs against KLK6.

| **Nb** | **Sequence** | ***IC50* [nM]** | ***K_D_* [nM]** |
| --- | --- | --- | --- |
| hK6-Nb3 | QVQLQESGGGLVQAGGSLRLSCAASGTISTYGGMGWYRQAPGKERELVASIGGGSSTNYADSVKGRFTISRDNAKNTVYLQMNSLKPEDTAVYYCAAVGATYIIHRYWGQGTQVTVSS | N/D | 811 ± 21.9 |
| hK6-Nb7 | QVQLQESGGGLVQAGGSLRLSCAASGNIFYGFPMGWYRQAPGKERELVATINDGGSTNYADSVKGRFTISRDNAKNTVYLQMNSLKPEDTAVYYCAVGSHTASGDVGYYGSDFAYWGQGTQVTVSS | N/D | 207 ± 10.5 |

Supplemental Table 7. Primer sequences for the assemblies referenced in the manuscript.

| **Primer Name** | **Primer Sequence** |
| --- | --- |
| TWIST-NbLib-Nested-F | ACCATGTACCATGTAAACCTGTTATAAC |
| TWIST-NbLib-Nested-R | TGGATCTTAAAAACCAAACGTGG |
| Making-NbLib-F | ATGCAGTTACTTCGCTG |
| Making NbLib-R (contains FEHDEL ERS) | AGTCGATTTTGTTACATCTACAC |
| Making NbLib-R (without ERS) | TATCAGATCTCGAGCTATTACTAGCCGCGGTACCAAGCTTAGCTGCTCACGGTCACCTGG |
| BsaI-NbLib-NE-PCR-F | TCCTTAGAGGTCTCGGGCAGCGGCAGCCAGGTGCAGCTGCAGGAAAGC |
| BsaI-NbLib-NE-PCR-R | ACATACAGGGTCTCGATGGGCGGCAGCGCTGCTCACGGTCACCTGGGTG |

Supplemental Table 8. Results of control BLI assay, exploring the influence of trace amounts of glycerol on resulting disassociation constant.

| **Nb** | ***K_D_*** **[nM]**  **TEVp** |
| --- | --- |
| TEVp-Nb5 (frozen stock, contains a trace of glycerol) | 103 ± 7.85 |
| TEVp-Nb5 (buffer swap to remove glycerol) | 117 ± 7.76 |

Methods and Materials

*System Construction*

Inhibitor plasmid

A receiver plasmid, pN3 **(Supplemental Table 1)**, was designed to contain a Cen6/ARS copy number gene, the hygromycin resistance (HygR) selection marker, and unique cute sites (*HindIII* and *XhoI*) for ease of cloning. Using the unique cut sites, pN3 was linearized and assembled with the Nb-containing insert (with homology regions to the 5’ and 3’ ends of the linear plasmid) using Gibson Assembly. The pN3-Nb assemblies **(Supplemental Table 1)** were transformed into competent *Escherichia coli* (*E. coli*) (NEB#C2984H), purified (QIAGEN #27106), and sent for sequencing to confirm correct assembly (Plasmidsaurus, standard plasmid).

*Proof-of-Concept MMP8 Nb Sort*

In preparation, Nb14 (MMP8-inhibiting nanobody) ^1^ and 5vnv (non-binding, non-inhibitory nanobody) ^2^ sequences were cloned into a receiver plasmid, pN3**,** as mentioned above. MMP8 and its canonical substrate were cloned into a pY2 receiver plasmid, with the protease under ꞵ-estradiol (ꞵE) induction and the substrate under galactose induction. Once the plasmids were sequenced to confirm correct assembly (Plasmidsaurus, standard plasmid), a double-plasmid transformation was conducted into an engineered *S. cerevisiae* strain **(Supplemental Table 1)**, following the protocol provided in the Frozen-EZ Yeast Transformation II Kit (ZymoResearch #T2001). The cells were plated on minimal media (YNB CAA) containing 2% glucose (GLU) and Hygromycin (HYGRO, 200 ug/uL); labeled according to the plasmids transformed into them, MMP8-Nb14 and MMP8-5vnv. Once the plates grew for 2-4 days at 30°C, mature colonies were picked and grown in 1 mL YNB CAA 2% GLU HYGRO at 30°C for 16-24 hours.

Once saturation was reached, the optical density (OD_600_) was taken to determine the approximate amount of culture required for a starting outgrow to have an OD_600_ of 1 in 1 mL media. The calculated culture was inoculated in 1 mL YNB CAA 2% GLU HYGRO and grown to an OD_600_ between 2-4 (30°C, 250 RPM) for approximately 4-6 hours. The OD_600_ of the outgrow was determined, and the amount of culture was calculated to have a starting induction OD_600_ of 0.5 in 1 mL media. Required culture volumes were washed in YNB CAA 2% Galactose (GAL) HYGRO media (to remove residual GLU), and then the cell pellets were resuspended in 1 mL YNB CAA 2% GAL HYGRO and 2 uM ꞵE. Induction cultures were grown for 12-16 hours (30°C, 250 RPM), after which the OD_600_ of each culture was taken. From the OD_600_, the amount of culture was calculated to stain 5 million cells with fluorescently labeled antibodies, as well as the volume of each culture (MMP8-Nb14 and MMP8-5vnv) would need to be combined to have a 1:1000 mixture Nb14: 5vnv (totaling to 5 million cells). The three samples, MMP8-Nb14, MMP8-5vnv, and 1:1000, were washed with PBS 0.5% BSA (0.5% BSA, Goldbio 9048-46-8) and then stained with anti-FLAG PE (Biolegend cat# 637309) and anti-HA Alexa 647 (Biolegend cat# 682404) at RT for 90 minutes. Samples were washed with PBS 0.5% BSA and resuspended for cytometric and FACS (BDFACS Melody, BD Biosciences).

All samples were analyzed using the PE-A (correlating with high anti-FLAG PE signal) and APC-A (correlating with high anti-HA Alexa 647) channels of the BDFACS Melody. To determine the sorting gate strategy, control samples (MMP8-Nb14, MMP8-5vnv) were assayed independently, and a gate was drawn to capture 1% of the cells with the non-inhibitory phenotype, corresponding with 5vnv expression **(Supplemental Figure 4)**. This gating strategy was selected to ensure that an abundance of the inhibitory phenotype, our positive control (MMP8-Nb14), was present in the gate, alluding to the notion that when the mock “library” is assayed, cells with the inhibitor present would reside in this gate. The 1:1000 mixed population sample was assayed, and cells falling in the sorting gate were captured, enriched, and assayed for inhibition enrichment. The cell preparation and cytometric sorts were repeated for two rounds, resulting in almost complete retention of the HA tag (correlating to an inhibited proteases) fluorescent epitope (anti-HA Alexa 647). Cells from the enriched population were cultured, and the Nb-containing plasmids were purified (QIAGEN #27106) and sent for sequencing to confirm Nb identity (Sanger Sequencing, Genewiz, Azenta Life Sciences).

*Library Construction*

DNA preparation

For the first protein scaffold to be screened in HARP, Nbs were selected based on their small size and the ability to purify them relatively easily from *E. coli*. Drawing inspiration from *McMahon* *et al.* ^2^, a starting DNA template was designed with degenerate amino acids within the complementary determining regions (CDRs) of a modified 5vnv Nb sequence **(Figure 2A)**. The Nb framework, along with additional assembly elements (i.e., homology regions, ER retention signal (ERS), identifiers) **(Figure 2B)**, were constructed and synthesized as a double-stranded DNA (dsDNA) pool with Twist Biosciences (variant library synthesis). A nested PCR was conducted to remove the identifier sequences and assemble the library into yeast. A secondary PCR (non-nested) was performed to amplify the library to a final mass of ~15 ug. An additional PCR was required to create the dsDNA required for NbLibrary-NE-TEVp (removal of the FEHDEL ERS). **Supplemental Table 7** contains primer sequences for the reactions mentioned.

A modified Nb receiver plasmid was required to mitigate antibiotic selection for large Nb library construction. This receiver plasmid was designed similarly to the previously reported protease-substrate plasmid, pY3^3^. Following the *pGAL1* promoter, a multiple cloning site (MSC) and a linker (for additional homology) were added to allow for the insertion of desired genes using restriction enzyme digestion and subsequent ligation. The resulting plasmid, pY3-NbLibRec **(Supplemental Table 1)**, was transformed into competent *Escherichia coli* (*E. coli*) (NEB#C2984H), purified (QIAGEN #27106), and sent for sequencing to confirm correct assembly (Plasmidsaurus, standard plasmid). pY3-NbLibRec was linearized through a large-scale *Afl-II* and *Sal-I* restriction enzyme digestion to expose the homology arms and produce the necessary mass of linearized backbone (~15 ug).

Protease yeast strain

Following the integration strategy established previously^3^, TEVp, a gift from David Waugh (Addgene plasmid #8827; http://n2t.net/addgene:8827; RRID: Addgene_8827), and its canonical substrate (ENLYFQS) were cloned into pY3-LYS-IV plasmid **(Supplemental Table 1)**. The plasmid was linearized with *NotI-HF* and was integrated into an engineered *S. cerevisiae* strain **(Supplemental Table 1)** using the lithium acetate (LiAc) transformation protocol established by Gietz and Schiestl^4^. The cells were plated on selection media, SC-LEU, and grow at 30°C for 2-4 days. Mature colonies were picked and grown in 1 mL selection media at 30°C for 16-24 hours until saturated.

Homologous-directed recombination (HDR) assembly for the Nb library in yeast

LLE-pY3-LYS-TEVp-FF yeast cells **(Supplemental Table 1)** were prepared for electroporation using the protocol reported by *Loock* *et al.* ^5^. Once prepared, the electro-competent pY3-LYS-TEV-LLE cells (600 uL/aliquot) were kept on ice, along with 1 2-mm electroporation cuvette, for five minutes before 10 ug of both prepared library and linearized backbone were added to the cells. Following a second 10-minute incubator, the cell-DNA mixture was transferred to the cold cuvette and electroporated using the Sc.2 setting on the Bio-Rad MicroPulser (#1652100). Freshly electroporated cells were transferred to a sterile flask containing 12 mL YPD: Sorbitol and incubated for 1 hour (30°C, 250 RPM). The cells were spun down (4°C, 5 minutes, 500 RCF) and resuspended in a 2-L sterile flask containing 375 mL YNB 2% GLU. Library culture was incubated for 2-3 days (30°C, 250 RPM) and passaged after saturation was reached to an OD_600_ of 0.2 in 500 mL YNB 2% GLU to give a culture 10-20 times larger than the starting library size. Actual library size was determined by serial dilution plating of starting library culture on selection media (YNB CAA 2% GLU) and incubated in a stationary incubator at 30°C for 2 days. The saturated culture, called NbLibrary-NE-TEVp henceforth, was stored at 4°C until the screening.

*Isolating and Purifying Inhibitors using Fluorescence-Activated Cell Sorting (FACS)*

Cell Preparation

**NbLibrary-NE-TEVp**

Saturated NbLibrary-NE-TEVp culture (see Methods: Homologous-Directed Recombination (HDR) Assembly of Nb Library in Yeast) was re-inoculated in a sterile 250-mL flask to an OD_600_ of 0.5 in 50 mL selection media (YNB CAA 2% GLU) and grown at 30°C (250 rpm) until OD_600_ between 2-4 was reached (approximately 5 hours). Once the desired OD_600_ was achieved, the culture was re-inoculated in a sterile 250-mL flask to an OD_600_ of 0.5 in 50 mL induction media (YNB CAA 2% GAL). The induced culture was incubated at 30°C (250 rpm) for 12-15 hours. Post-induction OD_600_ was taken, and the volume of culture required to contain approximately 10x NbLibrary-NE-TEVp size was calculated and added to 3 mL PBS 0.5% BSA (0.5% BSA, Goldbio 9048-46-8). Cells were washed and stained with fluorescently tagged antibodies (anti-FLAG PE, Biolegend, cat# 637309, anti-HA Alexa 647, Biolegend, cat #682404) at a concentration of 10^5^ cells/uL. Staining was done for 90 minutes in the dark at RT. The excess antibody was removed with a second PBS 0.5% BSA wash and resuspended for FACS (BDFACS Melody, BD Biosciences).

**Control Cells**

OD_600_ of the saturated cultures (inoculation from colony, 1 mL YPD, 30°C, 250 RPM) of LLE-pY2-MET-TEVsubstrate-FEH (positive control) and LLE-pY3-LYS-TEVp-FF (negative control) were taken, and volume calculated to re-inoculate to an OD_600_ of 0.5 in 2mL selection media (SC -URA 2% GLU) were determined. Calculated volumes of each were added to the media and inoculated at 30°C (250 rpm) for approximately 5 hours until an OD_600_ was reached between 2 and 4. The OD_600_ of the outgrows was taken, and the volume of cultures was calculated to have an induction starting OD_600_ of 0.5 in 2 mL induction media (SC -URA 2% GAL). Calculated volumes of each were added to the media and inoculated at 30°C (250 rpm) for 12-15 hours. Post-induction OD_600_ was taken, and the volume of culture required to contain approximately 2 million cells was calculated. The cells were washed in 1 mL PBA 0.5% BSA and resuspended in 50 uL staining mix containing PBS 0.5% BSA and fluorescently tagged antibodies (anti-FLAG PE, Biolegend – 0.5 uL/2 million cells, cat# 637309, anti-HA 374 Alexa 647 – 1 uL/2 million cells, Biolegend, cat #682404). Staining was done for 90 minutes in the dark at RT. The excess antibody was removed with a second PBS 0.5% BSA wash and resuspended for FACS (BDFACS Melody, BD Biosciences).

Cytometric Activity Check of Controls

To analyze the control populations using flow cytometry, fluorochrome channels on the BDFACS Melody were selected based on the excitation/emission wavelengths of the antibodies used in the cell staining. High display levels of our FLAG epitope tag (stained with anti-FLAG PE) were analyzed using PE-A, and high display levels of our HA epitope tag (stained with anti-HA Alexa 647) were analyzed using APC-A. Yeast cell populations were captured via a gate at ≥10^4^ FSC-A and ≥10^4^ SSC-A **(Supplemental Figure 8A)** and were assayed for fluorescence on a PE-A (x) v APC-A (y) dot plot. Displaying cells were captured using a gate at ≥10^4^ PE-A and all ranges of APC-A **(Supplemental Figure 8C)**. FCS Files were obtained and further analyzed using FlowJo (Version 10.9). These populations aim to serve as the phenotypic controls for the FACS rounds to isolate inhibitory Nbs against the target protease (i.e., TEVp).

Selection and Isolation of Inhibitory Nbs Against TEVp via FACS

Using the established controls, an initial Inhibition Sort Gate was drawn around the positive control (LLE-MET-TEVsubstrate-FEH) **(Supplemental Table 1)** to simulate the desired phenotypic outcome of this screen (complete inhibition of TEVp) **(Figure 2C, top left plot)**. The stained NbLibrary-NE-TEVp cells (10^8^ cells) were screened against the controls, and the population (~0.09%) that fell within the Inhibition Sort Gate were isolated and enriched in selection media (YNB CAA 2% GLU, 5 mL, 30°C, 250 RPM, 12-15 hours). This initial enrichment population (ENR1) was prepared for a second round of FACS as previously described, with the amendment that now 2 million cells were required. ENR1 was screened against the controls, and the Inhibition Gate was used to isolate a population (~3%) that was collected and enriched in selection media (YNB CAA 2% GLU, 5 mL, 30°C, 250 RPM, 12-15 hours). The second and third enrichments (ENR2 and ENR3) were prepared as mentioned and assayed on the BDFACS Melody, where a population expressing the complete inhibition phenotype desired (high fluorescent signals of anti-FLAG PE and anti-HA Alexa 647). The top 10% of this final population was isolated through a fourth FACS (ENR4) round and enriched in selection media (YNB CAA 2% GLU, 5 mL, 30°C, 250 RPM, 12-15 hours). Dot plots depicting each round of FACS are represented in **Figure 2**.

Cytometric Assay of Single Nb Constructs from Sorted NbLibrary-NE-TEVp

Dilutions of ENR4 were plated using **Equation 1** to plate single clones from the sorted NbLibrary-NE-TEVp (SC-NbLibrary-NE-TEVp) at cell densities of 200 and 500 cells/plate.

(1)	$\frac{\# of Cells Collected}{Volume of Collected Cells (uL)}=\frac{cells}{uL}$

These dilutions were plated on selection media (YNB CAA 2% GLU) and incubated at 30°C for 24-48 hours. Colonies (n=24) containing one Nb plasmid were picked, inoculated in 2 mL selection media, and inoculated at 30°C (250 RPM) for 24 hours. As previously mentioned, these cultures were prepared for cytometric assay (2 million cells). Just as with the FACS isolation, positive (LLE-MET-TEVsubstrate-FEH) and negative (LLE-pY3-LYS-TEVp-FF) controls were used to allow for the quantification of the inhibitory effect of the Nbs (TEVp-Nb 1-24) on TEVp **(Figure 2D)**. As previously reported, the ratio of anti-FLAG/anti-HA average fluorescent signals can be quantified to create a fold-change measurement of protease inhibition related to substrate cleavage ^3^. The negative control (LLE-pY3-LYS-TEVp-FF) is the baseline measurement, with its anti-FLAG/anti-HA average fluorescent ratio normalized to one. Fluorescent ratios from each Nb were calculated and normalized to the baseline anti-FLAG/anti-HA measurement. With this, the degree of inhibition of TEVp equating to the absolute value of the difference between the normalized ratio of the Nb construct and the baseline control, the greater this difference correlating to the strongest inhibitor. Example calculations of TEVp-Nb4 and TEVp-Nb9 degrees of inhibitions, as well as the baseline TEVp activity ratio, are outlined in **Equations 2-4**.

(2) Active TEVp (baseline activity ratio)	$\frac{Mean anti-FLAG Signal of Active TEVp}{Mean anti-HA Signal of Active TEVp}=\frac{35065}{1076}=32.6$

(3) TEVp-Nb4 Inhibition Ratio (normalized)	$\frac{\left( \frac{Mean anti-FLAG Signal of Nb4-NE}{Mean anti-HA Signal of Nb4-NE} \right)}{\left( \frac{Mean anti-FLAG Signal of Active TEVp}{Mean anti-HA Signal of Active TEVp} \right)}=\frac{\frac{11614}{7351}}{32.6}=0.05$

(4) TEVp-Nb9 Inhibition Ratio (normalize)	$\frac{\left( \frac{Mean anti-FLAG Signal of Nb4-NE}{Mean anti-HA Signal of Nb4-NE} \right)}{\left( \frac{Mean anti-FLAG Signal of Active TEVp}{Mean anti-HA Signal of Active TEVp} \right)}=\frac{\frac{25416}{7801}}{32.6}=0.1$

*Plasmid Purification and Sequencing of Single TEVp-Nb Constructs*

TEVp-Nb purification plasmids (TEVp-Nb4, TEVp-Nb5, TEVp-Nb7, TEVp-Nb9) were isolated from yeast colonies (1 colony, 1 mL YNB CAA 2% GLU, 30°C, 250 RPM, 12-15 hours) using the QIAGEN Spin Miniprep Kit (#27106, user-established protocol: Isolation of plasmid DNA from yeast PR04.doc Oct-01). The yeast-prepped (YP) plasmids were transformed into DH5α cells (NEB #C2987H), plated on selection media (LB AMP, 100 mg/mL), and grown at 37°C for 12-16 hours. Once colonies formed, single colonies were picked and inoculated in 5 mL selection media and grown at 37°C for 12-16 hours (250 RPM). Plasmids were purified using the QIAGEN Spin Miniprep Kit (#27106) and sent for Sanger Sequencing (Genewiz, Azenta Life Sciences). Primers **(Supplemental Table 9)** were designed to cover the entire Nb (~500 bp) to determine the amino acid composition of the whole Nb sequence **(Table 1)** while also adding *BsaI* restriction enzyme cut sites to each terminus.

*Purifying TEVp-Nb Inhibitors*

pET26b_Nb.b201 (a gift from Andrew Kruse (Addgene plasmid #131404; http://n2t.net/addgene:131404 ; RRID:Addgene_131404)^2^, was modified to remove Nb.b201, replacing it with a linker region containing two *BsaI* restriction enzyme cut sites, pET26b-BsaI **(Supplemental Table 1)**, was used as the purification vector. Using a *BsaI* Golden Gate Assembly, each Nb (TEVp-Nb4, TEVp-Nb5, TEVp-Nb7, TEVp-Nb9) was cloned into pET26b-BsaI, transformed (unpurified) into DH5α cells (NEB #C2987H), plated on selection media (LB KAN, 50 mg/mL), and grown at 37°C for 12-16 hours. Once colonies formed, single colonies were picked and inoculated in 5 mL selection media and grown at 37°C for 12-16 hours (250 RPM). Plasmids were purified using the QIAGEN Spin Miniprep Kit (#27106) and sent for full plasmid sequencing (Plasmidsaurus, standard plasmid) to confirm correct assembly.

Following a modified protocol from *McMahon et al.*^2^, pET26_Nb **(Supplemental Table 1)** plasmids were transformed into BL21 (DE3) *E. coli* cells (NEB #C2527) in preparation for periplasmic protein purification. Purification was conducted via centrifugation of gravity columns containing high-density nickel agarose beads (Goldbio #H-350-100) with 100 uL of unpurified lysate and all wash flowthroughs collected for post-purification SDS-PAGE. Purified proteins were eluded in a solution containing 400 mM imidazole and dialyzed in 1 L high-salt buffer for 15 hours (10 kDa MWC SnakeSkin^TM^ Dialysis Tubing, ThermoFisher Scientific #68100). After dialysis, purified Nbs were concentrated using Amicon^R^ Ultra-15 Centrifugal Filters (10 kDa MWC, Millipore Sigma #UFC901024), and concentration was taken using the NanoDrop ONE (ThermoFisher Scientific #ND-ONE-W). Each purified Nb was run on SDS-PAGE for purity confirmation **(Supplemental Figure 18)** and stored at -80°C in a 15% glycerol solution.

*TEVp Purification*

pRK793, a gift from David Waugh (Addgene plasmid #8827; hhtp://n2t.net/addgene:8827; RRID: Addgene_8827) ^6^ was transformed into BL21 DE3 Codon Plus RIL *E. coli* (Agilent #230245). Using the Ni-NTA FPLC purification protocol similar to that outlined in *Kapust et al*. ^6^, a starting culture (1L inoculation volume) was induced to purify the His-tagged TEVp. The purified enzyme was aliquoted and flash frozen, with size and purity confirmation conducted using SDS-PAGE **(Supplemental Figure 19)**. The activity of TEVp *in vitro* was confirmed using a FRET kinetic assay of varying concentrations of the purified enzyme compared to a purchased enzyme (NEB TEVp, #P8112S), canonical FRET TEVp substrate (ABZ-SENLYFQSG-Lys(DNP)) synthesized by BIoresiliance® (MilliporeSigma) held constant at 20 uM **(Supplemental Figure 20)**.

*Characterization of Isolated Inhibitors*

Deep-Sequencing Analysis of NGS Results

A Jupyter notebook based on Python was developed on HiperGator, the high-performance supercomputer at the University of Florida, to analyze the NGS results. Reverse and forward frames were converted from FASTQ to FASTA format and imported into the Pandas data frame after removing the sequencing ID. An alignment based on a 12-base mutual region was then performed between the forward reads and the reverse complement of the reverse reads to obtain the full sequence. Next, each round of data was separated using a 3-base unique identifier at the start and end of each NB sequence. The data were translated into amino acid sequences using the Biopython library. After data cleaning, which involved removing unusually short and long reads, the CDRs were extracted from each sequence using the three amino acids flanking each side. The number of occurrences of each unique NB sequence and the frequency of each amino acid in each position were calculated for enrichment analysis. Based on 24 inhibitory NB and their corresponding round 4 CDR3 ranks, a dataset of inhibitory (round 4 repeats higher than 40) and non-inhibitory (Repeats in round 4 / repeats in round 2 <1) was created. CDR3 sequences numerical representation created using the pre-trained Facebook ESM-2B 3B parameters language model. To visualize CDR3 sequence similarity, embeddings were reduced to two dimensions using the UMAP-supervised algorithm. The entire NGS data processing, alignment, translation, cleaning, and visualization pipeline has been backed up to GitHub for future use (https://github.com/nimaajayebi/HARP).

FRET-based Assays

For all FRET-based assays, a running buffer (20 mM Hepes (pH 6.5), 120 mM NaCl, 0.4 mM EDTA, 20% glycerol, and 1 mM DTT) was used to prepare the samples.

**TEVp Activity Assays**

TEVp was assayed in triplicate across a range of concentration gradients (0-50 uM) of the Abz-DNP substrate (Biomatik, Custom Peptide Synthesis). For each experiment, triplicates of 8 50-uL TEVp samples were prepared in the first three columns of a matte black flat bottom 96-well plate (Greiner Bio-One, #655076) at a fixed concentration of 0.4 uM (for a reaction concentration of 0.2 uM in 100 uL). The substrate was prepared in the fifth column, at a volume of 150 uL/well, with each well containing a solution of Abz-DNP substrate at a concentration 2 times that of the reaction concentration. A multichannel pipette was used to add 50 uL of the substrate to each enzyme sample. Fluorescence intensity (320/405 nm) was measured on a BioTek Synergy Neo2 plate reader at 1 read/20 seconds for 30 minutes. Once all assays were completed, the data was exported into GraphPad Prism (10.1.1) for analysis. Slopes were calculated from the linear regions of each time (x-axis) vs. RFU (y-axis) to get the average reaction velocity, which was subsequently plotted against substrate concentration (on the x-axis) **(Supplemental Figure 21)**. Average *K_M_* and *v_max_* were extrapolated using a nonlinear regression Michaelis-Menten curve fit, with the error based on the standard deviation of each *K_M_* calculated for each set of substrate concentrations.

**Nb-mediated Inhibition of TEVp Assays**

TEVp was assayed in triplicate across a concentration gradient of Nb for each inhibition assay, while the enzyme and substrate concentrations were held constant at 0.2 and 20 uM, respectively. For each experiment, triplicates of 8 50-uL TEVp samples were prepared in the first three columns of a matte black flat bottom 96-well plate (Greiner Bio-One, #655076) at a fixed concentration of 0.4 uM (for a reaction concentration of 0.2 uM in 100 uL). The Nb being assayed in the experiment (one Nb was tested at a time) was prepared in the fourth column of the plate using a 2-fold dilution down the column, each well with a final volume of 100 uL. Using a multichannel pipette, 25 uL of each Nb dilution was added to the TEVp samples, creating triplicates of each enzyme-Nb sample (the final Nb concentration for a 100 uL reaction is ¼ the dilution concentration). The enzyme-Nb samples were incubated in the dark for 30 minutes at RT. The substrate was prepared in the fifth column at a volume of 100 uL/well, with each well containing a solution of Abz-DNP substrate at a concentration of 80 uM, 4 times that of the reaction concentration. Once the incubation period was completed, a multichannel pipette was used to add 25 uL of the substrate to each enzyme-Nb sample. Fluorescence intensity (320/405 nm) was measured on a BioTek Synergy Neo2 plate reader at 1 read/20 seconds for 30 minutes. Once all assays were completed, the data was exported into GraphPad Prism (10.1.1) for analysis. Slopes were calculated from the linear regions of each time (x-axis) vs. RFU (y-axis) to get the average reaction velocity, which was then normalized to the slope of a TEVp-substrate sample (no Nb) at the reaction concentrations. The normalized slopes were converted to a percentage, with the TEVp-substrate sample being 100% active, and the activity percentages were plotted against Nb concentration (on the x-axis) **(Supplemental Figure 10)**. Average *IC50* values for each Nb were extrapolated using a nonlinear regression [inhibitor] v. response variable slope (four parameters) curve fit, with the error based on the standard deviation of each *IC50* calculated for each sample within the triplicate. Inhibition constants, K_I_, for each Nb, were computed using **Equation 5**, and the error was subsequently calculated for each Nb using **Equation 6**.

(5)	$K_{I, avg}= \frac{{IC50}_{avg}}{1+\frac{[S]}{K_{M,avg}}}$

(6)	$K_{I,error}=\frac{1}{K_{I, avg}}\sqrt{(}$ ${{(IC50}_{error})}^{2}+{{(K}_{M,error})}^{2})$

This assay was conducted as described for all FRET-based inhibition assays.

Biolayer Interferometry (BLI) Binding Assays

For each BLI assay, PBST (0.05% Tween) was used as the neutral/running buffer and Regeneration Buffer (No Salt) (GatorBio, #120063) was used as the regeneration buffer. A single set of probes was used for all the BLI experiments to maintain consistent probe batches **(Supplemental Figure 22)**. To confirm the BLI assay design, a previously published Nb, Nb.b201 was tested against human albumin serum (HSA). The resulting *K_D_* (277 nM) fell within range with the published value (420 nM)^2^. We conducted an additional control assay in which we had a Nb sample TEVp-Nb5 with and without the presence of trace amounts of glycerol, with the resulting *K_D_* values being nearly identical **(Supplemental Table 8)** while additionally falling in the same order of magnitude as the reported value **(Supplemental Table 4)**. We attribute the slight discrepancies to variability within the lab environment (i.e., temperature, humidity) and the potential of batch-to-batch variability with Nb samples (as many purifications were conducted during manuscript completion) and running buffer preparation.

The binding affinity of each Nb-NE to TEVp was determined via a standard BLI Kinetic (K) Assay conducted on the GatorBio GatorPlus BLI. Using anti-VHH probes (GatorBio, #160032), each assay began with saturating the probes with Nb at a constant concentration of 34.4 nM for two minutes between the probes were introduced to varying concentrations of TEVp for 300-second association and dissociation steps. Curve binding of the association and disassociation curves were calculated to an R^2^>0.9 to obtain the *k_on_* and *k_off_* values, and thus *K_D_* values for each Nb, with the errors (σ) calculated using **Equation 7**.

(7)	$\sigma_{avg}=K_{D, avg}\times\left( \sqrt{\left( \frac{\sigma_{kon,avg}}{k_{on, avg}} \right)^{2}+\left( \frac{\sigma_{koff,avg}}{k_{off, avg}} \right)^{2}} \right)$

This assay was conducted as described for all BLI binding assays.

Supplemental References

1.	Demeestere, D. et al. Development and Validation of a Small Single-domain Antibody That Effectively Inhibits Matrix Metalloproteinase 8. *Mol Ther* **24**, 890-902 (2016).

2.	McMahon, C. et al. Yeast surface display platform for rapid discovery of conformationally selective nanobodies. *Nature Structural & Molecular Biology* **25**, 289-296 (2018).

3.	Martinusen, S.G. et al. Modular and integrative activity reporters enhance biochemical studies in the yeast ER. *Protein Engineering, Design and Selection* **37** (2024).

4.	Gietz, R.D. & Schiestl, R.H. High-efficiency yeast transformation using the LiAc/SS carrier DNA/PEG method. *Nat Protoc* **2**, 31-34 (2007).

5.	Loock, M. et al. High-Efficiency Transformation and Expression of Genomic Libraries in Yeast. *Methods Protoc* **6** (2023).

6.	Kapust, R.B. et al. Tobacco etch virus protease: mechanism of autolysis and rational design of stable mutants with wild-type catalytic proficiency. *Protein Eng* **14**, 993-1000 (2001).
